# Maternal transfer and long-term population effects of PCBs in Baltic grey seals using a new toxicokinetic-toxicodynamic population model

**DOI:** 10.1101/2022.02.01.478642

**Authors:** Karl Mauritsson, Jean-Pierre Desforges, Karin C. Harding

**Affiliations:** Department of Biology & Bioinformatics, University of Skövde, Skövde, Sweden; Department of Environmental Studies and Sciences, University of Winnipeg, Winnipeg MB, Canada; Department of Biological & Environmental Sciences, University of Gothenburg, Gothenburg, Sweden

**Author notes:** Correspondence: Karl Mauritsson.

**Keywords:** PCBs, Population model, TKTD, maternal transfer

## Abstract

Empirical evidence has shown that historical exposure of polychlorinated biphenyls (PCBs) to Baltic grey seals not only severely affected individual fitness, but also population growth rates and most likely caused the retarded recovery rate of the depleted population for decades. We construct a new model which we term a toxicokinetic-toxicodynamic (TKTD) population model to quantify these effects. The toxicokinetic sub model describes in detail the bioaccumulation, elimination, and vertical transfer from mother to offspring of PCBs, and was linked to a toxicodynamic model for estimation of PCB-related damage, hazard, and stress impacts on fertility and survival rates. Both sub-models were then linked to a Leslie matrix population model to calculate changes in population growth rate and age structure given different rates of PCB exposure. Toxicodynamic model parameters related to reproductive organ lesions were calibrated using published historical data on observed pregnancy rates in Baltic grey seal females. Our model showed that increased PCB exposure caused reduced fertility, decreased vertical transfer, and increased biomagnification. Compared to empirical data, the TKTD population model described well the age-specific bioaccumulation pattern of PCBs in Baltic grey seals, and thus, the toxicokinetic parameters, deduced from literature, are believed to be reliable. The model also captured well the general effects of PCBs on historical population growth rates. The developed model can be used to perform population viability analyses of Baltic grey seals with multiple stressors, also including by-catches and different hunting regimes. The model can also be extended to other marine mammals and other contaminants than PCB by adjustments of model parameter values and thus provides a test bed in silico for new substances.

## Introduction

During the past century, human activities have dramatically affected the ecological balance in the Baltic Sea. Recent assessments of biodiversity report inadequate status in all levels of the food web including indications of decreased nutritional status in fish and mammals (Kauhala et al. 2014, Hansson et al. 2017, HELCOM 2018b, Sonne et al. 2020). The semi-enclosed Baltic Sea is surrounded by nine highly industrialized countries and acts as a sink for municipal and industrial discharges and land run-off. Hazardous substances have been identified as one of the seven distinct threats to the Baltic Sea (HELCOM 2018b). The most harmful substances are those that are persistent, toxic and accumulate in organisms. These include among others persistent organic pollutants (POPs), and one of the POP classes of major concern is polychlorinated biphenyls (PCBs). Based on measurements of sediment accumulation, the PCB deposition in the Baltic Sea gradually increased during the 1940–1960s, reached peak values in the 1970s, and substantially decreased during the late 1970s and 1980s and continue to decline (Eckhéll et al. 2000). Despite the decline, concentrations of PCB in fish are still too high to be acceptable as food for humans according to the Swedish Food Administration (Bjurlid et al. 2018). The issue is even worse for marine mammals that can accumulate even higher levels of PCBs and other persistent contaminants because they are long-lived top predators with large depots of blubber that can store fat-soluble chemicals over decades and recycle them across generations via vertical transfer to offspring during gestation and lactation (Bignert et al. 1998, Klanjscek et al. 2007, Desforges et al. 2012).

Four species of marine mammals occur in the Baltic Sea, whereof the grey seal (*Halichoerus grypus*) is the most abundant. The Baltic grey seal population approached 100 000 individuals in the early 1900s, but bounty hunting caused a decline to about 20 000 animals in the 1940s. When hunting ceased in 1974, the population did not increase as expected, and in the 1970s, the size of the Baltic grey seal population was estimated to only 3000 animals (Harding and Härkönen 1999). A likely explanation is a number of pathological changes, including reproductive failure, termed the Baltic Seal Disease Complex (BSDC) related to PCB exposure (Bergman and Olsson 1986, Bäcklin et al. 2003, Bredhult et al. 2008). Besides PCBs, POPs found in Baltic seals also include dioxins and furans, polychlorinated diphenyl ethers, toxaphenes and chlordanes, among many others (Nyman et al. 2002). PCB and DDT are considered as the most important factors behind BSDC, but they are probably not the only significant ones (Bergman 2007). Despite lower PCB levels in grey seals and their prey today than in the past, population health assessments have found that indicators of reproductive and nutritional status remain below identified threshold values, likely due to combined effects from multiple stressors like pollutants, climate change, and reduced food availability (Bäcklin et al. 2011, Kauhala et al. 2012, Kauhala et al. 2014, HELCOM 2016, Silva et al. 2020, Sonne et al. 2020). Along with decreasing PCB levels during the last decades, the Baltic grey seal population has increased. The population size in 2020 was estimated to about 50 000 animals (HaV and SMHI 2022), but still the population growth rate is suppressed compared to healthy populations. The growing population has intensified conflicts between seals and commercial fisheries by resource competition and damages to catch and fishing gear (Tverin et al. 2019), and in 2001 hunting quotas have been increased up to 3000 seals (Bäcklin et al. 2011), without any risk assessment or population viability analysis, however the hunt has not reached the annual quotas yet. To better understand long term population effects from POPs and energetic stress, better models are urgently required.

Although the most conspicuous adverse effects of PCBs on Baltic grey seals, such as uterine occlusions, have declined during recent decades, PCBs still affect the population through indirect effects such as impaired immunity which in turn influence risk of infection by pathogens and parasites (Bäcklin et al. 2011). According to HELCOM (2018b), the current main pressures to Baltic grey seals are hunting, by-catches in fishery, contaminants and climate change. More than 2000 grey seals are caught incidentally each year in the Baltic fisheries (Vanhatalo et al. 2014). Long-term climate change is predicted to cause shorter and warmer winters with decreasing sea ice coverage (Sundqvist et al. 2012). This will probably reduce the breeding success of Baltic grey seals, since pups born on land have a significantly higher mortality (Jüssi et al. 2008). It is of uttermost importance to develop risk assessment models that consider the sum of many smaller stressors and adverse effects on fitness in order to avoid another wave of overexploitation (Cervin et al. 2020, Silva et al. 2021).

The growth rate of a population is governed by its age-specific fecundity and survival rates. The long-term maximum growth rate of Baltic grey seals is limited by several factors. Females have at most one pup each year, of which 48 % are female (HELCOM 2016). The first parturition occurs on average at an age of 5.5 years and not all females bear a pup each year, especially not young females. Some ovulating females do not complete pregnancy. Further, mortality of adults limits the population growth and senescence and pathological changes in reproductive organs decrease pregnancy rates. Environmental stress, such as parasites, pollution and limitations in food or breeding sites, reduce average fecundity and survival rates. Healthy, undisturbed populations of grey seals have an expected growth rate of 10 % each year (Harding et al. 2007), but observed growth rates are significantly lower.

In order to better understand the temporal dynamics of persistent and toxic chemicals like PCBs in marine long-lived predators like the grey seal it is necessary to develop appropriate modeling approaches. Models need to capture fine and long scale dynamics that apply over an animal’s life span. Since PCBs accumulate in a seal’s body during its whole life span, some health effects may arise only after many years. For instance, PCBs and other lipophilic POPs may have limited adverse effects on the health of an individual as long as they are bound in the blubber of the animal, but when blubber energy is mobilized to fuel various physiological processes during periods of food stress, pollutants are released to the blood and can circulate to sensitive tissues and organs and increase the risk of effects on fertility and survival (Lydersen et al. 2002, Klanjscek et al. 2007). Further, grey seals have some capacity to metabolise PCBs, using xenobiotic metabolising enzymes such as CYP1A (Nyman 2000). Studies of contaminant effects in marine mammals have focused largely on the molecular and individual levels, though some analyses on the population level exist (Pavlova et al. 2016, Hall et al. 2018, Silva et al. 2020). A reason for the limited number of population analyses is probably lack of long-term monitoring studies and data on contaminant levels and associated health effects. To characterize and predict contaminant levels and effects in populations of marine mammals over time, and thus to assess the effectiveness of past, current, or planned management and mitigation efforts, it is necessary to develop a mechanistic understanding of the processes of bioaccumulation (at different temporal scales) and inter-generational transfer as well as to understand how contaminants impose hazard or stress to individuals and how this relates to population dynamics.

Here a population model is developed and applied to predict the temporal dynamics of PCBs and its associated influence on vital rates, population growth, and extinction risk of Baltic grey seals over the past century. The model links dietary PCB exposure to internal concentrations, characterizes vertical transfer from mother to offspring, and relates dynamic internal concentrations to adverse effects on fertility and survival, then integrates this toxicokinetic and toxicodynamic (TKTD) model into an age-structured Leslie matrix model of temporal population dynamics. We focus on PCBs as a model contaminant because of the availability of empirical data, including time trends for PCB levels in Baltic fish and seals as well as frequencies of PCB-related reproductive defects in Baltic grey females. Despite the focus on Baltic grey seals, the model framework is broadly applicable to other marine mammals and bioaccumulative contaminants through informed changes to model parameters. Thereby, the model can be a useful tool in future viability assessments targeting bioaccumulating contaminants in populations of marine mammals.

## Methods

### Model Overview

Our matrix model, called the TKTD population model, for analysing adverse effects of PCBs on Baltic grey seals was developed to link toxicokinetics (TK) of dietary and vertically transferred PCB accumulation to toxicodynamics (TD) of adverse effects on reproduction and survival to ultimately predict changes to population demographics and growth rates (Fig. 1a). The TKTD model is used to modify survival and fecundity parameters of an age-structured Leslie matrix model for an ideal Baltic grey seal population due to internal PCB concentrations following DEBtox and threshold damage model (TDM) principles of damage, hazard, stress, and recovery. While the model runs on an annual basis for population dynamics, each year is divided into three periods (*lactation, delay* and *gestation*), representing different life history phases for a reproducing grey seal female. The year starts with the *lactation period* (lasting 18 days), during which pups are nursed by their mothers. After weaning, females mate and the *delay period* starts (lasting 100 days), after which the fertilized egg is implanted. During the *gestation period* (lasting 247 days), the fetus develops in the uterus (SI-Table 2). The year ends with the birth of new pups. Since all deaths are assumed to occur just before the birth events, all pups that are born have mothers that nurse them. Different TD effects of PCBs occur after specific time periods (e.g., fertility effects estimated after the gestation period).

**Fig. 1.**
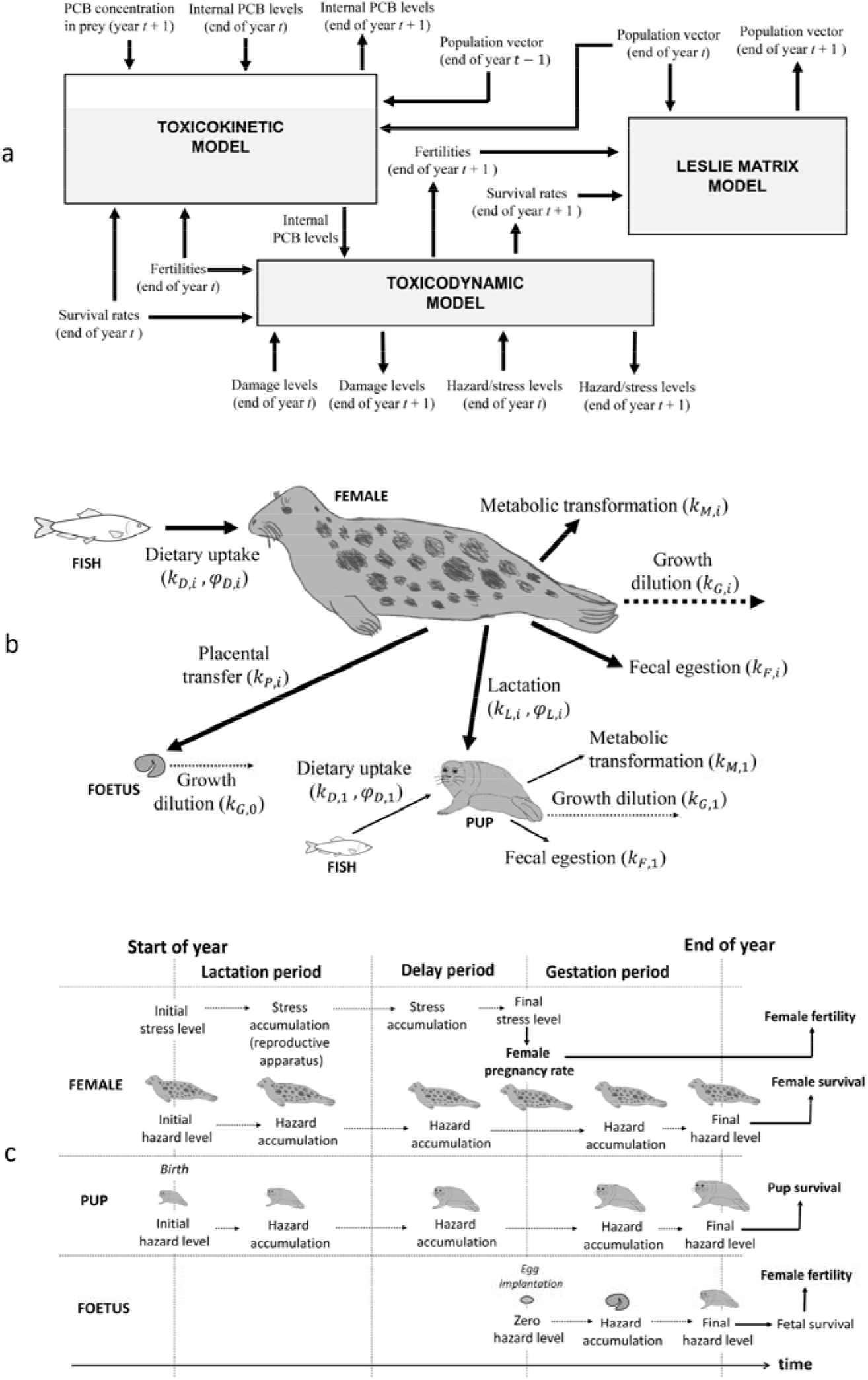
a) Overview of the TKTD population model. Text boxes represent sub-models, performing calculations for year *t* + 1. Free texts are inputs and outputs, with arrows showing relations to sub-models. b) Bioaccumulation model showing the major routes for uptake, transfer and elimination of PCB in grey seals, including associated rate constants (*k_D,i_*, *k_M,i_*, *k_F,i_*, *k_L,i_*, *k_P,i_*, *k_G,i_*) and assimilation efficiencies (*φ_D,i_, φ_L,i_*). c) Effects of PCB on survival and fertility considered in the toxicodynamic model as consequences of hazard/stress accumulation in females, pups and fetuses during a year.

The model is used to perform stepwise annual calculations of PCB concentrations, which are then converted to adverse effects through calculation of damage, hazard, and stress levels (see below for details), which ultimately determine fertilities, survival rates, and number of individuals in different age classes. The TKTD population model consists of the following three sub-models (Fig. 1a): 1) Toxicokinetic model: bioaccumulation model for dietary uptake, elimination and vertical transfer through placenta (during gestation) and breast milk (during lactation); 2) Toxicodynamic model: TDM for damage and recovery as response to internal toxicant levels, obtained from the toxicokinetic model; cumulative damage causes hazard/stress to fetuses, pups, and adults, resulting in reduced fertilities and survival rates; and 3) Leslie matrix model: age-structured matrix model for population growth, with baseline ideal fertility and survival rates adjusted according to the toxicodynamic model of PCB effects.

A brief description of the model framework is provided here, and a complete detailed description and equation formulation is provided in the Supplemental Information (SI). Environmental variables and initial conditions are first established. These include initial seal PCB concentrations, initial damage levels, initial hazard/stress levels, the initial population vector, and annual prey PCB concentrations (SI-Table 1, SI-Fig. 2). The initial population *n*(O) is calculated from the stable age distribution for a population with ideal fertilities and survival rates (SI-Table 4), based on an initial female population size *N_tot_*(0) of 8820 individuals. New state variables are calculated for one year at a time, based on values of state variables the previous years. For each year, the following calculations are performed, and the procedure is repeated until the final year: 1) calculation of PCB concentration in fetuses, pups and females as a result of bioaccumulation and vertical transfer from females to offspring; 2) calculation of cumulated damage in fetuses, pups and females, based on internal PCB concentration; 3) calculation of cumulated hazard/stress in fetuses, pups and females, based on internal PCB concentration and cumulated damage; 4) calculation of reduced fertilities, based on fetal hazard and female stress (accounting for damage to the reproductive apparatus); 5) calculation of reduced survival rates, based on cumulated hazard in pups and females; and 6) calculation of new population size and structure, based on reduced fertilities, reduced survival rates, and previous year’s population data. It is assumed that all deaths and births occur at the end of the year, though hazard and stress accumulated during the year.

The TKTD population model was implemented as a set of functions and script files in the numerical software MATLAB^®^ (Mathworks Inc., Natick, MA, USA) and all analyses were performed with these.

### Toxicokinetic Model

The toxicokinetic model describes PCB bioaccumulation in offspring and females through dietary uptake and elimination (metabolic transformation and fecal egestion) as well as vertical transfer from mother to embryo through the placenta during gestation and from mother to pup through breast milk during lactation (Fig. 1b). During lactation, neither the mother nor the pup consumes prey, thus the primary exchange of PCBs occurs through vertical transfer via the milk (plus environmental losses). During the *delay period*, both females and pups feed on prey. During the *gestation period*, females and pups continue to feed, but females also transfer PCB to fetuses through the placenta.

#### Growth model

The change of PCB concentrations as individuals grow is termed growth dilution. This is captured in the TK model through a generalized sub model describing animal growth. Fetuses and pups are assumed to grow exponentially during the gestation and lactation periods, after weaning, grey seal pups typically lose weight during some months due to reduced nutritional intake, but when they become more skilled hunters they start to regain weight (Kauhala et al. 2017). For simplicity, it is here assumed that pups keep the weaning weight until they reach sub adult age (1 year). Grey seal females typically reach asymptotic body size at age six, about the time when they become sexually mature. When they nurse pups, they lose a considerable amount of weight, but this is regained when they feed during the remaining part of the year. Our model assumes a von Bertalanffy growth function to describe body growth of grey seal females from weaning to sexual maturity. Once females reach mature body length, they start to breed. We assume that growth dilution rates are approximately constant within each period of a year (delay, gestation, lactation) for each seal life stage (fetuses, pups, adults).

Since primiparous females (females which give birth for the first time) are typically smaller than multiparous females (Lang and Iverson 2009), the mature body weight *W_mat_* is assumed to be somewhat lower than the maximum body weight *W_max_* reached by multiparous females just before they start the lactation period. Since females lose significant weight during lactation, it is assumed that their weight decreases exponentially throughout the period, from the initial body weight (*W_mat_* or *W_max_*) to a final lactation body weight (*W*_*lac*,6_ or *W*_*lac*,7_, for primiparous and multiparous females, respectively). During the delay period, mature females are assumed to linearly regain the weight they lost during lactation. During the succeeding gestation period, they continue to grow according to von Bertalanffy and reach the maximum body weight *W_max_* at the end of the year. The succeeding years, females are assumed to repeat the pattern of exponential decline (lactation period), linear regain (delay period) and von Bertalanffy growth (gestation period) (SI-Fig. 6). The lactation body weight *W*_*lac*,7_ for multiparous females is specified from data, whereas the corresponding body weight for primiparous females is calculated from the assumption *W_lac,6_/W_mat_* = *W_lac,7_/W_max_*. Values of parameters used to describe body growth were chosen based on published data for Baltic grey seals (SI-Table 2).

#### Bioaccumulation model

Bioaccumulation in females (age 2 to 46) includes dietary uptake, elimination (metabolic transformation and fecal egestion) and vertical transfer to offspring (Fig. 1b). The toxicokinetics is described by a physiologically based equation similar to the one-compartment model commonly used in DEBtox models. The rate of change of the total amount of PCBs in a female is expressed as the rate by which PCBs are assimilated (assumed to be proportional to the concentration in the prey) subtracted by the rate by which PCBs are eliminated (assumed to be proportional to the concentration in the body). Thus, the concentration of PCBs in females of age-class *i* during time-period *j* 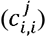 can be approximated using the following first-order ordinary differential equation:

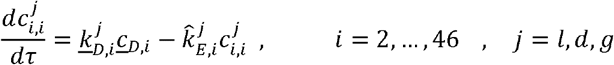

where 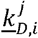 is the mean PCB dietary uptake rate of seals of different age-classes and time-periods, 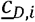 is the mean dietary PCB concentration, and 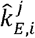 is the total PCB elimination rate. Dietary uptake rates are a function of the prey consumption rate (kg/yr) and the diet assimilation efficiency. We follow the general principles of DEB theory such that food uptake is proportional to the surface area of the structural body volume and a fixed prey consumption factor based on empirically-derived maximum yearly food intake. We assume that mean PCB concentration in prey is constant throughout the year. The diet includes fish of different species and each seal age-class consumes a diet of specific prey composition. According to Hansson et al. (2017), 80 % of the Baltic fish biomass is constituted by sprat (*Sprattus sprattus*), herring (*Clupea harengus*) and cod (*Gadus morhua*). It was assumed that Baltic grey seals feed exclusively on these three species. Prey preference indices, describing the preference of prey items by age class, were calculated from published mean fractions of total prey biomass found in analyses of gut content in Baltic grey seals (Lundström et al. 2010, Tverin et al. 2019) (SI-Table 3).

Dietary PCBs are thus a function of the mean weight fraction of each prey item and their respective PCB levels. Elimination of PCBs is accounted for through fixed rate constants for fecal egestion, metabolic transformation, placental transfer, and lactation transfer. Since fecal egestion is accounted for via assimilation efficiencies for dietary uptake and lactation, the fecal egestion rate constants are neglected. All rate constants are assumed to be age-specific, though some of them have similar or same values for all age classes. We use estimated metabolic transformation rate constants from Hickie et al. (2005) for total PCBs in ringed seals after scaling for differences in body size between species and age-classes assuming transformation rate constants scale according to Kleiber’s law for metabolic rates (Kleiber 1962) (SI-Table 6).

Fetuses assimilate PCBs only through vertical transfer via the placenta (Fig. 1b). It is assumed that all PCBs that are eliminated by a mother through placental transfer are assimilated by her fetus, which has no capacity to eliminate them. The concentration of PCB in the fetus 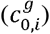 is therefore a function of the PCB concentration in the mother of age-class *i* during the gestation period (*j* = *g*) 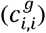, body weight ratio between mother and fetus (*ω*_0,*i*_), the placental transfer rate constant (*k_p,i_*), and fetal growth dilution (*k*_*G*,0_) according to the following equation:

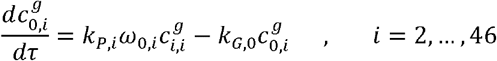

The accumulation of PCBs in pups (*i* = 1) is calculated similar to that of older age classes, but with no placental transfer and a positive contribution from vertical transfer through lactation (Fig. 1b). In other words, the PCB concentration in pups 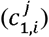 is a function of maternal PCB concentrations 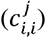, the mean lactation PCB transfer rate 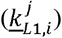, the mean annual PCB dietary uptake rate of pups 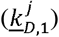, the mean dietary PCB concentration 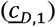, and the total PCB elimination rate of the pup 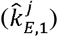 according to the following equation:

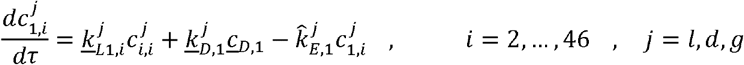

#### Conversion of PCB concentrations

The TK model estimates total body concentrations of PCBs for seals and fish, while empirical data is reported on a lipid weight basis. We thus convert calculated whole-body PCBs to lipid concentrations assuming all PCBs in the body of a seal is bound in fat tissue. Thus, the concentration in lipids is calculated as the concentration in the body divided by the body fat index (total body weight/total lipid weight) for each seal age-class (SI-Table 6). As a simplification, it is assumed that mean body fat indices are the same for all age classes at the time of field sampling in autumn when seals are at their fattest (Kauhala et al. 2017). The prey lipid PCB concentration is calculated in the same way as seals.

### Toxicodynamic model

The toxicodynamic model describes how PCB body loads of fetuses, pups, and females cause accumulation of damage, hazard, and stress, and how these are translated into reduced survival and fertility (Fig. 1c, SI-Fig. 8-9). The toxicodynamics are based on the TDM, introduced by Ashauer et al. (2007b). TDM is in turn based on DEBtox theory but adds cumulative effects and capacity to recover from damage. Here, internal contaminants induce *damage*, which accounts for physiological disturbances from all kinds of biochemical and physiological processes involved in toxicity. *Hazard* is then probability to die during a time interval based on the cumulative amount of accrued damage. Recovery is included as a physiological means to repair damages caused by internal contaminants, for instance cell repair mechanisms and physiological adaptations.

The current TD model follows the general principles of DEBtox in assuming that the toxic effects of PCBs on grey seals are considered through the lens of particular physiological modes of action (pMoAs) affecting fundamental physiological processes like somatic maintenance or maturation. While our model is not based on DEB theory in the description of energetics and life history traits, we interpret internal PCB concentration effects using the lense of DEBtox pMoAs with a hazard model that represents damage to reproductive organs and mortality of fetuses, pups and females. In this context, we assume decreased survival can result from increased somatic maintenance costs (e.g., repair of lesions) due to cumulative damage causing hazard in pups, juveniles, and adult females. Similarly, decreased reproduction can result from increased fetal hazard (fetal mortality) and reproductive stress. Assuming a hazard model (Billoir, et al., 2007), we consider *reproductive stress* as an indirect effect on reproduction via increased costs for somatic maintenance due to cumulated damage causing stress to the reproductive apparatus of adults (e.g., reproductive organ lesions). Other indirect effects on reproduction, such as decreased feeding rate and increased growth costs are neglected. Notice that three different kinds of damage are considered: 1) damage to pups and females (causing hazard that decreases survival); 2) damage to reproductive apparatus (causing stress that reduces fertility); and 3) damage to fetuses (causing hazard that kills them and reduces the fertility of their mothers).

Our TD model follows the TDM approach to calculate hazard, stress, and recovery (Ashauer et al. 2007b). The cumulation of all three types of damage listed above are described by the same governing equations, which only differ in values of model parameters (SI-Table 7). According to TDM, the contaminant-induced damage rate increases linearly with the internal contaminant concentration, whereas the recovery rate is proportional to the current damage. Damage at any time point is a function of internal PCB concentration, the killing or stress rate constant, the recovery rate constant, and current level of damage. The hazard or stress at any time is a function of current hazard/stress levels (cumulative), and current damage above an age-specific damage threshold. Since fetuses and pups do not invest energy in reproduction, damage to reproductive organs begins to accumulate when seals reach the age of 2.

Fertilities are affected by PCB exposure in two different ways: reduction due to reproductive stress and reduction due to fetal hazard. It is here assumed that the state of females at the end of the delay period (just before gestation starts) determines fertility reduction due to reproductive stress (e.g., reproductive lesions) and that the state of fetuses at the end of the gestation period determines fertility reduction due to fetal hazard. Fertility impacts are implemented as a reduction factor (hazard/stress) multiplied to baseline age-specific fertility values. Survival rates of pups and females are decreased by cumulated hazards, also implemented as a multiplicative reduction factor to baseline survival values.

The three sets of TD parameters include *reproductive stress* parameters, *fetal hazard* parameters, and hazard parameters related to *survival* of pups and adults. Each parameter set includes three type of constants: *stress/killing rate constants, damage threshold levels*, and *recovery rate constants* (SI-Table 7). With respect to the many empirical observations of reproductive organ lesions in Baltic grey seals during the 1970s and the 1980s, stress to the reproductive apparatus is probably the most crucial path in which PCBs affect vital rates. The reproductive stress parameters were estimated by a calibration procedure where pregnancy rates predicted by the TKTD population model were compared to reported pregnancy rates (Roos et al. 2012). For the other toxicodynamic parameters, values were chosen based on parameters estimated in Desforges et al. (2017) for their model on PCB exposure and effects in captive fed mink. Since the mink model did not account for cumulated damage and recovery, we had to estimate these TDM model parameters using exposure times for the mink dataset as well as several simplifying assumptions described in the SI. Ultimately, estimated mink TDM parameters were converted to grey seal TDM parameters assuming that contaminant sensitivity (no effect concentrations and damage threshold levels) among species scales with specific metabolic rates (Baas and Kooijman 2015) (SI-Table 7). We assumed that tolerance parameters are the same for different mammals exposed to a specified contaminant, and thus killing rate constants also remain unchanged when exposure times are similar.

### Leslie Matrix Model

The Leslie matrix model follows Harding et al. (2007) to implement a full age-structured model for Baltic grey seals using annual time steps and 46 age classes. Since an insufficient number of males is unlikely to restrict growth of a seal population (Harding et al. 2007), only the size of the female population is considered. Individuals of age class *i* = 1 (with an age of 0-1 year) are here referred to as *pups*, whereas individuals of age class *i* = 2,…,46 are referred to as *females*. Ideal vital rates (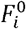 and 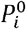) for a population with maximal possible reproduction, derived by Harding et al. (2007) were used as baseline values in our model, representing fertilities and survival rates in absence of PCB exposure (SI-Table 4). These vital rates result in a predicted maximum possible population growth rate of *λ* = 1.10.

Unlike Harding et al. (2007), the Leslie matrix elements in our model are not constants. Fertilities (*F_i_*) and survival rates (*P_i_*) are linked to damage levels that have accumulated as a result of the history of internal PCB concentrations, which in turn depend on the history of PCB concentrations in the prey. It is assumed that all deaths occur at the end of a year and immediately after, all pups are born. Since only females that survive to the end of a year give birth to new pups, the first row of the Leslie matrix contains products of survival rates and fertilities, also a difference from Harding et al. (2007). Since the population size is accounted for just after breeding, the Leslie matrix model may be characterized as a *post-breeding model*.

#### Density dependence and stochasticity

Recent observations of reduced pregnancy rates in Baltic grey seals have been explained as a consequence of the population approaching carrying capacity (Kauhala et al. 2014) and observations of thinner blubber layers have been explained by food limitation (HELCOM 2018a). Fertilities and survival rates of different age classes are thus assumed to be differently affected by population density. We set the carrying capacity to 100 000, since the Baltic grey seal population likely approached this size in the early 1900s (Harding and Härkönen 1999). It was assumed that adult survival and fertility are only dependent on the number of adults, not on the number of pups. Since pups are more sensitive to harsh conditions than older animals, it was also assumed that pup survival is more affected by population density than other age-classes. Density-dependent changes in age-specific survival and fertility were calculated from population size and age-class specific density effect factors (SI-Table 5). The adult density effect factor was parameterized such that the total population (females and males) approached carrying capacity under ideal conditions (no adverse effects from contaminants).

We included a sub model for environmental stochasticity to account for fluctuations in birth and death rates due to natural environmental changes that affect all individuals in the population simultaneously (e.g. Engen et al. 2005). This is included for survival and fertility through normally distributed synchronized fluctuations, calculated using survival/fertility fluctuation factors that capture the relative magnitude of observed stochastic fluctuations in the population (Harding et al. 2007, Silva et al. 2021). These factors are normally distributed random numbers restricted to expected ranges for survival (0-1) and fertility (0-0.5). It was assumed that environmental stochasticity poses small fluctuations in survival rates and five times as large fluctuations in fertilities (SI-Table 5). Because simulations including environmental stochasticity did not provide any meaningful or relevant changes to model results compared to simulations without stochasticity, all results in the current study exclude stochasticity for simplicity. We rather include it here and in detail in the SI for consideration in future studies.

### Model Simulations

#### Steady state simulations

If prey PCB concentrations are constant over time, a stable state will finally be reached where PCB, damage, hazard, and stress levels are stationary. If density dependence is neglected, also fertilities and survival rates stabilize at constant values and the population grows (or declines) at constant rate. Under these conditions, biomagnification factors and stable population growth rate can be defined. Age-specific biomagnification factors (based on lipid concentrations) are calculated as the ultimate concentration ratios in seals and their prey. The stable population growth rate is calculated as the dominant eigenvalue of the Leslie matrix. For steady-state simulations we test the effect of different levels of constant PCB lipid concentrations in prey (0-10 mg/kg) and run the TKTD population model for 100 years. Endpoints of interest were simulated temporal changes in mean PCB lipid concentrations and population size for Baltic grey seals.

#### Sensitivity analysis

To describe the impact of parameter uncertainty on model outputs we run a global sensitivity analysis (change one parameter at a time). A sensitivity analysis investigates how uncertainty in input parameters causes uncertainty in population growth and can be used to identify parameters that are critical for population viability (Lacy et al. 2018). Given that Harding et al. (2007) performed a detailed sensitivity analysis of their grey seal Leslie matrix model for Leslie matrix parameters, we did not perform a similar analysis here. Instead, we focus the sensitivity analysis to TK and TD parameters, assuming steady-state conditions. This analysis was performed for three of the TK model parameters: the lactation rate constants *k_L,i_*, the placental transfer rate constants *K_P,i_* and the metabolic transformation rate constants *K_M,i_*. The lactation rate constant *k*_*L*,7–46_ refers to multiparous females and fixed ratios between lactation rate constants for different age classes were adopted. The placental transfer rate constants *k*_*P*,6–46_ were the same for all fertile age classes. The metabolic transformation rate constant *k_M,i_* were dependent on age class body mass. Sensitivity was also assessed against three TD model parameters across life stages: killing/stress rate constants (*σ*_0_, *σ_i_*, 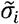), recovery rate constants (*r*_0_, *r_i_*, 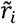), and damage threshold levels (*d*_*T*,0_, *d_T,i_*, 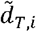). A constant PCB lipid concentration in prey, generating positive stable growth rate over time, was used as input and the toxicokinetic rate constants were varied one at time, whereas other parameters were held constant. Stable population growth rate was plotted as a function of the perturbed relative value of investigated model parameters, defined as the as the perturbed value (*p_pert_*) divided by the value adopted in the model (*p_mod_*):

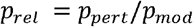

#### Realistic simulations of temporal dietary PCBs

Segerstedt (2019) compiled data from previous studies of PCB concentrations in grey seals and their primary prey items (cod, sprat, and herring) from all regions of the Baltic Sea during 1966-2015, converted all concentrations to a lipid weight basis and clustered data into time intervals of 5 years. We linearly interpolated prey PCBs concentrations over each five-year period to obtain yearly PCB concentrations, used as input to the toxicokinetic model (Fig. 3a; SI-Fig. 2). The TKTD population model was used to run simulations for the time-period 1966-2015, where data on PCB levels in prey and seals were available. A five-year pre-simulation period (1961-1965) was included to initiate realistic values of PCB, damage, and hazard/stress levels. Since PCB levels in Baltic fish were low in the early 1960s (Bignert et al. 1998), all PCB, damage, and hazard/stress levels were put to zero at the start of year 1961. Prey concentrations were assumed to increase linearly from zero (at 1961) to the reported levels 1966. According to data from Harding et al. (2007), the total population size year 1961 was 17 639 seals (including males and females). The initial population size in the simulation (accounting for females exclusively) was put to half that amount. Simulation outputs for temporal PCB levels in seals and population size were compared to empirical data.

## Results

### PCB and population dynamics with constant exposure

Simulating time-constant prey PCB concentrations in our population TKTD model results in steady-state dynamics for PCB and population metrics over time (Fig. 2). The mean biomagnification of PCBs from fish to seals (all age classes included) increased with prey PCB concentrations up to a plateau of approximately 35 as prey levels reach 20 mg/kg lw (Fig.2a). Assuming only TK processes (no TD effects), the biomagnification factor remains constant at 11 independent of prey PCB concentrations (Fig. 2a). Using the observed range of mean prey PCB levels since the 1960s (0-10 mg/kg lw), steady-state mean PCB levels for seals of all age classes over a 100-year simulation period were predicted (Fig. 2b). Here, PCB levels in seals increase over time for each constant diet exposure scenario and reach steady-state levels after 10 to 40 years. In accordance with the biomagnification results, higher prey PCBs resulted in higher seal PCB levels and longer times to reach steady state. While population sizes approach different stable levels for each constant diet exposure, the population growth rate decreases over time and approaches zero due to PCB impacts on fertility and survival (Fig. 2c). In the absence of PCBs, the population approaches its carrying capacity of 100000 animals and increasing exposure to dietary PCBs reduces relative population size and growth rates, with a constant dietary level of approximately 4 mg/kg lw representing a critical threshold in which the growth rate is negative and the population size starts to decline over time (Fig. 2c).

**Fig. 2.**
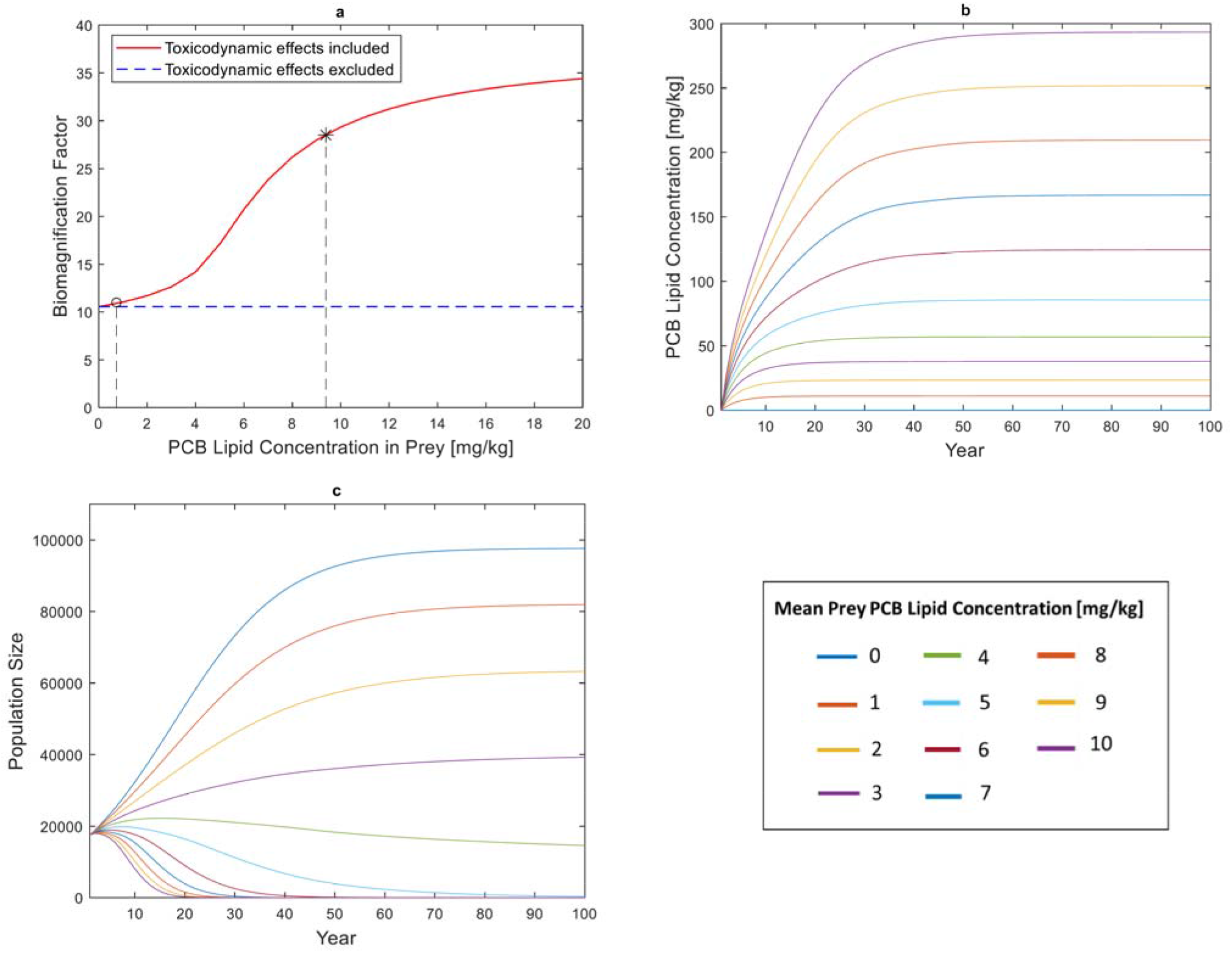
Steady State model results. a) Mean biomagnification factors (from prey to seals) predicted by the TKTD population model under steady state conditions for different levels of constant mean PCB lipid concentration in prey. Different results are obtained depending on whether toxicodynamic effects (adverse effects of PCB on fertilities and survival rates) are included. Indicated are the historical peak level of 9.4 mg/kg (*) and the 2015 level of 0.73 mg/kg (o). b) Mean PCB lipid concentration in seals for different scenarios of constant PCB lipid concentrations in prey (ranging from 0 to 10 mg/kg) for a 100-year simulation period. PCB concentrations in seals are zero at the start of the first year. c) Population size in seals for different scenarios of constant PCB lipid concentrations in prey (ranging from 0 to 10 mg/kg) for a 100-year simulation period.

### PCB and population dynamics with time varying exposure

We used historical fish PCB data on three important prey items for Baltic grey seals (Fig. 3a) to model temporal PCB dynamics in grey seal females across their full life cycle (Fig. 3b-d). The predicted lipid concentrations of PCBs in female seals are higher when TD processes are included (full model) as compared to a model with only TK process (Fig. 3b-c). Model predicted PCB levels followed observed temporal patterns for all seals (Fig. 3b) and especially for juveniles (Fig. 3c). Modeled PCBs in seals for all age-classes followed observed declines in prey levels over time, though empirical data for seals fluctuated to much greater degree across years at short and long timespans (Fig. 3b). The peak PCB level in seals neared 120 mg/kg lw around 1976, which was several years later than peak PCB levels in sprat and herring but similar to cod (Fig. 3a-c). Following lifetime PCB profiles of seals born at different years revealed contrasting patterns over time (Fig. 3d). Consistent across almost all cohorts is the large vertical transfer of PCBs from mother to pup during lactation that results in peak lifetime PCB levels in pups. This is followed by a period of growth dilution when PCB levels decline as the animal grows rapidly during juvenile years, and finally by decreasing concentrations in adults over time because of recurring vertical transfers (Fig. 3d). The 1966 cohort is unique as it was exposed to the greatest historical levels of PCBs and consequently had the greatest PCB impacts on reproduction, which was captured by the increasing PCB levels in young adult females that could not reproduce and offload their PCB burdens (Fig. 3d).

**Fig. 3.**
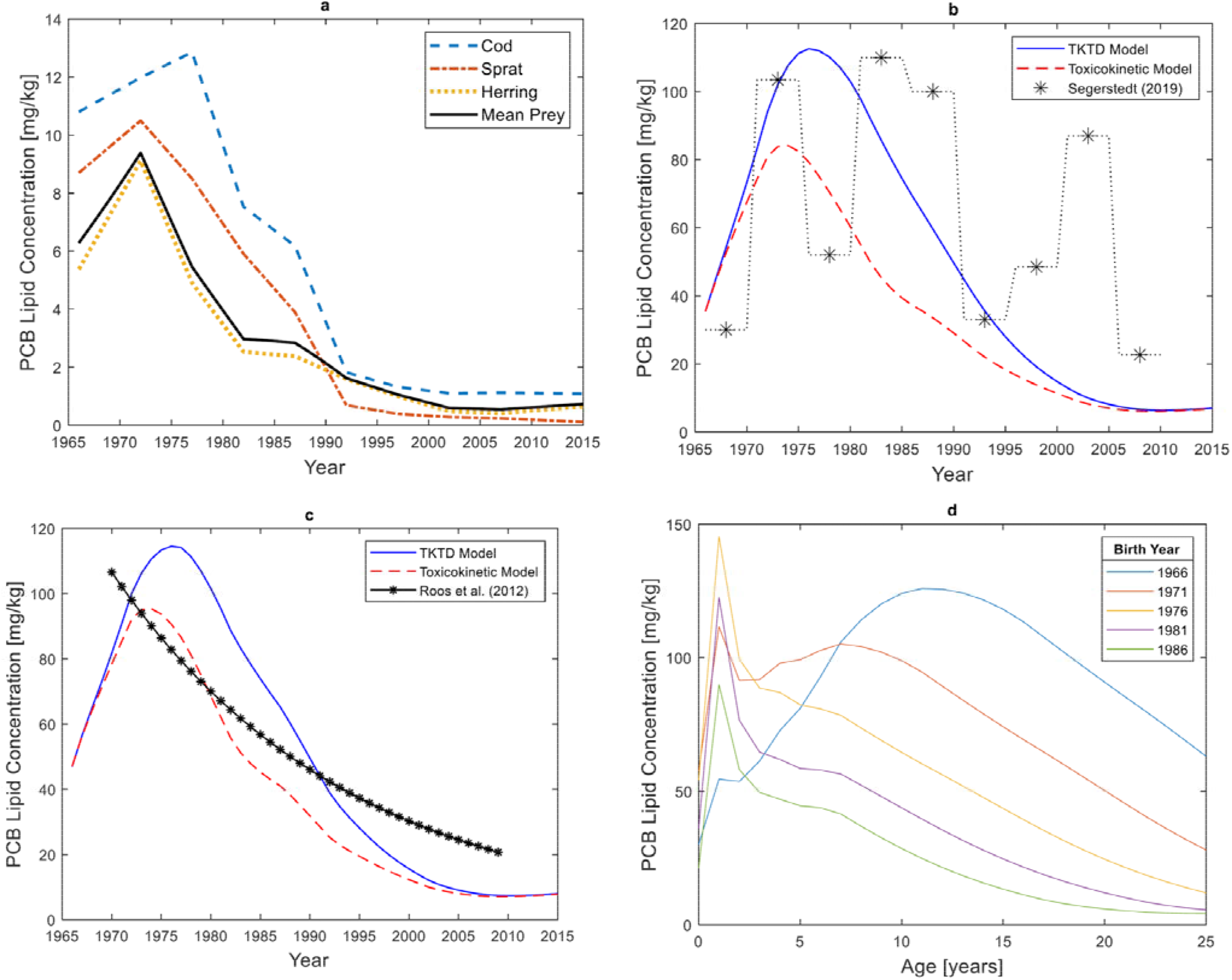
Time-varying dietary exposure model results for PCB dynamics. a) Temporal sample mean PCB lipid concentrations in different prey species from the Baltic Sea during 1966-2015, according to data compiled by Segerstedt (2019). b) Mean PCB lipid concentrations in female Baltic grey seals (i=1-46) between 1966 and 2015. Shown are model predictions and published data (Segerstedt 2019). c) Mean PCB lipid concentrations in juvenile Baltic grey seals (*i*=1,2,3) according to model predictions and published data (Roos et al. 2012). d) Mean age-specific PCB lipid concentrations in different cohorts of female Baltic grey seals between 1966 and 2015.

Empirical data on pregnancy rates in Baltic grey seals showed near zero values in the 1970s and a return to near 100% in 2015 (Fig. 4a). Model predictions for pregnancy rates followed the temporal increase in pregnancy rates observed in the empirical data, though the model overpredicted pregnancy rates in earlier years of the dataset and underpredicted them in later years (Fig. 4a). Translating PCB effects to population dynamics, the TKTD model accurately predicted temporal changes in the total population size, albeit with a time-lag in the population decline in the 1960s and 1970s (Fig. 4b). Comparing cohorts, the model showed that fertility rates were lowest in the first and most highly exposed cohort of 1966 and improved thereafter (Fig. 4c). In all cohorts, fertility increased with age, with the exception of the 1966 cohort where fertility drops between age 7 and 11 before increasing again. The 1986 cohort eventually reached the ideal fertility rate of an exposed population in its mature females. Survival rates were predicted to be quite low in all cohorts for the youngest age classes and increased substantially in juveniles and adults (Fig. 4d). Not surprisingly, survival was lowest in early cohorts that were exposed to the highest historical levels of PCBs through their diet.

**Fig. 4.**
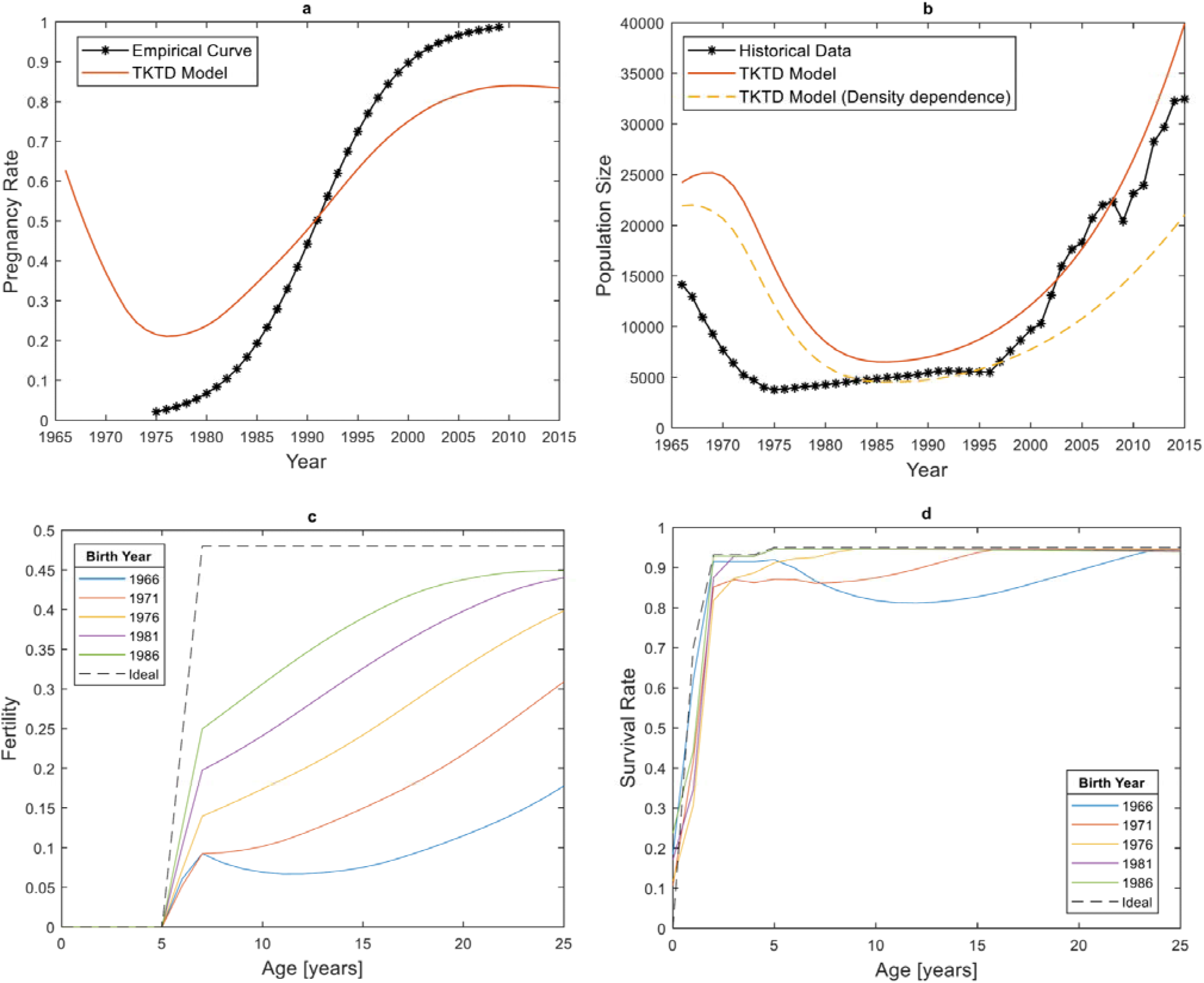
Time varying dietary exposure model results for population dynamics between 1966 and 2015. a) Pregnancy from Roos et al. (2012) and mean pregnancy rates for females of ages 5-46 years according to the TKTD population model. b) Population size according to historical data (Harding et al. 2007) and the TKTD population model. c) Age-specific fertility rate in different cohorts of female Baltic grey seals between 1966 and 2015. d) Age-specific survival rate in different cohorts of female Baltic grey seals between 1966 and 2015.

### Time to removal analysis

We modeled PCB dynamics in Baltic grey seals under theoretical scenarios of complete PCB elimination in the environment to assess the importance of vertical intergenerational transfer on long-term contaminant trends (Fig. 5). Assuming PCB exposure stopped at peak levels in 1976 where seal PCB concentrations reached on average 113 mg/kg (according to model), model simulations revealed concentrations in the seal population reached below 0.1 mg/kg (near zero) after 28 years, corresponding to a yearly decline of 23 % (Fig. 5a). A similar exercise assuming PCB exposure stopped at current PCB tissue concentrations of 7 mg/kg (predicted by the model), equivalent to model year 2015, simulations found that PCB concentrations reached near zero after 15 years, corresponding to a yearly decline of 26 % (Fig. 5b).

**Fig. 5.**
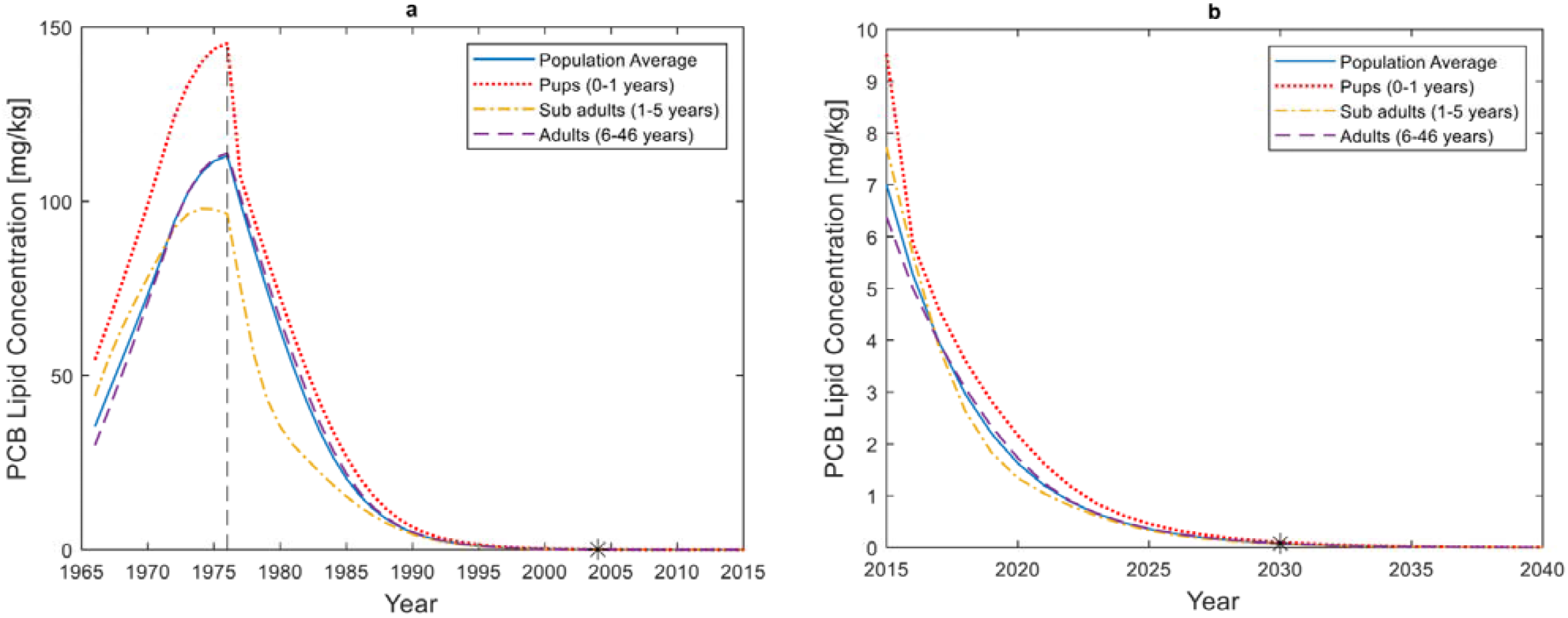
Scenarios with initial historical prey PCB exposure levels, followed by zero prey PCB exposure. PCB levels are average values for all age classes according to TKTD model. a) PCB exposure stops at the peak PCB concentration in seals (113 mg/kg) at year 1976. b) PCB exposure stops at year 2015 (with a PCB concentration of 7 mg/kg in seals).

### Sensitivity Analysis

A sensitivity analysis was carried out to evaluate the impact of TK and TD variable uncertainty on the stable population growth rate (*λ*) of Baltic grey seals (Fig. 6). For the TK parameters, the sensitivity analysis showed that *λ* decreased with increased placental PCB transfer from female to embryo during gestation, whereas it increased with increased milk PCB transfer from female to pup during lactation (Fig. 6a). The *λ* also increased with increased PCB metabolic transformation (Fig. 6a).

**Fig. 6.**
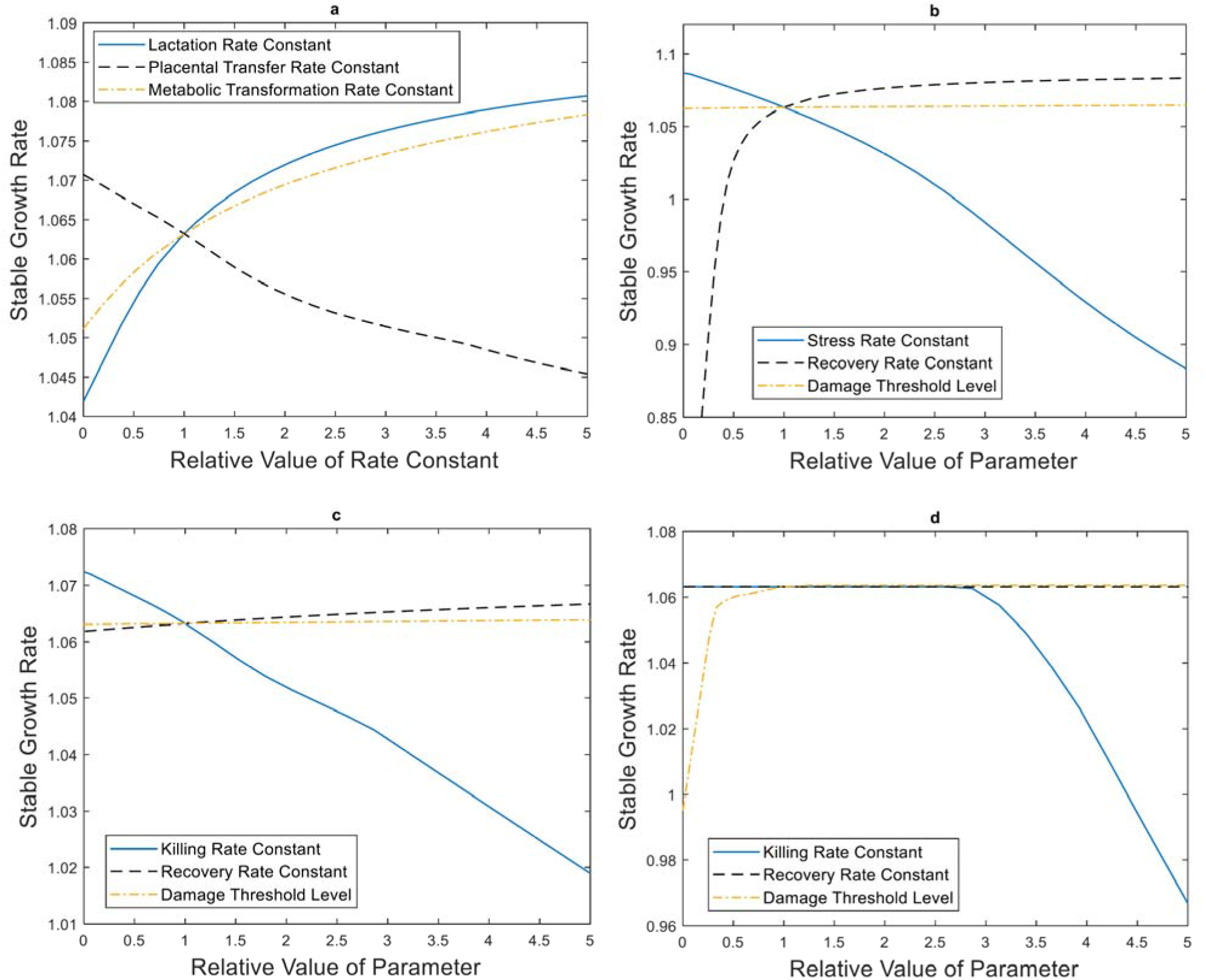
Sensitivity analysis of TKTD parameters on the stable population growth rate after long-term exposure to constant mean prey PCB concentrations (2 mg/kg lw). a) Effect of TK parameters. b) Effect of TD parameters related to reproductive stress. c) Effect of TD parameters related to fetal survival. d) Effect of TD parameters related to female survival (*i* > 0).

The effect of TD parameter variability was estimated for reproductive stress parameters (Fig. 6b), fetal survival parameters (Fig. 6c), and non-fetus female survival (Fig. 6d). For reproductive stress, *λ* was maximal (*λ* = 1.09) when the stress rate constant was zero and decreased with increased values of the stress rate constant. *λ* initially increased rapidly with increased recovery rate constants, but then approached maximum growth, corresponding to a state where accumulated damage was immediately recovered. *λ* increased very slowly with increasing damage threshold levels until the max *λ* was reached. For fetal survival, *λ* decreased with increased killing rate constants and increased slowly with both increased recovery rate constants and increased damage threshold levels. For survival in all other age-classes, *λ* was initially unaffected by increased killing rate constants since the damage threshold level was not exceeded, though eventually decreased with greater killing rate constants. *λ* initially increased rapidly with increased damage threshold levels, but when accumulated damage exceeded threshold levels in all age classes, the increase reached a plateau. Finally, *λ* was unaffected by changes in the recovery rate constant since the other parameters had values such that no damage was accumulated (the threshold level was not exceeded).

## Discussion

We developed a toxicokinetic model for dietary uptake, elimination, and vertical transfer of PCBs by grey seals. This toxicokinetic model was in turn linked to a toxicodynamic model in order to estimate the resulting adverse effects on grey seal fertility and survival rates. Finally, the toxicodynamic model was linked to an age structured Leslie matrix population model with the aim to estimate population responses from different pollutant exposures. This three-step process is a toxicokinetic-toxicodynamic-population-model and we term it a “TKTD-population model”. Most models for adverse effects of toxicants on populations are based on empirically obtained dose-response relationships, instead of some underlying theory for toxicity (such as DEBtox or TDM). The new model is the first to combine mechanistic descriptions of both toxicokinetics and toxicodynamics and link them to effects at the population level for a marine mammal, applying the Leslie matrix approach which facilitates computation compared to earlier work on modelling effects from PCB exposure in marine mammals by individual based models (Hall et al. 2006a, Hall et al. 2006b, Hall 2013a, Hall 2013b, Desforges et al. 2018). The TKTD-population model also captures vertical toxicant transfer between age-classes, which enables detailed predictions of toxicant concentrations in animals of different age classes over long time periods, avoiding the computerpower-consuming computations of an individual-based model (IBM).

We explored how different constant exposure concentrations affect population growth rate. Steady state analyses can be used to theoretically analyse how large contaminant concentrations a population can resist over time. According to the model, the Baltic grey seal population crashes if the mean prey PCB lipid concentration is 4 mg/kg or more. This level was found in Baltic fish in the early eighties (Fig. 3a). The level peaked at 9.4 mg/kg in the early seventies and if this level had remained, the population is predicted to have gone extinct 20 years later.

### Sensitivity Analyses

Our sensitivity analysis of model parameters showed that the population growth rate increased with increased vertical PCB transfer from mothers to pups during lactation (Fig. 6a). Increased vertical transfer decreases female PCB body burdens and increases female survival, while it increases pup PCB body burdens and decreases pup survival. The result is a net positive effect on population growth. On the contrary, population growth rate decreased with increased placental PCB transfer from mothers to fetuses during gestation (Fig. 6a). This result is due to the fact that placental transfer has a minor role in female detoxification, whereas fetuses are small and sensitive to PCB burden. Obviously, population growth rate increased with increased metabolic PCB transformation with only positive effects on PCB elimination (Fig. 6a). The model does not however account for the fact that some metabolites may be more toxic than the parent substance (Eisler and Belisle 1996).

Among the toxicokinetic rate constants considered in the sensitivity analyses (*k_L,i_, k_p,i_, k_M,i_*), the model was most sensitive to changes in lactation rate constants (Fig. 6a), but estimations of vertical transfer rate constants are based on detailed empirical data for grey seals (Lang et al. 2011, Berghe et al. 2012) and are probably quite valid. The estimation of metabolic transformation rate constants is more uncertain. The total PCB elimination rate constant for arctic ringed seals (Hickie et al. 2005) was adopted for 60 kg grey seals; *k_M,RS_* = 0.17/*year*. In their bioaccumulation model, Hickie et al. (2005) calculated estimations of elimination rate constants for different PCB congeners, ranging from 0.03/year to 2.5/year. Changes of *k_M,RS_* within this span (corresponding to relative values 0.18 ≤ *k_rel_* ≤ 15) yield non-negligible effects on stable growth rate (Fig. 6a). With *k_rel_* = 5, stable growth rate is *λ* = 1.078, to be compared with *λ* = 1.063 for *k_rel_* = 1. However, since good agreement was obtained between model predictions and observed PCB levels in Baltic grey seal juveniles (Fig. 3c), the estimation of metabolic transformation rate constants is likely valid.

### Toxicokinetic Model Outputs

Predicted seal/prey PCB concentration ratios in year 2001 (ranging from 15 to 19), were lower than the PCB biomagnification factor (59±40), calculated by Routti et al. (2005). The model also predicted lower PCB concentrations in seals during 1966-2015 compared to data (Fig. 3b). These findings are expected, since the TKTD population model only includes females, whereas Routti et al. (2005) and Segerstedt (2019) based their results on data for both genders. Males accumulate PCBs throughout their lifetime whereas female PCB levels stabilize over time due to vertical transfer. Hence, studies including males generally report higher mean PCB concentrations. Better agreement between model and data is expected for PCB concentrations in juveniles (0-3 years old), which have not yet transferred PCBs to offspring. Gender difference between juveniles of the same age is primarily a matter of body weight and similar PCB concentrations can be expected. Accordingly, temporal PCB concentrations predicted by the model agree well with empirical data for juveniles (Fig. 3c).

The TKTD population model links toxicokinetics to toxicodynamics, accounting for adverse effects of PCB on fertility and survival rates. An age structure different from that of a healthy population is then obtained. Since toxicokinetics differ between age classes and vertical transfer from mother to offspring is an important mechanism, toxicokinetics on the population level is altered by the toxicodynamics. Some model outputs were compared with and without toxicodynamic influence, including biomagnification factors (Fig. 2a) and PCB concentrations (Fig. 3b-c). When toxicodynamic effects were included, higher PCB accumulation was predicted. Moreover, the biomagnification factor increased with prey PCB concentration. These findings are primarily explained by changes in vertical transfer. Fertility rates decreased with increasing PCB exposure. Since PCB transfer from females to offspring then decreases, bioaccumulation in females increases. Since pup survival is considerably lower than adult survival (SI-Table 4), PCBs are more effectively eliminated from the population (through vertical transfer) when fertilities are high.

### Toxicodynamic Model Outputs

Model predicted population sizes between 1966 and 2015 started to decline later than the observed historical trend (Fig. 4b). This is expected since the historic population size is reduced by hunting and by-catches, effects not included in the TKTD population model. Inclusion of density dependence yields predictions with minimum levels closer to historical data, but also slower recovery during later years, where curves depart from the historical trend. The adopted carrying capacity of 100 000 animals is probably not to low, corresponding to the population size in the early 1900s when conditions probably where more favorable for Baltic grey seals. However, values on density parameters affecting adult survival rates may have been put too high in relation to density parameters affecting fertilities and pup survival. Accurate relative adjustments of density parameters, without changing the carrying capacity, would probably give a more realistic population recovery and a bottom population size above historical data. However, with inclusion of hunting and by catches, better agreement is expected.

### Model Assumptions and Limitations

#### Population-Based Model

The TKTD population model is a population-based model (PBM). More details could have been included if an individual-based model (IBM) had been adopted, but the simplicity of a PBM allows easier interpretation of model outputs and much faster computation. It is also possible to extend the TKTD population model with more details, such as using a multicompartment model to describe toxicokinetics.

In PBMs based on mechanisms at the individual level, the *pooling effect* occurs (Klanjscek et al. 2006). Properties are expressed as mean values for different age classes, but a real population has individual variation within each age class. Some individuals may have considerably higher toxicant and hazard/stress levels than model predictions. In IBMs, individuals with a high level of accumulated hazard/stress die or become sterile before they have accumulated enough energy to reproduce. In PBMs, negative effects on fertility are averaged over many individuals and the energy for reproduction is pooled from all adults without losses. Hence, fertility is overestimated and extinction risks may be underestimated in population viability analyses.

#### Measurement of PCB Body Burden

Since PCBs mainly accumulate in the blubber of marine mammals, it was assumed that blubber concentration could be used as an index for total body burden. A simple approach for conversion between total body concentration and lipid concentration was adopted. This may not always be accurate since lipophilic contaminants are slowly excreted, whereas blubber thickness can change fast. An emaciated animal with a thin blubber layer may have a very high blubber PCB concentration, although the total body burden is similar to that of an animal in normal nutritional condition (Bergman 2007). However, conversion of concentrations is only used for comparison with empirical data, primarily collected in the autumn when blubber layers are thick.

In the model, total body PCB concentration governs cumulation of damage, whereas toxic effects of PCB on real individuals are primarily governed by the concentration in the blood (Klanjscek et al. 2007). While these are usually correlated, the current model may be refined by considering distribution of PCBs among different tissues.

The composition of fish species in the Baltic Sea have changed throughout time (HELCOM 2018b). The simulations performed here are based on only three prey species and fixed prey preference indices for each age class. However, it might be of interest to model temporal changes in prey composition since different fish species are associated with different contaminant loads.

#### Metabolic Transformation Rate Constants

Kleiber’s law for allometric scaling of metabolic rate (Kleiber 1962) was used to scale metabolic transformation rate constants from ringed seals to grey seals and between age classes of grey seals. Since grey seals are larger than ringed seals, they have a lower mass-specific metabolic rate and should thus have a lower capacity to eliminate PCBs per unit of body mass. This is empirically supported by some publications. From collected samples, Routti et al. (2005) found that bioaccumulation was higher in Baltic grey seals than in Baltic ringed seals, indicating differences in the metabolic system. Routti et al. (2005) also showed that biomagnification of PCBs in Baltic seals was very high compared to Arctic seals. They suggested that Baltic seals have higher capacity to metabolize PCBs due to increased expression of the metabolizing enzymes, a response to the heavy contaminant burden in the Baltic Sea (Routti et al. 2005). On the contrary, other studies indicate that the induction capacity of the important xenobiotic metabolising enzyme CYP1A is better in grey seals than in ringed seals (Nyman 2000). There are indications that grey seals have a specific mechanism for metabolism of the very toxic congener IUPAC 118 (Roots and Talvari 1997). Obviously, the adopted method of scaling elimination rate constants between seal species may be further elaborated with appropriate empirical evidence.

To obtain a more realistic population growth rate, relative adjustments of density parameters should be performed, without changing the carrying capacity. Model parameters can be estimated by adopting a maximum likelihood method where model outputs are compared to empirical data and uncertainties in estimated parameters can be analysed by Monte Carlo simulations, an approach previously adopted for viability analysis of fish populations (Xiao et al. 2008).

#### Toxicodynamic Model

The adopted toxicodynamic model is based on the threshold damage model (TDM), including damage accumulation, damage threshold levels, and recovery. Uterine occlusions in Baltic grey seal females occurred at earliest in 7 year old females, whereas uterine leiomyoma usually appeared after 15 years of age (Bergman 2007). Leiomyoma prevalence increased with life-time exposure (Bredhult et al. 2008). These findings motivate a model where damage accumulates over time and adverse effects occur when threshold levels are reached. Grey seal females can probably recover from sterility caused by PCB exposure when PCB concentrations decrease, at least to some extent. PCBs also cause adverse effects on the immune system of grey seals adding an extra maintenance cost acting on survival reducing the energy available to deal with for example hook worms and other pathogens (Klanjscek et al. 2007). These immune effects were not included in the current model design.

#### Parameterization of the Toxicodynamic Model

Reproductive stress parameters were calibrated by fitting predicted pregnancy rates to observed pregnancy rates in Baltic grey seals (Fig. 4a). However, the only negative effect on pregnancy rates in the model is PCB exposure, whereas observed pregnancy rates may have been affected also by other contaminants (like DDT) or other factors, possibly in interaction with each other. Despite parameter adjustments, the model for negative effect of reproductive stress on fertility could not generate the fast reduction in pregnancy rate seen in empirical data between mid-60s and mid-70s.

No DEBtox or TDM parameters for grey seals affected by PCBs are currently available in the literature, but a DEB model for mink, accounting for adverse effects of PCBs on fecundity and kit survival, has previously been developed (Desforges et al. 2017). Since there is a strong correlation between mass-specific metabolic rate and sensitivity to a toxicant (Baas and Kooijman 2015), we consider it wise to apply Kleiber’s law to perform allometric scaling of some TDM parameters. We suggest ways to translate DEBtox parameters between species and relate them to TDM parameters in SI. The parameter values adopted for the TKTD population model should be considered as a starting point for future model refinements.

### Possible Model Extensions

#### Multi-Compartment Model

In the model, blubber thickness is a specified fraction of the body weight. However, blubber thickness changes during the year and is density dependent. An age-specific model for annual variation of blubber weight is a simple approach that can be immediately implemented to the current model without modification of equations. It is also possible to describe the toxicokinetics by a *multi-compartment model* that differentiates bioaccumulation in different tissues and link them through diffusion rates. The simplest version is a *two-compartment model*, where the seal body is divided into *structure* (core) and *reserve* (blubber), with differentiated growth, PCB kinetics and damage effects.

#### Combination of Risk Factors

Besides PCBs there may be other contaminants with adverse effects on fertility and survival rates, possibly in interaction. DEBtox models and TDMs allow inclusion of combined effects of multiple toxicants. The accumulated damage from the mixture of all toxicants may be calculated as the sum of the internal damages from all toxicants separately (Ashauer et al. 2007a). However, this effect is only additive. There are also ways of accounting for interaction between different compounds, such as using non-linear stress functions in DEBtox models (Kooijman 2000). Hunting, by-catches and decreasing ice coverage (due to climate change) may also be relevant aspects to consider in population viability analyses of Baltic grey seals. The population is currently faced with a sharp increase in hunting quotas and also vulnerable to declining ice fields due to global warming in addition to prevailing high levels of POPs, it will be important to elaborate scenarios with all these stressors in combination (Cervin et al. 2020, Silva et al. 2021) order to avoid a new era of overexploitation.

## Acknowledgments

Funding was provided by Viltforskningsanslaget, Swedish Environmental Protection Agency, and by the BONUS program BaltHealth (Art. 185).

## Figure Captions

Fig. 1 was created with PowerPoint. Fig. 2-6 were created with Matlab.

## Statements & Declarations

### Author Contributions

Model development was primarily made by Karl Mauritsson, but with important contributions from the other authors. All simulations were performed by Karl Mauritsson. The manuscript was written by Jean-Pierre Desforges and Karin C. Harding, based on a previous report written by Karl Mauritsson. All authors commented on previous versions of the manuscript. All authors read and approved the final manuscript.

## References

Ashauer, R., A. B. Boxall, and C. D. Brown. 2007a. Modeling Combined Effects of Pulsed Exposure to Carbaryl and Chlorpyrifos on Gammarus Pulex. Environmental Science & Technology 41:5535–5541. doi:10.1021/es070283w

Ashauer, R., A. B. Boxall, and C. D. Brown. 2007b. New Ecotoxicological Model To Simulate Survival of Aquatic Invertebrates after Exposure to Fluctuating and Sequential Pulses of Pesticides. Environmental Science & Technology 41:1480–1486. doi:10.1021/es061727b

Baas, J., and S. A. L. M. Kooijman. 2015. Sensitivity of animals to chemical compounds links to metabolic rate. Ecotoxicology 24:657–663. doi:10.1007/s10646-014-1413-5

Berghe, M. V., L. Weijs, S. Habran, K. Das, C. Bugli, and J.-F. Rees. 2012. Selective transfer of persistent organic pollutants and their metabolites in grey seals during lactation. Environment International 46:6–15. doi:10.1016/j.envint.2012.04.011

Bergman, A. 2007. Pathological Changes in Seals in Swedish Waters: The Relation to Environmental Pollution - Tendencies during a 25-year Period. Swedish University of Agricultural Sciences, Uppsala, Sweden, PHD thesis.

Bergman, A., and M. Olsson. 1986. Pathology of Baltic grey seal and ringed seal females with special reference to adrenocortical hyperplasia: Is environmental pollution the cause of a widely distributed disease syndrome? Finn. Game Res. 44:47–62.

Bignert, A., M. Olsson, W. Persson, S. Jensen, S. Zakrisson, K. Litzfn, U. Eriksson, L. Häggberg, and T. Alsberg. 1998. Temporal trends of organochlorines in Northern Europe, 1967-1995. Relation to global fractionation, leakage from sediments and international mesures. Environmental Pollution 99:177–198. doi:10.1016/S0269-7491(97)00191-7

Bjurlid, F., A. Roos, I. E. Jogsten, and J. Hagberg. 2018. Temporal trends of PBDD/Fs, PCDD/Fs, PBDEs and PCBs in ringed seals from the Baltic Sea (Pusa hispida botnica) between 1974 and 2015. Science of the Total Environment 616:1374–1383. doi:10.1016/j.scitotenv.2017.10.178

Bredhult, C., B.-M. Bäcklin, A. Bignert, and M. Olovsson. 2008. Study of the relation between the incidence of uterine leiomyomas and the concentrations of PCB and DDT in Baltic gray seals. Reproductive Toxicology 25:247–255. doi:10.1016/j.reprotox.2007.11.008

Bäcklin, B.-M., L. Eriksson, and M. Olovsson. 2003. Histology of Uterine leiomyoma and occurrence in relation to reproductive activity in the Baltic gray seal (Halichoerus grypus). Veterinary Pathology 40:175–180. doi:10.1354/vp.40-2-175

Bäcklin, B.-M., C. Moraeus, A. Roos, E. Eklöf, and Y. Lind. 2011. Health and age and sex distributions of Baltic grey seals (Halichoerus grypus) collected from bycatch and hunt in the Gulf of Bothnia. Journal of Marine Science 68:183–188. doi:10.1093/icesjms/fsq131

Cervin, L., T. Harkonen, and K. C. Harding. 2020. Multiple stressors and data deficient populations; a comparative life-history approach sheds new light on the extinction risk of the highly vulnerable Baltic harbour porpoises (Phocoena phocoena). Environment International 144. doi:10.1016/j.envint.2020.106076

Desforges, J. P., A. Hall, B. McConnell, A. Rosing-Asvid, J. L. Barber, A. Brownlow, S. De Guise, I. Eulaers, P. D. Jepson, R. J. Letcher, M. Levin, P. S. Ross, F. Samarra, G. Vikingson, C. Sonne, and R. Dietz. 2018. Predicting global killer whale population collapse from PCB pollution. Science 361:1373–1376. doi:10.1126/science.aat1953

Desforges, J. P., P. S. Ross, and L. L. Loseto. 2012. Transplacental transfer of polychlorinated biphenyls and polybrominated diphenyl ethers in arctic beluga whales (Delphinapterus leucas). Environmental Toxicology and Chemistry 31:296–300. doi:10.1002/etc.750

Desforges, J. P., C. Sonne, and R. Dietz. 2017. Using energy budgets to combine ecology and toxicology in a mammalian sentinel species. Scientific Reports 7:46267. doi:10.1038/srep46267

Eckhéll, J., P. Jonsson, M. Meili, and R. Carman. 2000. Storm Influence on the Accumulation and Lamination of Sediments in Deep Areas of the Northwestern Baltic Proper. Ambio 29:238–245. doi:10.1639/0044-7447

Eisler, R., and A. A. Belisle. 1996. Planar PCB Hazards to Fish, Wildlife and Invertebrates: A Synoptic Review. Patuxent Wildlife Research Center.

Engen, S., R. Lande, B.-E. Sæther, and H. Weimerskirch. 2005. Extinction in relation to demographic and environmental stochasticity in age-structured models. Mathematical Biosciences 195:210–227. doi:10.1016/j.mbs.2005.02.003

Hall, A. J., K. Hugunin, R. Deaville, R. J. Law, C. R. Allchin, and P. D. Jepson. 2006a. The risk of infection from polychlorinated biphenyl exposure in the harbor porpoise (Phocoena phocoena): A case-control approach. Environmental Health Perspectives 114:704–711. doi:10.1289/ehp.8222

Hall, A. J., B. J. McConnell, T. K. Rowles, A. Aguilar, A. Borrell, L. Schwacke, P. J. H. Reijnders, and R. S. Wells. 2006b. Individual-Based Model Framework to Assess Population Consequences of Polychlorinated Biphenyl Exposure in Bottlenose Dolphins. Environmental Health Perspectives 114:60–64. doi:10.1289/ehp.8053

Hall, A. J., B. J. McConnell, L. H. Schwacke, G. M. Ylitalo, R. Williams, and T. K. Rowles. 2018. Predicting the effects of polychlorinated biphenyls on cetacean populations through impacts on immunity and calf survival. Environmental Pollution 233:407–418. doi:10.1016/j.envpol.2017.10.074

Hall, A. J. K., Joanna L.; Schwacke, Lori H.; Ylitalo, Gina; Robbins, Jooke McConnell, Bernie J.; Rowles, Teri K. 2013a. Assessing the Population Consequences of Pollutant Exposure in Cetaceans (Pollution 2000+) - from Ingestion to Outcome. SC/E65A/E4.

Hall, A. J. S., Lori H.; Kershaw, Joanna K.; McConnell, Bernie J.; Rowles, Teri K. 2013b. An Individual Based Modelling Approach to Investigate the Impact of Pollutants on Cetacean Population Dynamics - Effects on Calf Survival and Immunity. SC/64/E5.

Hansson, S., U. Bergström, E. Bonsdorff, T. Härkönen, N. Jepsen, L. Kautsky, K. Lundström, S.-G. Lunneryd, M. Ovegård, J. Salmi, D. Sendek, and M. Vetemaa. 2017. Competition for the fish - fish extraction from the Baltic Sea by humans, aquatic mammals, and birds. ICES Journal of Marine Science. doi:10.1093/icesjms/fsx207

Harding, K., and T. Härkönen. 1999. Development in the Baltic grey seal (Halichoerus grypus) and ringed seal (Phoca hispida) populations during the 20th century. Ambio 28:619–627.

Harding, K., T. Härkönen, B. Helander, and O. Karlsson. 2007. Status of Baltic grey seals: Population assessment and extinction risk. NAMMCO Scientific Publications 6:33–56. doi:10.7557/3.2720

HaV, and SMHI. 2022. Grey seal population data. SHARKweb.

HELCOM. 2016. Population trends and abundance of seals. HELCOM core indicator report. HELCOM.

HELCOM. 2018a. Nutritional status of seals. HELCOM core indicator report. HELCOM.

HELCOM. 2018b. State of the Baltic Sea - Second HELCOM holistic assessment 2011-2016. HELCOM.

Hickie, B. E., D. C. G. Muir, R. F. Addison, and P. F. Hoekstra. 2005. Development and application of bioaccumulation models to assess persistent organic pollutant temporal trends in arctic ringed seal (Phoca hispida) populations. Science of the Total Environment 351–352:413–426. doi:10.1016/j.scitotenv.2004.12.085

Jüssi, M., T. Härkönen, I. Jüssi, and E. Helle. 2008. Decreasing ice coverage will reduce the breeding success of Baltic grey seal (Halichoerus grypus) females. Ambio 37:80–85. doi:10.1579/0044-7447

Kauhala, K., M. P. Ahola, and M. Kunnasranta. 2012. Demographic structure and mortality rate of a Baltic grey seal population at different stages of population change, judged on the basis of the hunting bag in Finland. Annales Zoologici Fennici 49:287–305. doi:10.5735/086.049.0502

Kauhala, K., M. P. Ahola, and M. Kunnasranta. 2014. Decline in the pregnancy rate of Baltic grey seal females during the 2000s. Ann. Zool. Fennici 51:313–324. doi:10.5735/086.051.0303

Kauhala, K., B. M. Backlin, J. Raitaniemi, and K. C. Harding. 2017. The effect of prey quality and ice conditions on the nutritional status of Baltic gray seals of different age groups. Mammal Research 62:351–362. doi:10.1007/s13364-017-0329-x

Klanjscek, T., H. Caswell, M. G. Neuberta, and R. M. Nisbet. 2006. Integrating dynamic energy budgets into matrix population models. Ecological Modelling 196:407–420. doi:10.1016/j.ecolmodel.2006.02.023

Klanjscek, T., R. M. Nisbet, H. Caswell, and M. G. Neubert. 2007. A model for energetics and bioaccumulation in marine mammals with applications to the right whale. Ecological Applications 17:2233–2250. doi:10.1890/06-0426.1

Kleiber, M. 1962. The Fire of Life. John Wiley & Sons, New York.

Kooijman, S. A. L. M. 2000. Dynamic Energy and Mass Budgets in Biological Systems. Cambridge University Press, Cambridge, UK.

Lacy, R. C., P. S. Miller, and K. Traylor-Holzer. 2018. Vortex 10 - A Stochastic Simulation of the Extinction Process - User’s Manual. IUCN SSC Conservation Breeding Group & Chicago Zoological Society, Chicago.

Lang, S. L. C., and S. J. Iverson. 2009. Repeatability in lactation performance and the consequences for maternal reproductive success in gray seals. Ecology 90:2513–2523. doi:10.1890/08-1386.1

Lang, S. L. C., S. J. Iverson, and W. D. Bowen. 2011. The Influence of Reproductive Experience on Milk Energy Output and Lactation Performance in the Grey Seal (Halichoerus grypus). PLoS ONE 6:e19487. doi:10.1371/journal.pone.0019487

Lundström, K., O. Hjerne, S.-G. Lunneryd, and O. Karlsson. 2010. Understanding the diet composition of marine mammals: grey seals (Halichoerus grypus) in the Baltic Sea. ICES Journal of Marine Science 67:1230–1239. doi:10.1093/icesjms/fsq022

Lydersen, C., H. Wolkers, T. Severinsen, L. Kleivane, E. S. Nordoy, and J. U. Skaare. 2002. Blood is a poor substrate for monitoring pollution burdens in phocid seals. Science of the Total Environment 292:193–203. doi:10.1016/s0048-9697(01)01121-4

Nyman, M. 2000. Biomarkers for exposure and for the effects of contamination with polyhalogenated aromatic hydrocarbons in Baltic ringed and grey seals. University of Helsinki, Finland, PhD thesis.

Nyman, M., J. Koistinen, M. L. Fant, T. Vartiainen, and E. Helle. 2002. Current levels of DDT, PCB and trace elements in the Baltic ringed seals (Phoca hispida baltica) and grey seals (Halichoerus grypus). Environmental Pollution 119:399–412. doi:10.1016/S0269-7491(01)00339-6

Pavlova, V., V. Grimm, R. Dietz, C. Sonne, K. Vorkamp, F. F. Riget, R. J. Letcher, K. Gustavson, J. P. Desforges, and J. Nabe-Nielsen. 2016. Modeling Population-Level Consequences of Polychlorinated Biphenyl Exposure in East Greenland Polar Bears. Archives of Environmental Contamination and Toxicology 70:143–154. doi:10.1007/s00244-015-0203-2

Roos, A. M., B.-M. V. M. Bäcklin, B. O. Helander, F. F. Rigét, and U. C. Eriksson. 2012. Improved reproductive success in otters (Lutra lutra), grey seals (Halichoerus grypus) and sea eagles (Haliaeetus albicilla) from Sweden in relation to concentrations of organochlorine contaminants. Environmental Pollution 170:268–275. doi:10.1016/j.envpol.2012.07.017

Roots, O., and A. Talvari. 1997. Bioaccumulation of toxic chlororganic compounds and their isomers into the organism of Baltic grey seal. Chemosphere 35:979–985. doi:10.1016/S0045-6535(97)00183-5

Routti, H., M. Nyman, C. Bäckman, J. Koistinen, and E. Helle. 2005. Accumulation of dietary organochlorines and vitamins in Baltic seals. Marine Environmental Research 60:267–287. doi:10.1016/j.marenvres.2004.10.007

Segerstedt, R. 2019. PCB’s Bioaccumulation Levels in the Food Chain of Marine Top Predators in the Baltic Sea. University of Gothenburg, Gothenburg, Sweden, Degree project for Master of Science in Conservation Biology.

Silva, W., E. Bottagisio, T. Harkonen, A. Galatius, M. T. Olsen, and K. C. Harding. 2021. Risk for overexploiting a seemingly stable seal population: influence of multiple stressors and hunting. Ecosphere 12. doi:10.1002/ecs2.3343

Silva, W., K. C. Harding, G. M. Marques, B. M. Backlin, C. Sonne, R. Dietz, K. Kauhala, and J. P. Desforges. 2020. Life cycle bioenergetics of the gray seal (Halichoerus grypus) in the Baltic Sea: Population response to environmental stress. Environment International 145. doi:10.1016/j.envint.2020.106145

Sonne, C., U. Siebert, K. Gonnsen, J. P. Desforges, I. Eulaers, S. Persson, A. Roos, B. M. Backlin, K. Kauhala, M. T. Olsen, K. C. Harding, G. Treu, A. Galatius, E. Andersen-Ranberg, S. Gross, J. Lakemeyer, K. Lehnert, S. S. Lam, W. X. Peng, and R. Dietz. 2020. Health effects from contaminant exposure in Baltic Sea birds and marine mammals: A review. Environment International 139. doi:10.1016/j.envint.2020.105725

Sundqvist, L., T. Harkonen, C.-J. Svensson, and K. C. Harding. 2012. Linking Climate Trends to Population Dynamics in the Baltic Ringed Seal: Impacts of Historical and Future Winter Temperatures. Ambio 41:865–872.

Tverin, M., R. Esparza-Salas, A. Stromberg, P. Tang, I. Kokkonen, A. Herrero, K. Kauhala, O. Karlsson, R. Tiilikainen, T. Sinisalo, R. Kakela, and K. Lundstrom. 2019. Complementary methods assessing short and long-term prey of a marine top predator □ Application to the grey seal-fishery conflict in the Baltic Sea. PLoS ONE 14:1–26. doi:10.1371/journal.pone.0208694

Vanhatalo, J., M. Vetemaa, A. Herrero, b. Aho, and R. Tiilikainen. 2014. By-Catch of Grey Seals (Halichoerus grypus) in Baltic Fisheries—A Bayesian Analysis of Interview Survey. PLoS ONE 9. doi:10.1371/journal.pone.0113836

Xiao, Z., Z. Yalan, and L. Shiyu. 2008. Ecological Risk Assessment of DDT Accumulation in Aquatic Organisms of Taihu Lake, China. Human and Ecological Risk Assessment 14:819–834. doi:10.1080/10807030802235268

