## Appendix for "Maternal transfer and long-term population effects of PCBs in Baltic grey seals using a new toxicokinetic-toxicodynamic population model"

Correspondence:

### 1. Model Overview and Calculation Scheme

Our matrix model, called the TKTD population model, for analysing adverse effects of PCBs on Baltic grey seals was developed to link toxicokinetics (TK) of dietary and vertically transferred PCB accumulation to toxicodynamics (TD) of adverse effects on reproduction and survival to ultimately predict changes to population demographics and growth rates (Fig. 1). The TKTD model is used to modify survival and fecundity parameters of an age-structured Leslie matrix model for an ideal Baltic grey seal population, which also includes effects of density-dependence and environmental stochasticity, due to internal PCB concentrations following DEBtox and TDM principles of damage, hazard, stress, and recovery. While the model runs on an annual basis for population dynamics, each year is divided into three periods (*lactation*, *delay* and *gestation*), representing different lifehistory phases for a reproducing grey seal female. The year starts with the *lactation period* (lasting for 18 days), during which pups are nursed by their mothers. After weaning, females mate and the *delay period* starts (lasting for 100 days), after which the fertilized egg is implanted. During the *gestation period* (lasting for 247 days), the fetus develops in the uterus. The year ends with the birth of new pups. Since all deaths are assumed to occur just before the birth events, all pups that are born have mothers that nurse them. Different TD effects of PCBs occur after specific time periods (e.g., fertility effects estimated after the gestation period).

The TKTD population model was implemented as a set of function and script files in the numerical software MATLAB® (MathWorks Inc., Natick, MA, USA) and all analyses were performed with these.

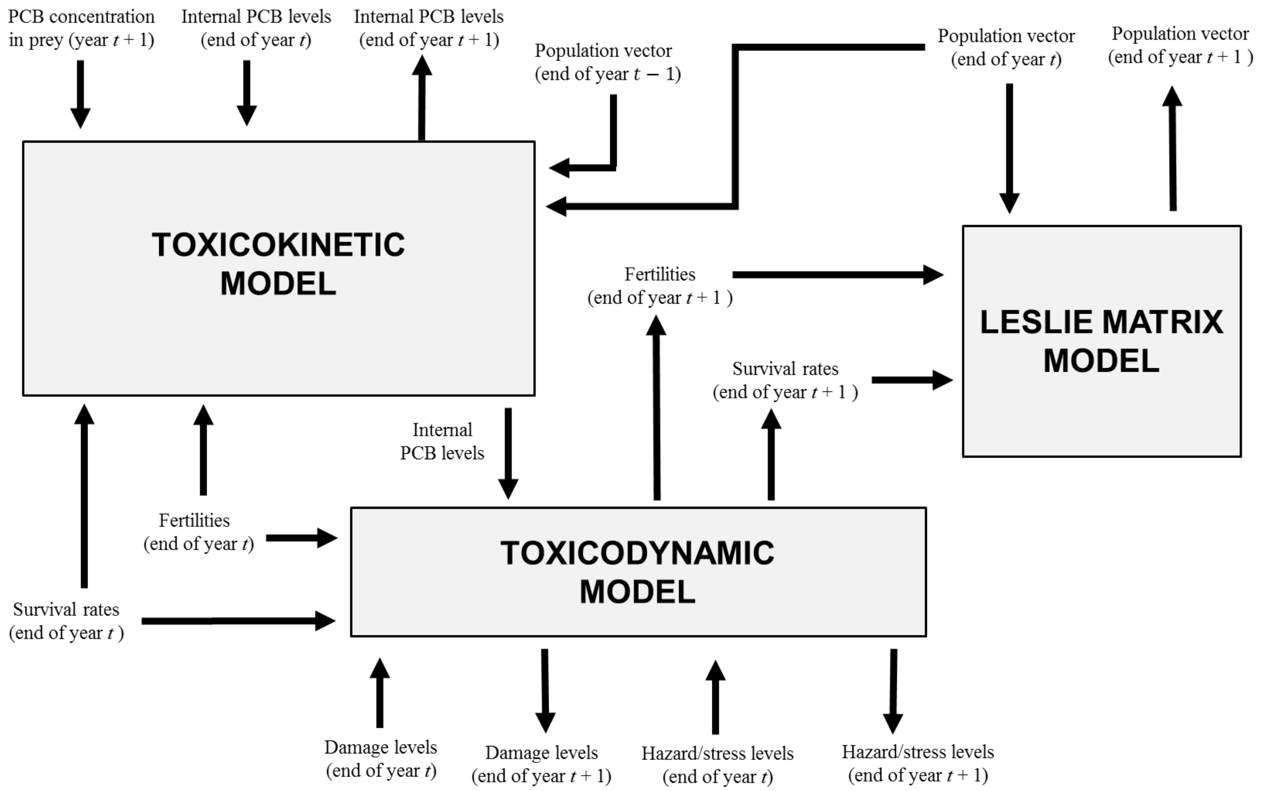

**Fig. 1** Overview of the TKTD population model. Text boxes represent sub-models, performing calculations for year $t + 1$. Free texts are inputs and outputs, with arrows showing relations to sub-models.

The model is used to perform stepwise annual calculations of PCB concentrations, which are then converted to adverse effects through calculation of damage, hazard, and stress levels (see below for details), which ultimately determine fertilities, survival rates, and number of individuals in different age classes (Fig. 1). For each year, the following calculations are performed, and the procedure is repeated until the final year:

1. Calculation of **PCB levels** in pups and females at the end of the **lactation period**, as a result of vertical transfer from females to pups, based on PCB levels in fetuses, pups and females at the end of previous year. Transfer levels are dependent on fertilities, determining the mean number of pups per female.

2. Calculation of cumulated **damage levels** in pups and females at the end of the **lactation period**, based on PCB and damage levels in fetuses, pups and females at the end of previous year.

3. Calculation of cumulated **hazard/stress levels** in pups and females at the end of the **lactation period**, based on PCB, damage and hazard/stress levels in fetuses, pups and females at the end of previous year.

4. Calculation of accumulated **PCB levels** in pups and females at the end of the **delay period**, based on PCB levels at the end of the lactation period. PCB is assimilated through dietary uptake and eliminated through metabolic transformation and fecal egestion.

5. Calculation of cumulated **damage levels** in in pups and females at the end of the **delay period**, based on PCB and damage levels in pups and females at the end of the lactation period.

6. Calculation of **cumulated hazard/stress levels** in pups and females at the end of the **delay period**, based on PCB, damage and hazard/stress levels at the end of the lactation period.

7. Calculation of reduced **fertilities** (with respect to reproductive stress) at the end of the **delay period**, based on stress levels in females (stress to the reproductive apparatus) at the end of the delay period.

8. Calculation of **PCB levels** in fetuses, pups and females at the end of the **gestation period**, as a result of bioaccumulation and placental transfer from females to fetuses, based on PCB levels in pups and females at the end of the delay period. Transfer levels are dependent on fertilities at the end of the delay period, determining the mean number of produced fetuses per female.

9. Calculation of cumulated damage **levels** in fetuses, pups and females at the end of the **gestation period**, based on PCB and damage levels in pups and females at the end of the delay period.

10. Calculation of cumulated hazard/stress levels in fetuses, pups and females at the end of the **gestation period**, based on PCB, damage and hazard/stress levels in pups and females at the end of the delay period.

11. Calculation of reduced **survival rates** at the end of the year, based on hazard levels in pups and females at the end of the gestation period.

12. Calculation of reduced **fertilities** at the end of the year, based on fertilities at the end of the delay period (affected by reproductive stress) and fetal hazard levels at the end of the gestation period.

13. Calculation of new **population vector** (accounting for population growth during one year), based on updated fertilities and survival rates, and previous year’s population vector.

#### 2. Model Variables and Parameters

The TKTD population model simulates the temporal dynamics of a number of state variables for Baltic grey seals over their full lifespan (Table 1). While the matrix model runs on an annual time step to estimate population dynamics, the TKTD model is updated three times a year to capture grey seal life history events. The three time periods of a year distinguish the delayed implantation period of seal pregnancy, the active gestation period of foetal development, and the lactation period. In order to capture the temporal dynamics of variables for seals of different age classes at different times of the year, we introduce the notations $\hat{c}_1^j(t)$ and $\hat{c}_{1,\mathrm{i}}^j(t)$, where $i = 2, \ldots, 46$. The first letter represents the variable of interest, here being seal PCB concentration in mg/kg. The upper index *j* indicates the time period of a year ($l$ = lactation, $d$ = delay, $g$ = gestation) and the lower index *i* indicates age class. Individuals of age class $i = 1$ (with an age of 0-1 year) are here referred to as *pups*, whereas individuals of age class $i = 2, \ldots, 46$ are referred to as *females*. Lastly, the PCB concentrations are calculated at time $\tau$ in years. For example, $\hat{c}_\mathrm{i}^l(t)$ and $\hat{c}_{1,\mathrm{i}}^l(t)$ would be PCB concentrations at the end of the lactation period (*l*) of year $t$ in females of age class $i$ and pups (age class = 1) with mothers of age class $i$, respectively. This convention is applied for all variables. Furthermore, we use the notation where a dot above letters indicates a rate with respect to time, a straight overline indicates an average value, and a hat over a letter denotes an estimated value.

The state variables in the TKTD model can be divided by sub-model (Table 1). For the matrix population model, we follow the number of individuals and their survival rate, fertility, and stressed fertility in each age class and the end of the year. The TK model of PCB accumulation follows the total body PCB concentrations of seals of different age classes at each time of the year. The TD model follows damage and hazard in seals of different age classes at each time of the year as well as damage and stress to reproductive tissue of females of different age classes (age class > 2) at each time of the year.

Before any calculations can be performed, a number of initial conditions have to be specified. These are initial PCB concentrations, initial damage levels, initial hazard/stress levels and the initial population vector (Table 1). Also, PCB concentrations in different prey are specified for each year of the simulation. Thereafter, PCBs concentrations, damage levels and hazard/stress levels in fetuses, pups and females at the end of the different time periods within a year are successively calculated. Fertilities and survival rates are updated based on these computations and a new population vector is calculated, including the number of individuals in different age classes at the end of a year. Outputs from calculations for one year are used as inputs to calculations for the next year. The procedure is repeated for one year at a time and may be continued for as many years as desired.

We use the TKTD population model to run simulations for the time period 1966-2015, where data on PCB levels in prey species were available. Segerstedt (2019) compiled data from previous studies of PCB concentrations in the main prey items (cod, sprat and herring) of Baltic grey seals from all regions of the Baltic Sea during 1966-2015. The data are provided as lipid weight concentrations clustered into time intervals of 5 years (Fig. 2). The prey data illustrate that PCB levels peaked in the 1970s for all three fish species. We use the time clustered prey PCB data to linearly interpolate over each five-year period to obtain yearly PCB concentrations in prey, used as input to the TK model (Fig. 2).

A five year long pre-simulation period (1961-1965) was included to initiate realistic values of seal PCB, damage and hazard levels. Since PCB levels in Baltic fish were low in the early 1960s (Bignert et al. 1998), all PCB, damage and hazard/stress levels were put to zero at the start of year 1961. Prey concentrations were assumed to increase linearly from zero (at 1961) to the reported levels in 1966. According to data from Harding, et al. (2007), the total population size year 1961 was 17 639 seals (including males and females). The initial population size in the simulation (accounting for females exclusively) was put to $N_0 = 17\,639/2 \approx 8820$.

**Table 1**. State variables and initial conditions in the TKTD model.

| variable | definition | value | unit |
|---|---|---|---|
| $N_i(t)$ | Number of individuals in age class $i$ at the end of year $t$ | - | - |
| $P_i(t)$ | Survival rate of individuals in age class $i$ at the end of year $t$ | - | - |
| $F_i(t)$ | Fertility of individuals in age class $i$ at the end of year $t$ | - | - |
| $F_i^d(t)$ | Fertility (with respect to reproductive stress) of individuals in age class $i$ at the end of the delay period year $t$ | - | - |
| $\hat{c}_{0,i}(t)$ | PCB concentration in fetuses with mothers of age class $i$ at the end of the gestation period year $t$ | - | mg/kg |
| $\hat{c}_{1,\mathrm{i}}^j(t)$ | PCB concentration in pups with mothers of age class $i$ at the end of time period $j$ year $t$ | - | mg/kg |
| $\hat{c}_i^j(t)$ | PCB concentration in individuals of age class $i$ at the end of time period $j$ year $t$ | - | mg/kg |
| $\hat{d}_{0,i}(t)$ | Damage to fetuses with mothers of age class $i$ at the end of year $t$ | - | - |
| $\hat{d}_{1,i}^j(t)$ | Damage to pups with mothers of age class $i$ at the end of time period $j$ year $t$ | - | - |
| $\hat{d}_i^j(t)$ | Damage to individuals of age class $i$ at the end of time period $j$ year $t$ | - | - |
| $\hat{\hat{d}}_i^j(t)$ | Damage to reproductive apparatus of individuals in age class $i$ at the end of time period $j$ year $t$ | - | - |
| $\hat{h}_{0,i}(t)$ | Hazard to fetuses with mothers of age class $i$ at the end of year $t$ | - | - |
| $\hat{h}_{1,i}^j(t)$ | Hazard to pups with mothers of age class $i$ at the end of time period $j$ year $t$ | - | - |
| $\hat{h}_i^j(t)$ | Hazard to individuals of age class $i$ at the end of time period $j$ year $t$ | - | - |
| $\hat{s}_i^j(t)$ | Stress to reproductive apparatus of individuals in age class $i$ at the end of time period $j$ year $t$ | - | - |
| $\hat{c}_i(0)$ | Initial PCB concentrations in pups and females | 0 | mg/kg |
| $\hat{d}_i(0)$ | Initial damage levels (Survival of pups and females) | 0 | $\mathrm{year}^{-1}$ |
| $\hat{\hat{d}}_i(0)$ | Initial damage levels (Reproductive stress) | 0 | $\mathrm{year}^{-1}$ |
| $\hat{h}_i(0)$ | Initial hazard levels (Survival of pups and adults) | 0 | - |
| $\hat{s}_i(0)$ | Initial stress levels (Reproductive stress) | 0 | - |
| $N_0$ | Initial population size (Total number of females in population) | 8820 | - |
| $\bar{c}_{D,i}(t)$ | Prey concentration function | 0-13 | mg/kg |

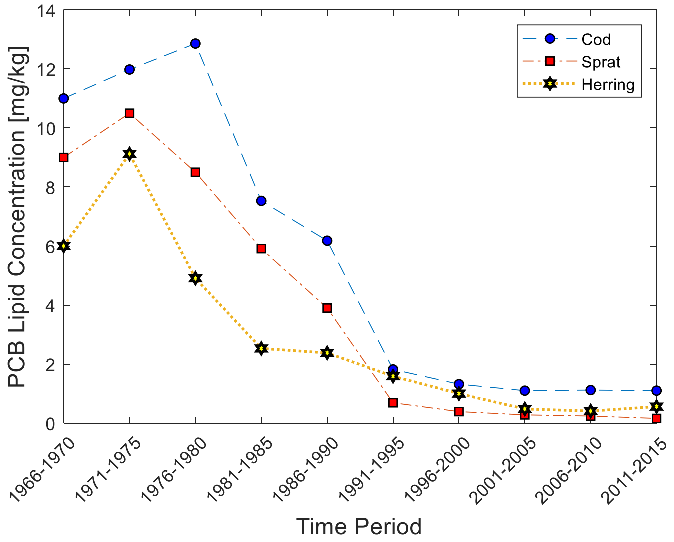

**Fig. 2** Temporal sample mean PCB lipid concentrations in different species from the Baltic Sea during 1966-2015, according to data compiled by Segerstedt (2019). Three species of Baltic fish used as prey by grey seals; cod (*Gadus morhua*), sprat (*Sprattus sprattus*) and herring (*Clupea harengus*).

#### 2.1 Baltic grey seal biology and ecology

The TK model of temporal PCB dynamics incorporates aspects of grey seal life history, feeding habits, and growth as these have important effects on tissue PCB concentrations. The parameters defining the biology and ecology of grey seals in the model are detailed in Table 2 and dietary preference for different seal age classes in Table 3.

##### 2.1.1. Grey seal growth and life history parameters

In the UK, adult grey seal males on average have a weight of 233 kg and a length of 200 cm, whereas mature females on average have a weight of 155 kg and a length of 180 cm (Härkönen 2016). Females mature at an age of 3–5 years whereas males mature at around 6 years, although not socially mature until an age of 8 years (Jefferson et al. 2008). The maximum life span for Baltic grey seals is about 35-45 years, though only a minority of the seals become older than 24 years (HELCOM 2018). Baltic grey seals breed and mate in February to March (Jüssi et al. 2008). The fertilized egg is implanted in the uterus after a *delay period* of about 100 days. The succeeding *gestation period* of embryonic and foetal growth to parturition lasts for about 240 days. Mature females give birth to at most one pup a year (Bergman 2007). The new-born pup has a weight of about 12 kg (Härkönen 2016) and a length of 60-100 cm (Bergman 2007). The pup is nursed for 15-18 days and weaning is abrupt (Härkönen 2016). During the lactation period, pups daily consume about 4 kg of breast milk (Iverson 1993). The milk contains about 52 % fat. At weaning, the pup has increased its weight to 37-48 kg, whereas the mother's weight loss is massive (Härkönen 2016). Grey seals breed on pack ice formations if they are available and otherwise on land (Jüssi et al. 2008). During two weeks in May and June, hundreds of seals gather at common sites to moult, without feeding (Hansson et al. 2017).

**Table 2**. Life history and growth parameters

| parameter | definition | value | unit | source |
|---|---|---|---|---|
| $\Delta\tau_l$ | Lactation period length | 18 | d | Härkönen (2016) |
| $\Delta\tau_d$ | Delay period length | 100 | d | Bergman (2007) |
| $\Delta\tau_g$ | Gestation period length | 247 | d | Calculation |
| $\tau_{SA}$ | Sub adult age | 1 | yr | Harding, et al (2007) |
| $\tau_{mat}$ | Sexual maturation age | 5 | yr | Jefferson, et al.(2008) |
| $W_{00}$ | Initial foetal weight | $3.0 \times 10^{-7}$ | kg | Calculation |
| $W_{birth}$ | Birth weight | 12 | kg | Härkönen (2016) |
| $W_{wean}$ | Weaning weight | 43 | kg | Härkönen (2016) |
| $W_{mat}$ | Mature body weight | 153 | kg | Estimation |
| $W_{max}$ | Maximum body weight | 160 | kg | Härkönen (2016) |
| $W_\infty$ | Asymptotic adult body weight | 180 | kg | Estimation |
| $W_{lac,6}$ | Primiparous lactation body weight | 105 | kg | Calculation |
| $W_{lac,7}$ | Multiparous lactation body weight | 110 | kg | Estimation |
| $\alpha$ | Body shape factor | 27 | kg/m$^3$ | Calculation |
| $L_{\max}$ | Maximum body length | 180 | cm | Härkönen (2016) |
| $g_0$ | Exponential growth rate constant for fetus and pups | 26 | year$^{-1}$ | Calculation |
| $g_L$ | Lactation body weight decline rate constant | 7.6 | year$^{-1}$ | Calculation |
| $k_6$ | Primiparous body weight regain rate constant | 201 | kg/year | Calculation |
| $k_7$ | Multiparous body weight regain rate constant | 183 | kg/year | Calculation |
| $\gamma$ | von Bertalanffy growth rate constant | 0.49 | year$^{-1}$ | Calculation |

Body growth parameters (Table 2) were estimated based on body size data for Baltic grey seals. Foetuses are assumed to grow exponentially during the gestation period:

$$W_0(\hat{\tau}) = W_{00} e^{g_0(\hat{\tau}-(\Delta\tau_l+\Delta\tau_d))}, \qquad \Delta\tau_l + \Delta\tau_d \le \hat{\tau} \le 1 \tag{1}$$

At the end of the gestation period, the fetus has reached the birth weight; $W_0(1) = W_{birth}$. Hence, the initial foetal weight $W_{00}$ is calculated as:

$$W_{00} = W_{birth} e^{-g_0 \Delta\tau_g} \tag{2}$$

New-born pups continue to grow exponentially, with the same growth rate as fetuses, during the lactation period:

$$W_1(\hat{\tau}) = W_{00} e^{g_0(\Delta\tau_g+\hat{\tau})}, \quad 0 \le \hat{\tau} < \Delta\tau_l \tag{3}$$

At the end of the lactation period, pups reach weaning weight; $W_1(\Delta\tau_l) = W_{wean}$. Hence, the exponential growth rate constant $g_0$ is calculated as:

$$g_0 = ln(W_{wean}/W_{birth})/\Delta\tau_l \tag{4}$$

Pups keep weaning weight until they reach sub adult age $\tau_{SA} = 1$ year.

Grey seal females grow according to von Bertalanffy growth function from weaning ($\tau = \Delta\tau_l$) until the age of sexual maturation ($\tau_{mat} = 5$ years), reaching mature body length $L_{mat}$. The body length $L(\tau)$ at age $\tau$ for non-mature females is:

$$L(\tau) = L_\infty - (L_\infty - L_{wean})e^{-\gamma(\tau-\tau_{SA})}, \qquad \tau_{SA} \le \tau \le \tau_{mat} \tag{5}$$

Here, $L_\infty$ is the asymptotic adult body length, $L_0$ is the initial length (the length at $\tau = \tau_0$) and $\gamma$ is the growth rate constant. In accordance with standard DEB theory, isometric growth and a body weight proportional to the structural body volume are assumed (Kooijman 2000):

$$W(\tau) = \alpha L(\tau)^3 \tag{6}$$

With specified maximum body length and weight ($L_{max}$ and $W_{max}$), based on data for full-grown Baltic grey seal females, the body shape constant $\alpha$ is calculated as:

$$\alpha = W_{max}/L_{max}^3 \tag{7}$$

The asymptotic body length $L_\infty$ and the weaning body length $L_{wean}$ are obtained from specified values of the asymptotic body weight $W_\infty$ and the weaning body weight $W_{wean}$:

$$L_\infty = (W_\infty/\alpha)^{1/3}\ , \qquad L_{wean} = (W_{wean}/\alpha)^{1/3} \tag{8}$$

At sexual maturation, females reach mature body weight $W_{mat}$, assumed to be somewhat lower than maximum body weight $W_{max}$, reached by multiparous females just before they start to lactate. Females decrease weight exponentially throughout the lactation period, from the initial body weight ($W_{mat}$ or $W_{max}$) to the lactation body weight; $W_{lac,6}$ or $W_{lac,7}$ (for primiparous and multiparous females, respectively). During the delay period, females linearly regain the weight loosed during lactation. During the gestation period, females continue to growth according to von Bertalanffy and reach maximum body weight $W_{max}$ at the end of the year. Succeeding years, females repeat the pattern of exponential decline, linear regain and von Bertalanffy growth. The von Bertalanffy growth rate constant $\gamma$ is calculated as:

$$\gamma = \frac{ln[(L_\infty - L_{wean})/(L_\infty - L_{max})]}{\tau_{mat} - \tau_{SA} + \Delta\tau_g} \tag{9}$$

The mature body length and weight ($L_{mat}$ and $W_{mat}$) are calculated as:

$$L_{mat} = L_\infty - (L_\infty - L_{wean})e^{-\gamma(\tau_{mat}-\tau_{SA})} \quad , \quad W_{mat} = \alpha L_{mat}^3 \tag{10}$$

The body length of females of age class $i$ at time $\hat{\tau}$ during the growth period of a year is obtained from von Bertalanffy growth function:

$$\begin{aligned} L_i(\hat{\tau}) &= L_\infty - (L_\infty - L_{wean})e^{-\gamma(i-1+\hat{\tau}-\tau_{SA})} \quad i = 2,\dots,5 \quad 0 \le \hat{\tau} \le 1 \\ L_i(\hat{\tau}) &= L_\infty - (L_\infty - L_{mat})e^{-\gamma[\hat{\tau}-(\Delta\tau_l+\Delta\tau_d)]} \quad i = 6,\dots,46 \quad \Delta\tau_l + \Delta\tau_d \le \hat{\tau} \le 1 \end{aligned} \tag{11}$$

Corresponding body weights are obtained as:

$$W_i(\hat{\tau}) = \alpha[L_i(\hat{\tau})]^3 \quad , \quad i = 1,\dots,46 \quad , \quad 0 \le \hat{\tau} \le 1 \tag{12}$$

The lactation body weight $W_{lac,7}$ for multiparous females is specified from data, whereas the corresponding body weight for primiparous females is calculated as:

$$W_{lac,6} = \frac{W_{mat}}{W_{max}} W_{lac,7} \tag{13}$$

The exponentially declining body weight of females at time $\hat{\tau}$ during lactation is obtained as:

$$\left.\begin{aligned} W_6(\hat{\tau}) &= W_{mat}e^{-g_L\hat{\tau}} \\ W_i(\hat{\tau}) &= W_{max}e^{-g_L\hat{\tau}} \quad (i = 7,\dots,46) \end{aligned}\right\} 0 \le \hat{\tau} < \Delta\tau_l \tag{14}$$

The exponential body weight decline rate constant is calculated as:

$$g_L = ln\left(W_{max}/W_{lac,M}\right)/\Delta\tau_l \tag{15}$$

The linearly increasing body weights during the delay period are obtained as.

$$\left.\begin{array}{l} W_6(\hat{\tau}) = W_{lac,6} + k_6(\hat{\tau} - \Delta\tau_l) \\ W_i(\hat{\tau}) = W_{lac,7} + k_7(\hat{\tau} - \Delta\tau_l) \quad (i = 7, \dots, 46) \end{array}\right\} \quad \Delta\tau_l \le \hat{\tau} < \Delta\tau_l + \Delta\tau_d \tag{16}$$

The regain rate constants $k_6$ and $k_7$ (for primiparous and multiparous, respectively) are calculated as:

$$k_6 = \left(W_{mat}/W_{lac,6}\right)/\Delta\tau_d \quad , \quad k_7 = \left(W_{max}/W_{lac,7}\right)/\Delta\tau_d \tag{17}$$

##### 2.1.2. Grey seal diet parameters

Hansson, et al. (2017), estimated the daily mean prey consumption of Baltic grey seals to 4.5-5.0 kg, corresponding to an annual per capita consumption of 1750 kg. It was here assumed that the largest females (with a weight of $W_{\max} = 160$ kg) have a yearly fish consumption of $\overline{W}_{\max}^{\text{fish}} = 1800$ kg. According to Hansson, et al. (2017), 80 % of the Baltic fish biomass is constituted by sprat (*Sprattus sprattus*), herring (*Clupea harengus*) and cod (*Gadus morhua*). Our model assumes that Baltic grey seals feed exclusively on these three species. Prey preference indices $\phi_{ij}$, describing the preference of prey $j$ by age class $i$, were calculated from published mean fractions of total prey biomass ($\Phi_{ij}$) found in analyses of gut content in Baltic grey seals (Table 3):

$$\phi_{ij} = \Phi_{ij}/\sum \Phi_{ij} \qquad i = 1, \dots, 46 \quad , \quad j = 1, 2, 3 \tag{18}$$

The mean body lipid indices $\bar{\rho}_{P,j}$ for different prey species $j$ were obtained based on their fat content according to Fiskbasen (1996).

**Table 3**. Dietary parameters for seals of different age classes. Estimated fraction of total prey biomass for different species consumed by Baltic grey seals and corresponding prey preference indices. Estimations of total prey biomass consumed by Baltic grey seals are based on gut analyses of Baltic grey seal males and females (Lundström et al. 2010) and females (Tverin et al. 2019). $\Phi_{ij}$: Fraction of total prey biomass. $\phi_{ij}$: Prey preference index. $\bar{\rho}_{P,j}$: Mean body lipid for prey species (Fiskbasen 1996).

| prey | prey index (*j*) | pups | | subadults | | adults | | $\bar{\rho}_{P,j}$ |
|---|---|---|---|---|---|---|---|---|
| | | $\Phi_{ij}$ | $\phi_{ij}$ | $\Phi_{ij}$ | $\phi_{ij}$ | $\Phi_{ij}$ | $\phi_{ij}$ | |
| herring | 1 | 59 % | 0.72 | 54 % | 0.78 | 42 % | 0.98 | 7.5 % |
| sprat | 2 | 20 % | 0.24 | 11 % | 0.16 | 0 % | 0 | 15 % |
| cod | 3 | 3 % | 0.04 | 4 % | 0.06 | 1 % | 0.02 | 1.0 % |

##### 2.1.3. Grey seal population parameters

Ideal vital rates ($F_i^0$ and $P_i^0$) for a population with maximal possible reproduction were used as background values, representing fertilities and survival rates in absence of PCB exposure. These were obtained from Harding et al (2007), who reviewed life history data for grey seals from different parts of the world, derived vital rates for Baltic grey seals and established a Leslie matrix model, generating the observed population growth rate of $\lambda = 1.075$. They also derived optimal vital rates and calculated a maximum possible population growth rate of $\lambda = 1.10$. Values of vital rates are presented in Table 4. Notice that females younger than six years ($i = 1 - 5$) do not reproduce and that the youngest mature females ($i = 6$) have a fertility half that of adult females ($i = 7, \dots, 45$). In the ideal population, adult females obtain almost one pup a year, half of them female pups.

**Table 4**. Age-specific fertilities and survival rates for Baltic grey seals. Realistic values ($F_i$, $P_i$) are values adopted in a population assessment by Harding, et al. (2007). Ideal values ($F_i^0$, $P_i^0$) correspond to a population with maximal possible reproduction (Harding et al. 2007). The formers are used as background values in the TKTD population model.

| **age class ($i$)** | **fertility ($F_i$)** | **survival ($P_i$)** | **fertility ($F_i^0$)** | **survival ($P_i^0$)** |
|---|---|---|---|---|
| 1 | 0 | 0.700 | 0 | 0.700 |
| 2-4 | 0 | 0.932 | 0 | 0.932 |
| 5 | 0 | 0.950 | 0 | 0.950 |
| 6 | $0.1875 \cdot P_6$ | 0.950 | 0.24 | 0.950 |
| 7-45 | $0.375 \cdot P_i$ | 0.950 | 0.48 | 0.950 |
| 46 | 0 | 0 | 0 | 0 |

Since seals are long-lived animals with high adult survival and low fecundity, populations typically consist of a large proportion of adults and the growth rate is low. This makes seal populations relatively resistant to short-term changes in the environment and the population dynamics are primarily governed by competition, predation, disease and availability of prey (Svensson et al. 2010). For long-lived K-strategic species, such as seals, density dependence is an important factor for long-term population dynamics (Kauhala et al. 2014). Seal populations show an age-specific response to increased population density with a rapid decrease in juvenile survival, decreased fecundity and a relatively constant adult survival, changing the age composition towards higher proportions of adults (Svensson et al. 2010). When a population approaches carrying capacity, the mortality is increased foremost among pups and sub-adults of 1-3 years. Hence, the most variable vital parameter in a seal population is pup survival. Pregnancy rates are lowest in the youngest females (4-5 years old), has a maximum during 6-24 years and then start to decline. However, females of 40 years may reproduce (Kauhala et al. 2014). The effects of population density on survival and fertility of grey seals is modelled using age-specific survival and fertility density effect factors (Table 5). The carrying capacity was put to 100 000, since the Baltic grey seal population likely approached this size in the early 1900s (Harding and Härkönen 1999). It was assumed that adult survival and fertility are only dependent on the number of adults, not on the number of pups ($p_{i,1} = f_{i,1} = 0\ ,\ i = 2, \dots, 46$). Since pups are more sensitive to harsh conditions than older animals, it was also assumed that pup survival is more affected by population density than other transition rates ($p_{1,j} > p_{i,j}\ ,\ p_{1,j} > f_{i,j}\ ,\ i,j = 2, \dots, 46$). Moreover, it was assumed that $p_{1,j} = p_{1,1}$ ($j = 2, \dots, 46$), $p_{1,1} = 2p_{2,2}$, $p_{i,j} = f_{i,j} = p_{2,2}$ ($i,j = 2, \dots, 46$). The adult density factor $p_{2,2}$ was adjusted such that the total population (females and males) approaches carrying capacity after long time under ideal conditions (no adverse effects from contaminants), yielding $p_{2,2} = 2 \cdot 10^{-6}$.

Environmental stochasticity is also included to affect survival and fecundity through fluctuation factors (Table 5). It was assumed that environmental stochasticity poses small fluctuations in survival rates and five times as large fluctuations in fertilities.

**Table 5**. Population and environmental stochasticity parameters. $p_{ij}$: Survival density effect (Effect of an individual in age class $j$ on the survival of an individual in age class $i$).
$f_{ij}$: Fertility density effect (Effect of an individual in age class $j$ on the fertility of an individual in age class $i$).

| **parameter** | **definition** | **value** |
|---|---|---|
| $p_{1,1-46}$ | Survival density effect of all seals on pups | $4 \times 10^{-6}$ |
| $p_{2-46,1}$ | Survival density effect of pups on subadults and adults | 0 |
| $p_{2-46,2-46}$ | Survival density effect of subadults and adults on subadults and adults | $2 \times 10^{-6}$ |
| $f_{1-46,1}$ | Fertility density effect of pups on all seals | 0 |
| $f_{1-5,1-46}$ | Fertility density effect of all seals on pups and subadults | 0 |
| $f_{6-46,2-46}$ | Fertility density effect of subadults and adults on adults | $2 \times 10^{-6}$ |
| $z_P$ | Survival fluctuation factor | 0.02 |
| $z_F$ | Fertility fluctuation factor | 0.10 |

#### 2.2 Toxicokinetics

Model parameters used in the TK model to describe temporal dynamics of PCB concentrations in seals throughout their lifespan are presented in Table 6. The TK parameters define the rate constants that regulate PCB accumulation and depuration in grey seals, covering vertical transfer processes (placental and lactational transfer), dietary uptake, metabolic transformation, fecal egestion, and growth dilution. The derived parameter values are described below.

##### 2.2.1 Vertical transfer rate constants

To estimate lactation rate constants ($k_{L,i}$) and placental transfer rate constants ($k_{P,i}$), published data on milk transfer in lactating Canadian grey seals (Lang et al. 2011) was combined with published data on PCB transfer through breast milk in Scottish grey seals (Berghe et al. 2012). Lang, et al. (2011) presented body mass, body composition (fractions of water, protein and fat), daily milk output and milk composition for females and their pups at early and late lactation. A distinction was made between *primiparous females* (first-time nurses) and *multiparous females* (experienced nurses). Primiparous females had lower daily milk production, shorter lactation periods and smaller pups at weaning, compared to multiparous females. Berghe, et al. (2012) presented data on PCB concentrations on a lipid-weight basis for female-pup pairs at early and late lactation, including PCB levels in maternal outer and inner blubber, maternal blood, breast milk and pup blood. PCB concentrations increased significantly in all analysed tissues and milk between early and late lactation. In order to estimate lactation rate constants from mentioned publications, it was assumed that the total PCB body burden in females decreases exponentially during lactation:

$$W_i^{PCB}(\tau) \approx W_i^{PCB}(0)e^{-k_{L,i}\tau}, \qquad i = 6, \dots, 46 \tag{19}$$

Here, $W_i^{PCB}(\tau)$ is the total PCB body burden of females at time $\tau$ during the lactation period ($0 \leq \tau \leq \Delta\tau_l$). Lactation rate constants were estimated as:

$$k_{L,i} \approx \frac{\ln\left[W_i^{PCB}(\tau_{early})/W_i^{PCB}(\tau_{late})\right]}{\tau_{late} - \tau_{early}}, \qquad i = 6, \dots, 46 \tag{20}$$

In order to estimate total PCB body burdens $W_i^{PCB}(\tau)$ at early lactation ($\tau = \tau_{early}$) and late lactation ($\tau = \tau_{late}$) from published data, the following assumptions were made:

1. All fat in a female body is either *outer blubber* or *inner blubber* and both layers are reduced during lactation.
2. The total amount of PCB in the outer blubber layer is constant throughout lactation. PCB concentration change is exclusively an effect of reduced blubber mass.
3. The inner blubber layer reduces linearly with time throughout lactation.
4. All inner blubber is consumed at the end of lactation.
5. The total PCB body burden of a female is reduced by the same amount as is transferred to her pup through breast milk during lactation.
6. Milk PCB level increases linearly with time throughout lactation.
7. Lost female body mass that is not blubber contains the PCB required to obtain the total PCB body burden according to assumption 5.

We also assumed that about 10 % of a seal body is constituted by bone, the blood volume of grey seals is about 213 ml/kg and mammals have a blood density of 1060 kg/m$^3$ (Castellini and Mellish 2016). A blood mass fraction of 22.6 % for grey seals was then estimated. Assumptions 1-4 were used in calculations of outer/inner blubber ratios and assumptions 5-7 in calculations of total PCB body burdens.

The lactation rate constant for primiparous females ($k_{L,6} = 4.36/\text{year}$) corresponds to a 19 % reduction of the total PCB body burden during lactation (18 days), whereas the lactation rate constant for multiparous females ($k_{L,i} = 5.37/\text{year}$) corresponds to a 23 % reduction. As a comparison, Addison (1977) proposed that female grey seals lose about 15 % of their total PCB body burden during the nursing period.

In order to estimate placental transfer rate constants from mentioned publications, it was assumed that that the total PCB body burden in females during gestation decreases exponentially under constant body weight:

$$c_i(\tau) \approx c_i(\Delta\tau_l + \Delta\tau_d)e^{-k_{P,i}[\tau-(\Delta\tau_l+\Delta\tau_d)]}, \qquad i = 6, \dots, 46 \tag{21}$$

Here, $c_i(\tau)$ are total body PCB concentrations in females at time $\tau$ during the gestation period ($\Delta\tau_l + \Delta\tau_d \leq \tau \leq 1$). Hence, placental transfer rate constants were estimated as:

$$k_{P,i} \approx \ln[c_i(\Delta\tau_l + \Delta\tau_d)/c_i(1)]\,/\Delta\tau_g \quad , \qquad i = 6, \dots, 46 \tag{22}$$

In order to estimate PCB concentrations $c_i(\tau)$ before gestation ($\tau = \Delta\tau_l + \Delta\tau_d$) and after gestation ($\tau = 1$) from published data, the following assumptions were made:

1. Blood PCB levels increase linearly in females and pups during lactation.
2. Pup body and blubber weights increase exponentially from birth to late lactation.
3. The female outer blubber layer has the same weight at parturition and early lactation.
4. The blood/blubber PCB concentration ratio is the same for females and pups at parturition.
5. All PCB in females and pups at parturition is located in blubber.
6. The reduction of PCB body burden in females during gestation equals the total PCB body burden in pups at birth (dietary uptake and metabolic transformation during gestation are neglected).

The estimated value of $k_{P,i} = 0.07/\text{year}$ ($i = 6, \dots, 46$) corresponds to a 4 % loss of the total PCB body burden in females during gestation (247 days), which can be compared to the 19-23 % loss during lactation (18 days). This agrees with the general view that vertical PCB transfer in marine mammals is dominated by lactation, with only a minor contribution from placental transfer (Subramanian et al. 1988).

##### 2.2.2. Assimilation efficiencies

In their bioaccumulation model for arctic ringed seals, Hickie, et al. (2005) adopted an assimilation efficiency of 90 %, both for diet and breast milk. The same values were here adopted for Baltic grey seals:

$$\varphi_{D,i} = 0.90\,, \quad i = 1, \dots, 46 \tag{23}$$

$$\varphi_L = 0.90 \tag{24}$$

Here, $\varphi_{D,i}$ are the diet assimilation efficiencies and $\varphi_L$ is the lactation assimilation efficiency.

##### 2.2.3. Removal rate constants

The removal rate constants $k_{R,i} = k_{M,i} + k_{F,i}$ account for metabolic transformation and fecal egestion. Since fecal egestion is accounted for via assimilation efficiencies for dietary uptake and lactation, the fecal egestion rate constants were neglected ($k_{F,i} = 0$) and the removal rate constants obtained as:

$$k_{R,i} = k_{M,i}, \qquad i = 1, \dots, 46 \tag{25}$$

According to Klanjscek et al. (2007), metabolic transformation of PCB in marine mammals is negligible. However, it was here assumed that a small fraction is removed through metabolic transformation. In their bioaccumulation model for artic ringed seals, Hickie, et al. (2005) estimated elimination rate constants for 20 different PCB congeners. Linear regression was used to fit model outputs with observed contaminant concentrations in seals. The estimated elimination rate constants ranged from 0.03/year (PCB 153) to 2.5/year (PCB 18), with 0.17/year for Total PCB. These estimates were here considered as metabolic transformation rate constants. It is also assumed that metabolic transformation rates scale between seal species according to Kleiber's law (Kleiber 1962), stating that metabolic rate MR (rate of energy spent per unit time) of any animal increases with body weight $W$ according to:

$$R = aW^{3/4} \tag{26}$$

The proportionality constant $a$ is assumed to be similar across different species of mammals. Larger animals thus metabolize PCB at a higher total rate than smaller animals, but smaller animals metabolize PCB faster per unit of body mass. The specific PCB metabolic transformation rate is assumed to be proportional to the specific metabolic rate:

$$\frac{\dot{W}_{PCB}}{W_{PCB}} \propto \frac{R}{W} \propto W^{-1/4} \tag{27}$$

It thus follows that the metabolic transformation rate constant $k_M$ scales with body mass $W$ as:

$$k_M \propto W^{-1/4} \tag{28}$$

Adult ringed seals have a body weight of 50-70 kg (Jefferson et al. 2008). Assuming a mean body weight of $W_{RS} = 60$ kg, metabolic transformation rate constants for ringed seals ($k_{M,RS}$) are transformed to metabolic transformation rate constants for grey seals ($k_{M,i}$) of different age classes $i$ according to:

$$k_{M,i} = \left[\frac{\overline{W}_i^j}{W_{RS}}\right]^{-1/4} k_{M,RS} \quad , \quad i = 0, \dots, 46 \tag{29}$$

Here, $\overline{W}_i^j$ is the mean the body weight for age class $i$ during time period $j$ (where $i$ = 0 represents fetuses). The metabolic transformation rate constant for Total PCB in arctic ringed seals $k_{M,RS} = 0.17/\text{year}$ (Hickie et al. 2005) was adopted to calculate values of metabolic transformation rate constants for Baltic grey seals of different age classes, ranging from 0.1/year (pregnant females) to 0.5/year (fetuses).

**Table 6**. TK model parameters and rate constants.

| variable/ parameter | definition | value | unit |
|---|---|---|---|
| $\bar{c}_{D,i}$ | PCB concentration in the diet of seals in age class $i$ during year $t$ | - | mg/kg |
| $\bar{k}^{j}_{D,i}$ | PCB dietary uptake rate constant | $\kappa\varphi_{D,i}[W_i(1)]^{2/3}/\bar{W}^{j}_{i}$ | $\text{year}^{-1}$ |
| $\hat{k}^{j}_{E,i}$ | PCB total elimination rate | $l$: $k_{L,i}+\bar{k}^{l}_{G,i}$<br>$d$: $k_{R,i}+\bar{k}^{d}_{G,i}$<br>$g$: $k_{R,i}+k_{P,i}+\bar{k}^{g}_{G,i}$ | $\text{year}^{-1}$ |
| $k_{R,i}$ | PCB removal rate constant | $k_{M,i}+k_{F,i}$ | $\text{year}^{-1}$ |
| $k_{M,i}$ | PCB metabolic transformation rate constant | 0.1-0.5 | $\text{year}^{-1}$ |
| $k_{F,i}$ | PCB fecal egestion rate constant | 0 | $\text{year}^{-1}$ |
| $k_{L,i}$ | PCB lactation rate constant | $i$ = 6: 4.36<br>$i$ > 6: 5.37 | $\text{year}^{-1}$ |
| $k_{P,i}$ | PCB placental transfer rate constants | 0.07 | $\text{year}^{-1}$ |
| $\bar{k}^{g}_{G,i}$ | Mean growth dilution rate | - | - |
| $k_{R,i}$ | | - | - |
| $\bar{W}^{fish}_{max}$ | Maximum yearly fish consumption | 1800 | kg |
| $\hat{\rho}$ | Body lipid index for seals | 30 | % |
| $\varphi_{D,i}$ | Diet assimilation efficiencies | 0.90 | - |
| $\varphi_{L}$ | Lactation assimilation efficiency | 0.90 | - |
| $\alpha_D$ | prey consumption factor | 66 | $\text{kg}^{1/3}\cdot\text{year}^{-1}$ |

#### 2.3 Toxicodynamics

Model parameters used in the TD model to describe temporal dynamics of PCB effects in seals throughout their lifespan are presented in Table 7. The TD parameters define the rate constants, threshold levels, and tolerance levels that regulate the impact of PCB concentrations on hazard and stress in grey seals of different age classes. The derived parameter values are described below.

The toxicodynamic model includes three different set of parameters; *reproductive stress* parameters, *foetal hazard* parameters and hazard parameters related to *survival* of pups and adults. Each parameter set includes three type of constants; *stress/killing rate constants*, *damage threshold levels* and *recovery rate constants*. With respect to the many empirical observations of reproductive organ lesions in Baltic grey seals during the 1970s and the 1980s, stress to the reproductive apparatus is probably the most crucial path in which PCB affects vital rates. The corresponding reproductive stress parameters were estimated by a calibration procedure where pregnancy rates predicted by the TKTD population model were compared to reported pregnancy rates (Roos et al. 2012). For the other (less crucial) toxicodynamic parameters, preliminary values were chosen based on the reproductive stress parameters and on DEBtox parameters from minks (Desforges et al. 2017).

##### 2.3.1 Calibration of Reproductive Stress Parameters

There are a number of publications, reporting historical data on pregnancy rates and prevalence of reproductive organ lesions associated with sterility (uterine occlusions, uterine stenosis and uterine leiomyoma) in Baltic grey seal females (Bergman 1999, Bäcklin et al. 2003, Bergman 2007, Bredhult et al. 2008, Roos et al. 2012, Bäcklin and Moraeus 2013). Roos, et al. (2012) analysed the reproductive outcome of Baltic grey seals in relation to organochlorine concentrations during the time period 1966-2015. Frequency of uterine obstructions (stenosis or occlusions) and uterine tumours were investigated in 277 adult females, whereas pregnancy rates were investigated in 144 adult females (five years or older), collected between 1973 and 2010. Blubber from 292 juvenile grey seals (0-3 years), collected between 1968 and 2010, where used to measure PCB levels on a lipid weight basis. To illustrate general patterns, Roos, et al. (2012) fitted curves of different forms to temporal trends for reproduction indices and Total PCB blubber concentrations. They used non-linear regression to fit the following logistic function to the temporal trend of mean pregnancy rates in Baltic grey seal females (age class $i = 6, \dots 46$) at year $t$:

$$\mathrm{PR}(t) = \frac{\exp[-3.83 + 0.24(t - 1975)]}{1 + \exp[-3.83 + 0.24(t - 1975)]} \quad (30)$$

We compared this curve to pregnancy rates predicted by the TKTD population model, using temporal data on PCB concentrations in prey (Segerstedt 2019) as model input. It was assumed that reduced pregnancy is an effect of reproductive stress, caused by damage to the reproductive apparatus. In order to obtain the mean pregnancy rate ($PR$) from the TKTD population model, the mean fertility rate for age class 6-46 at the end of the delay period were multiplied by 2 (to include offspring of both genders):

$$PR(t) = 2\bar{F}^d_{6-46}(t) \quad (31)$$

Notice that the fertility rate at the end of the year was not used in the calculation, since this fertility also account for reproductive failure due to foetal hazard in pregnant females. The mean fertility for females of age class 6-46 at the end of the delay period year $t + 1$ is obtained as:

$$\bar{F}^d_{6-46}(t+1) = \frac{\sum_{i=6}^{46} N_i(t) F_i^d(t+1)}{\sum_{i=6}^{46} N_i(t)} \quad (32)$$

Here, $N_i(t)$ is the number of females in age class $i$ at the end of year $t$. Pregnancy rates, derived from the TKTD population model, were compared to observed pregnancy rates by visual inspection of graphs (Fig. 3). Reproductive stress parameters were adjusted to fit model-generated curves to the empirical curve.

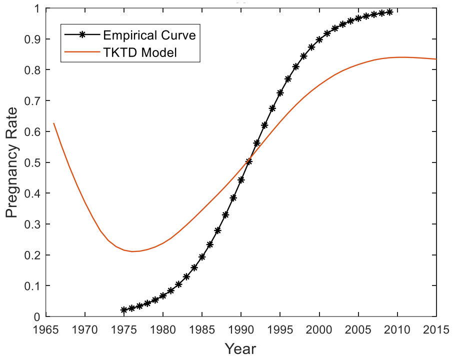

**Fig. 3** Pregnancy rates for the Baltic grey seal population between 1966 and 2015 according to empirical curve from Roos, et al. (2012) and mean pregnancy rates for females of ages 5-46 years according to the deterministic TKTD population model. For each model variant, the recovery rate constant $\tilde{r}_i$ was chosen to fit model curves to the empirical curve.

##### 2.3.2. Estimation of Toxicodynamic Model Parameters Related to Survival

The stress rate constant with respect to fetal hazard $\sigma_0$ was translated from DEBtox parameters for embryonal survival of minks (Desforges et al. 2017), see below (section 2.3.3). The corresponding damage threshold level $d_{T,0}$ was chosen arbitrarily as ten times smaller than the damage threshold level for reproductive stress in adults since younger age classes are known to be more sensitive. The recovery rate constant for foetal hazard $r_0$ was then scaled from the reproductive stress constants:

$$r_0 = \frac{\sigma_0}{\tilde{\sigma}_{2-46}} \tilde{r}_{2-46} \quad (33)$$

The killing rate constant with respect to pup survival $\sigma_1$ was translated from DEBtox parameters for kit survival of minks (Desforges et al. 2017), see below (section 2.3.3). The same killing rate constant was chosen for adults; $\sigma_{2-46} = \sigma_1$. The damage threshold level $d_{T,2-46}$ was chosen to obtain good agreement between model and historical data for population size.

The corresponding damage threshold level for pups was set to half of this value; $d_{T,1} = 0.5 d_{T,2-46}$. The recovery rate constants with respect to pup and adult survival were scaled from reproductive stress constants:

$$r_i = \frac{\sigma_i}{\tilde{\sigma}_{2-46}} \tilde{r}_{2-46} \quad (34)$$

**Table 7**. Toxicodynamic parameters. Mink TDM parameters were converted from DEBtox parameters (Desforges et al. 2017). Effects on *reproduction* include *embryo/fetus survival* and *reproduction costs*. Effects on *survival* refer to *kit/pup survival*.

| parameter | definition | value | unit |
|---|---|---|---|
| $\sigma_i$ | Killing rate constant (pups and adults) | 0.0073 | $\frac{\text{kg}}{\text{mg}} \cdot \text{year}^{-1}$ |
| $r_i$ | Recovery rate constant (survival) | 0.12 | $\text{year}^{-1}$ |
| $d_{T,i}$ | Damage threshold levels (survival) | $i = 1$: 0.10<br>$i > 1$: 0.20 | - |
| $\tilde{\sigma}_i$ | Stress rate constant (reproductive stress) | $i = 1$: 0<br>$i > 1$: 1 | $\frac{\text{kg}}{\text{mg}} \cdot \text{year}^{-1}$ |
| $\tilde{r}_i$ | Recovery rate constants (reproductive stress) | 16 | $\text{year}^{-1}$ |
| $\tilde{d}_{T,i}$ | Damage threshold levels (reproductive stress) | 0.010 | - |
| $\sigma_0$ | Killing rate constant (fetus) | 0.023 | $\frac{\text{kg}}{\text{mg}} \cdot \text{year}^{-1}$ |
| $r_0$ | Recovery rate constant (fetus) | 0.0010 | $\text{year}^{-1}$ |
| $d_{T,0}$ | Damage threshold level (fetus) | 0.37 | - |
| $\text{NEC}_S$ | DEBtox no effect concentration for stress on embryo survival and reproductive stress in mink | 0.017-0.65 | mg/kg |
| $c_T$ | DEBtox stress tolerance for embryo survival and reproductive stress in mink | 0.69-2.25 | mg/kg |
| $\text{NEC}_H$ | DEBtox no effect concentrations for hazard related to kit survival in mink | 2.94-3.71 | mg/kg |
| $c_{Th}$ | DEBtox hazard tolerance for kit survival in mink | $9.12 \text{ x}10^{-6}$ – $1.08 \text{ x}10^{-5}$ | mg/kg |
| $\tilde{d}_T$ | Damage threshold level for reproduction | mink: 0.0076-0.94<br>seal: 0.018–2.2 | - |
| $\tilde{\sigma}$ | Stress rate constant for reproduction | mink: 0.011-0.035<br>seal: 0.011-0.035 | $(\text{kg/mg}) \cdot \text{year}^{-1}$ |
| $\tilde{r}_{max}$ | Recovery rate constant limit for reproduction | mink: 0.025-3.15<br>seal: 0.0011-1.3 | $\text{year}^{-1}$ |
| $d_T$ | Damage threshold level for survival | mink: $2.\ 5 \cdot 10^{-6}$ - $3.7 \cdot 10^{-6}$<br>seal: $5.8 \text{ x}10^{-6}$ – $8.7 \text{ x}10^{-6}$ | - |
| $\sigma$ | Killing rate constant for survival | mink: 0.0064 – 0.0081<br>seal:0.0064-0.0081 | $(\text{kg/mg}) \cdot \text{year}^{-1}$ |
| $r_{max}$ | Recovery rate constant limit for survival | mink: 6482 – 9686<br>seal: 2741-4097 | $\text{year}^{-1}$ |

##### 2.3.3. Transformation of DEBtox parameters

The toxicodynamic model is a threshold damage model (TDM), including stress rate constants, recovery rate constants and damage threshold levels. It is not a DEBtox model and does not use stress tolerances and no effect concentrations as parameters. Here, a rough and simple way to transform DEBtox parameters for one mammal into TDM parameters for another mammal is suggested. However, the method can only be used as a way to obtain preliminary values before a more accurate parametrization is performed. As an example, DEBtox parameters for mink are transformed into TDM parameters for grey seal. Some of the transformed values were used as estimations of toxicodynamic model parameters in the TKTD population model for Baltic grey seals.

Desforges, et al. (2017) developed a DEBtox model for mink, parameterized from published data on growth and reproduction in captive minks fed with PCB contaminated fish. Embryos were exposed to PCB through the placenta throughout the gestation period ($\Delta\tau_g^{mink} = 42$ days). Physical modes of action (PMoAs) were identified by comparing dose-response-curves from the model and reported data. Tolerance parameters and no effect concentrations were adjusted to fit model outputs with data. The model accurately predicted PCB accumulation and negative effects on embryo survival, kit survival, growth and development. The DEBtox parameters according to Table 7 were estimated (Desforges et al. 2017).

First, DEBtox parameters are transformed into TDM parameters (for the same species). Assuming no recovery ($r = 0$), the damage rate according to TDM is:

$$\frac{\mathrm{d}d}{\mathrm{d}\tau} = \sigma c(\tau) \tag{35}$$

Assuming no initial damage and constant internal concentration $c$, the cumulated damage $d$ after exposure time $\Delta\tau$ is obtained as:

$$d = \sigma c \Delta\tau \tag{36}$$

Assuming no initial hazard, the cumulated hazard $h$ after exposure time $\Delta\tau$ is obtained from Eq. (120) as:

$$h = [d - d_\mathrm{T}]_+ = [\sigma c \Delta\tau - d_\mathrm{T}]_+ \quad , \quad [x]_+ = \max(0, x) \tag{37}$$

The hazard according to DEBtox theory is:

$$h = [c - \mathrm{NEC}]_+ / c_\mathrm{T} \tag{38}$$

Notice that hazard according to DEBtox theory only depends on the current toxicant concentration and is independent of exposure time. Comparing Eqs. (37) and (38) yields expressions for TDM parameters expressed in DEBtox parameters. The killing rate constant $\sigma$ is expressed in terms of the tolerance concentration $c_\mathrm{T}$ and the exposure time $\Delta\tau$ as:

$$\sigma \approx 1/(c_\mathrm{T}\Delta\tau) \tag{39}$$

The damage threshold level $d_\mathrm{T}$ is expressed in terms of the no effect concentration NEC and the tolerance concentration $c_\mathrm{T}$ as:

$$d_\mathrm{T} \approx \mathrm{NEC}/c_\mathrm{T} \tag{40}$$

An upper limit for the recovery rate constant $r$ may be estimated by assuming zero damage rate:

$$\frac{\mathrm{d}d}{\mathrm{d}\tau} = \sigma c(\tau) - r_{max} d(\tau) = 0 \tag{41}$$

The maximum recovery rate constant is thus obtained as:

$$r_{max} = \sigma c/d \approx \sigma c_T/d_\mathrm{T} = 1/(d_\mathrm{T}\Delta\tau) \tag{42}$$

Since TDM includes cumulative effects, not considered in a DEBtox model, translation of DEBtox parameters into TDM parameters, requires knowledge of exposure time $\Delta\tau$ used in the experiment. Mink DEBtox parameters were converted to TDM parameters by applying Eqs. (40) and (42) with exposure time $\Delta\tau_g^{mink} = 42$ days and are presented in Table 7. The TDM parameters for mink were then converted to TDM parameters for grey seal. According to Baas and Kooijman (2015) there is a strong correlation between specific metabolic rate (equated with specific somatic maintenance) and sensitivity to a toxicant (defined as having low NEC). This makes it possible to scale NEC between different animals. From analyses with different combinations of toxicants and animals, Baas & Kooijman (2015) found a strong negative correlation between specific somatic maintenance $p_m$ and no effect concentration NEC:

$$\log(p_m) = m - b \cdot \log(\mathrm{NEC}) \tag{43}$$

Here, $m$ and $b$ are toxicant-specific constants. If $p_m$ and NEC are known for two animals, exposed to a specified toxicant, $m$ and $b$ can be calculated. If $p_m$ is known for a third species, NEC can be estimated also for this one. The specific somatic maintenance for American mink (*Neovison vison*) and grey seal (*Halichoerus grypus*) is found in the Add-My-Pet database (Kooijman et al. 2019):

$$p_m^{mink} = 88.71\ \mathrm{J/(day \cdot cm^3)} \quad , \quad p_m^{seal} = 37.52\ \mathrm{J/(day \cdot cm^3)} \tag{44}$$

With known value on the PCB-specific parameter $b$, DEBtox data for mink can be converted to DEBtox data for grey seal:

$$\frac{\mathrm{NEC}^{seal}}{\mathrm{NEC}^{mink}} = \left[\frac{p_m^{mink}}{p_m^{seal}}\right]^{1/b} \tag{45}$$

It was here assumed that $b = 1$ and that damage threshold levels can be converted in the same way as no effect concentrations:

$$\frac{d_T^{seal}}{d_T^{mink}} = \left[\frac{p_m^{mink}}{p_m^{seal}}\right]^{1/b} \approx 2.36 \tag{46}$$

It follows that tolerance parameters are the same for different mammals exposed to a specified toxicant ($c_T^{mink} = c_T^{seal}$). Hence, also killing rate constants are the same, assuming equal exposure times.

$$\sigma^{seal} = \sigma^{mink} \tag{47}$$

By applying Eqs.(42), (46) and (47), TDM parameters for mink are converted to TDM parameters for grey seal.

### 3. Toxicokinetic Model

The toxicokinetic model describes PCB bioaccumulation in pups and females through dietary uptake and elimination as well as vertical transfer from mother to embryo through the placenta during gestation and from mother to pup through breast milk during lactation (Fig. 4).

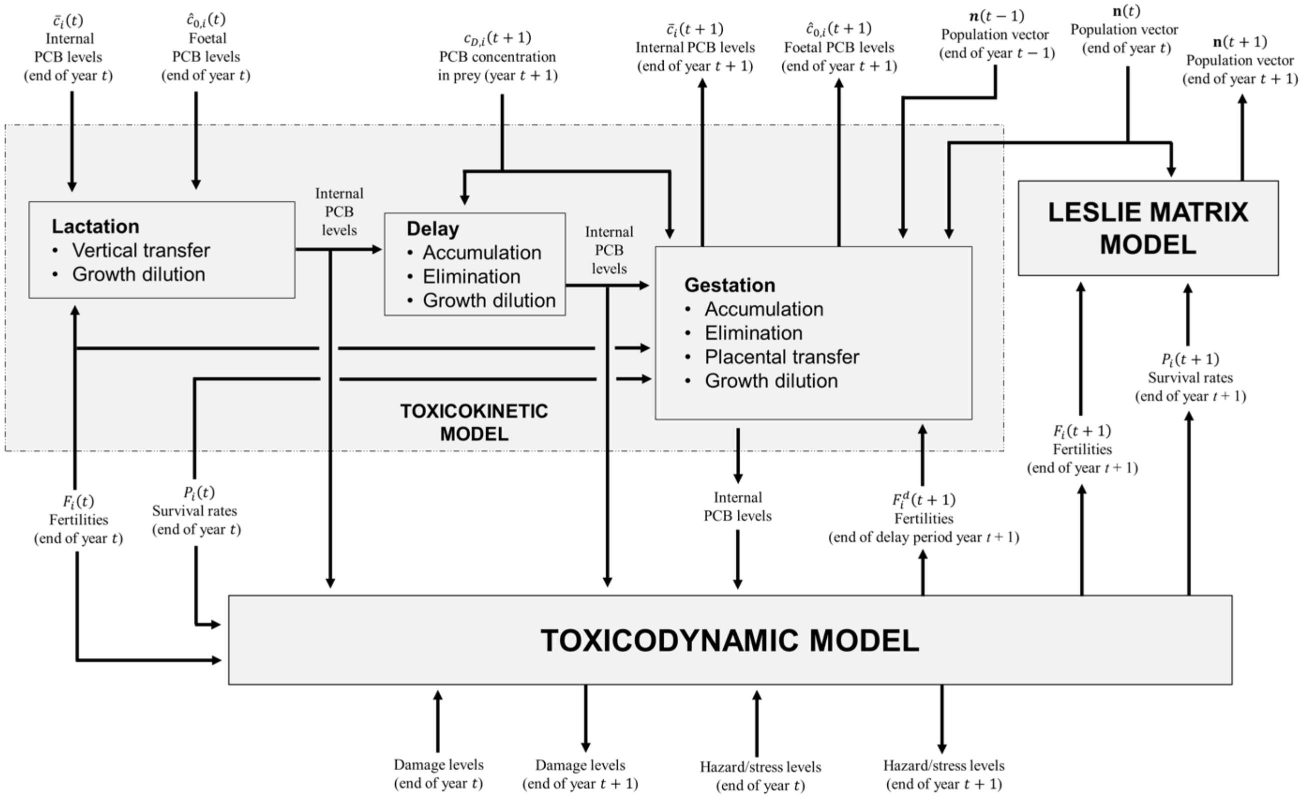

**Fig. 4** The toxicokinetic model and its connections to other sub models. Text boxes represent sub modules. Free texts are inputs and outputs, with arrows showing relations to sub modules. The flowchart illustrates computations performed for year $t + 1$, based on inputs from the previous year $t$.

A year is divided into three periods (*lactation*, *delay* and *gestation*) as illustrated in Fig. 5, representing different phases for a reproducing grey seal female. The year starts with the *lactation period* (lasting for 18 days), during which pups are nursed by their mothers. After weaning, females mate and the *delay period* starts (lasting for 100 days), after which the fertilized egg is implanted. During the *gestation period* (lasting for 247 days), the fetus develops in the uterus. The year ends with the birth of new pups. Since all deaths are assumed to occur just before the birth events, all pups that are born have mothers that nurse them.

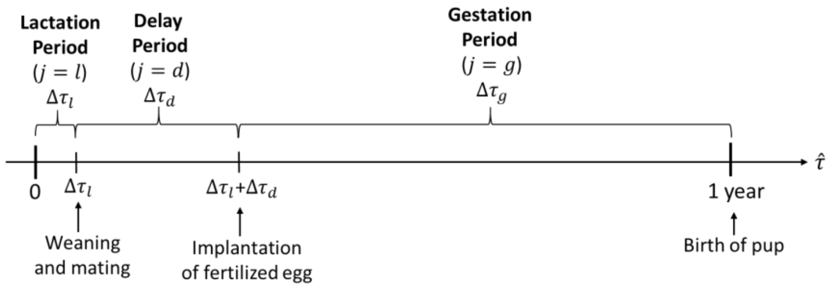

**Fig. 5** Division of a year into three periods, representing the reproduction cycle of grey seal females. $\Delta\tau_l = 18$ days, $\Delta\tau_d = 100$ days, $\Delta\tau_g = 247$ days.

During the *lactation period*, PCB is transferred from females to pups through the breast milk. Neither females nor pups consume prey during this period. Hence, no PCB is accumulated from the environment, but it is redistributed from females to pups through vertical transfer, including some losses to the environment. PCB concentrations also change due to growth dilution, especially in the fast growing pups. During the *delay period*, both females and pups feed on prey. Hence, PCB concentrations change due to dietary uptake, elimination (fecal egestion and metabolic transformation) and growth dilution. During the *gestation period*, females and pups continue to feed, but females also transfer PCB to fetuses through the placenta. Hence, concentration changes in pups are governed by dietary uptake, elimination, and growth dilution, whereas concentration changes in females are governed by dietary uptake, elimination, growth dilution and vertical transfer. Moreover, the PCB concentration increases in fetuses due to vertical transfer, which affect the PCB concentration in new-born pups at the start of the next year.

#### 3.1 Growth model

Fetuses are assumed to growth exponentially during the gestation period (Peters 1983). Hence, the body weight of a fetus at time $\hat{\tau}$ during a year is obtained as:

$$W_0(\hat{\tau}) = W_{00}e^{g_0(\hat{\tau}-(\Delta\tau_l+\Delta\tau_d))}, \qquad \Delta\tau_l + \Delta\tau_d \leq \hat{\tau} \leq 1 \tag{48}$$

Here, $W_{00}$ is the initial foetal weight (the weight of a fertilized egg) and $g_0$ is an exponential growth rate constant. At the end of the gestation period, the fetus has reached the birth weight; $W_0(1) = W_{birth}$. Hence, the initial fetal weight $W_{00}$ is calculated as:

$$W_{00} = W_{birth}e^{-g_0\Delta\tau_g} \tag{49}$$

For simplicity it is assumed that new-born pups continue to growth exponentially with the same growth rate as fetuses during the lactation period. Hence, the body weight of a pup at time $\hat{\tau}$ during the lactation period is obtained as:

$$W_1(\hat{\tau}) = W_{00}e^{g_0(\Delta\tau_g+\hat{\tau})}, \quad 0 \leq \hat{\tau} < \Delta\tau_l \tag{50}$$

At the end of the lactation period, the pup has reached the weaning weight; $W_1(\Delta\tau_l) = W_{wean}$. Hence, the exponential growth rate constant $g_0$ is calculated as:

$$g_0 = ln(W_{wean}/W_{birth})/\Delta\tau_l \tag{51}$$

After weaning, grey seal pups typically lose weight during some months due to reduced nutritional intake (Boyd 1984), but when they become more skilled hunters they start to regain weight. For simplicity, it is here assumed that pups keep the weaning weight until they reach sub adult age $\tau_{SA} = 1$ year.

Grey seal females are typically full-grown at an age of six years. When they nurse pups (during the lactation period) they lose a considerable amount of weight, but this is regained when they feed during the remaining part of the year. Under constant environmental conditions and dynamic energy budget theory (DEB) assumptions for feeding rate and somatic maintenance, DEB theory predicts body growth according to von Bertalanffy growth function (Kooijman 2000), in which the body length $L(\tau)$ at age $\tau$ is obtained as (von Bertalanffy 1938):

$$L(\tau) = L_\infty - (L_\infty - L_0)e^{-\gamma(\tau-\tau_0)} \tag{52}$$

Here, $L_\infty$ is the asymptotic adult body length, $L_0$ is the initial length (the length at $\tau = \tau_0$) and $\gamma$ is a growth rate constant. The von Bertalanffy growth function is here adopted to describe body growth of grey seal females from weaning ($\tau = \Delta\tau_l$) up to the age of sexual maturation ($\tau_{mat} = 5$ years). At that age, females reach the mature body length $L_{mat}$ and start to breed. The von Bertalanffy growth rate constant $\gamma$ is calculated as:

$$\gamma = \frac{ln[(L_\infty - L_{wean})/(L_\infty - L_{max})]}{\tau_{mat} - \tau_{SA} + \Delta\tau_g} \tag{53}$$

The body length $L(\tau)$ at age $\tau$ for non-mature females is given by:

$$L(\tau) = L_\infty - (L_\infty - L_{wean})e^{-\gamma(\tau-\tau_{SA})}, \qquad \tau_{SA} \leq \tau \leq \tau_{mat} \tag{54}$$

The asymptotic body length $L_\infty$ and the weaning body length $L_{wean}$ are obtained from the asymptotic body weight $W_\infty$ and the weaning body weight $W_{wean}$, assuming isometric growth and a body weight $W$ proportional to the structural body, in accordance with DEB theory (Kooijman 2000):

$$W(\tau) = \alpha L(\tau)^3 \tag{55}$$

Here, $\alpha$ is a body shape constant. With specified maximum body length and weight ($L_{max}$ and $W_{max}$), based on data for full-grown Baltic grey seal females, the body shape constant $\alpha$ is calculated as:

$$\alpha = W_{max}/L_{max}^3 \tag{56}$$

The asymptotic body length $L_\infty$ and the weaning body length $L_{wean}$ are obtained from specified values of the asymptotic body weight $W_\infty$ and the weaning body weight $W_{wean}$:

$$L_\infty = (W_\infty/\alpha)^{1/3}\,, \qquad L_{wean} = (W_{wean}/\alpha)^{1/3} \tag{57}$$

At sexual maturation, the female has reached its mature body weight $W_{mat}$. Since primiparous females (females which give birth for the first time) are typically smaller than multiparous females (Lang et al. 2011), the mature body weight $W_{mat}$ is assumed to be somewhat lower than the maximum body weight $W_{max}$, reached by multiparous females just before they start to lactate (when females are largest). Females lose significant weight during lactation and it is assumed that their weight is exponentially decreasing throughout the lactation period, from the initial body weight ($W_{mat}$ or $W_{max}$) to the post-lactation body weight; $W_{lac,6}$ or $W_{lac,7}$ (for primiparous and multiparous females, respectively):

$$\left.\begin{array}{l} W_6(\hat{\tau}) = W_{mat}e^{-g_L\hat{\tau}} \\ W_i(\hat{\tau}) = W_{max}e^{-g_L\hat{\tau}} \quad (i = 7, \dots, 46) \end{array}\right\} 0 \leq \hat{\tau} < \Delta\tau_l \tag{58}$$

Here, $g_L$ is the exponential body weight decline rate constant, which is calculated as:

$$g_L = ln\left(W_{max}/W_{lac,M}\right)/\Delta\tau_l \tag{59}$$

After lactation and during the delay period, mature females are assumed to linearly regain the weight they lost during lactation:

$$\left.\begin{array}{l} W_6(\hat{\tau}) = W_{lac,6} + k_6(\hat{\tau} - \Delta\tau_l) \\ W_i(\hat{\tau}) = W_{lac,7} + k_7(\hat{\tau} - \Delta\tau_l) \quad (i = 7, \dots, 46) \end{array}\right\} \Delta\tau_l \leq \hat{\tau} < \Delta\tau_l + \Delta\tau_d \tag{60}$$

Here, $k_6$ and $k_7$, are regain rate constants for primiparous and multiparous females, respectively, and are calculated as:

$$k_6 = \left(W_{mat}/W_{lac,6}\right)/\Delta\tau_d \quad , \quad k_7 = \left(W_{max}/W_{lac,7}\right)/\Delta\tau_d \tag{61}$$

After the delay period and during the gestation period, mature females continue to grow according to von Bertalanffy and reach the maximum body weight $W_{max}$ at the end of the year. In succeeding years, females are assumed to repeat the pattern of exponential decline (lactation period), linear regain (delay period) and von Bertalanffy growth (gestation period) (Fig. 6).

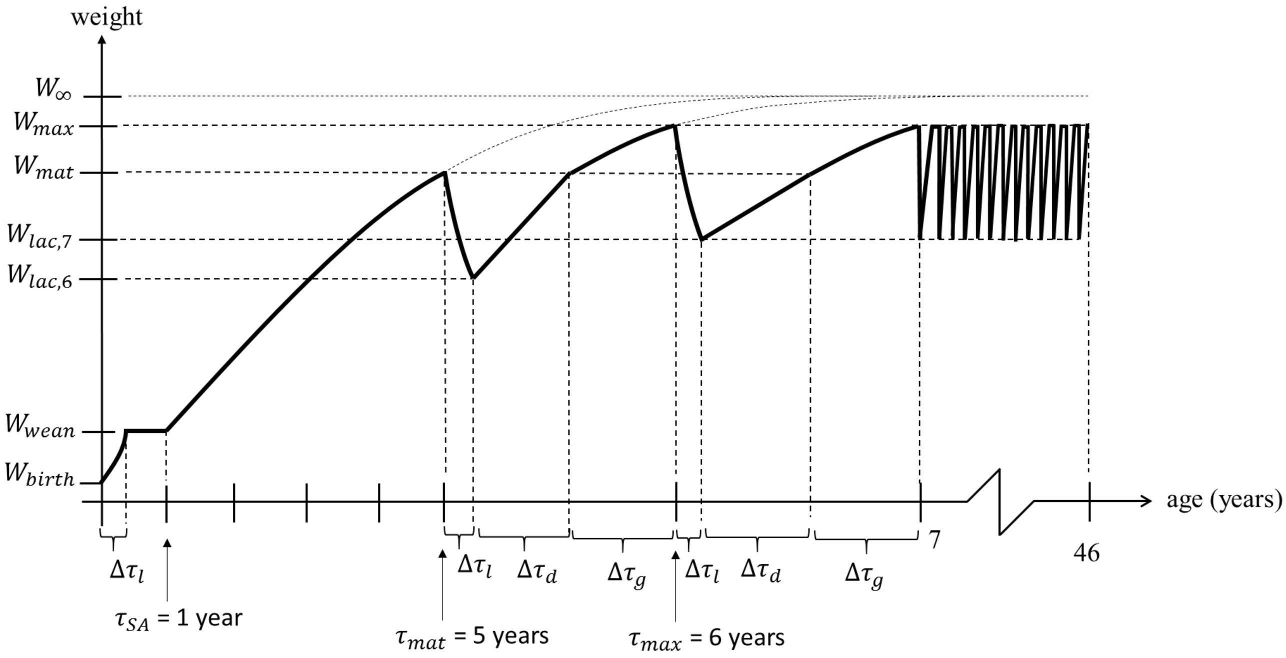

**Fig. 6** Body weight as a function of age according to model for body growth. Pups increase their weight exponentially during lactation and keep it constant during the remainder of the year. Sub-adults follow von Bertalanffy growth function until maturation. Mature females lose weight exponentially during lactation, regain it linearly during the delay period and grow according to von Bertalanffy during gestation. The same yearly pattern for mature females is then repeated.

The mean body weight $\bar{W}_i^j$ during time period $j$ for seals following von Bertalanffy growth is obtained from the mean body length $\bar{L}_i^j$, calculated as the integrated mean value over the period ($\hat{\tau}_0^j \le \hat{\tau} \le \hat{\tau}_0^j + \Delta\tau_j$):

$$\bar{W}_i^j = \alpha\left[\bar{L}_i^j\right]^3 \quad , \quad \text{where} \quad \bar{L}_i^j = \frac{1}{\Delta\tau_j}\int_{\hat{\tau}_0^j}^{\hat{\tau}_0^j+\Delta\tau_j} L_i(\hat{\tau})\mathrm{d}\hat{\tau} \qquad (62)$$

Remaining mean body weights during different time periods are calculated as integrated mean values:

$$\bar{W}_i^j = \frac{1}{\Delta\tau_j}\int_{\hat{\tau}_0^j}^{\hat{\tau}_0^j+\Delta\tau_j} W_i(\hat{\tau})\mathrm{d}\hat{\tau} \qquad (63)$$

#### 3.2 Bioaccumulation model

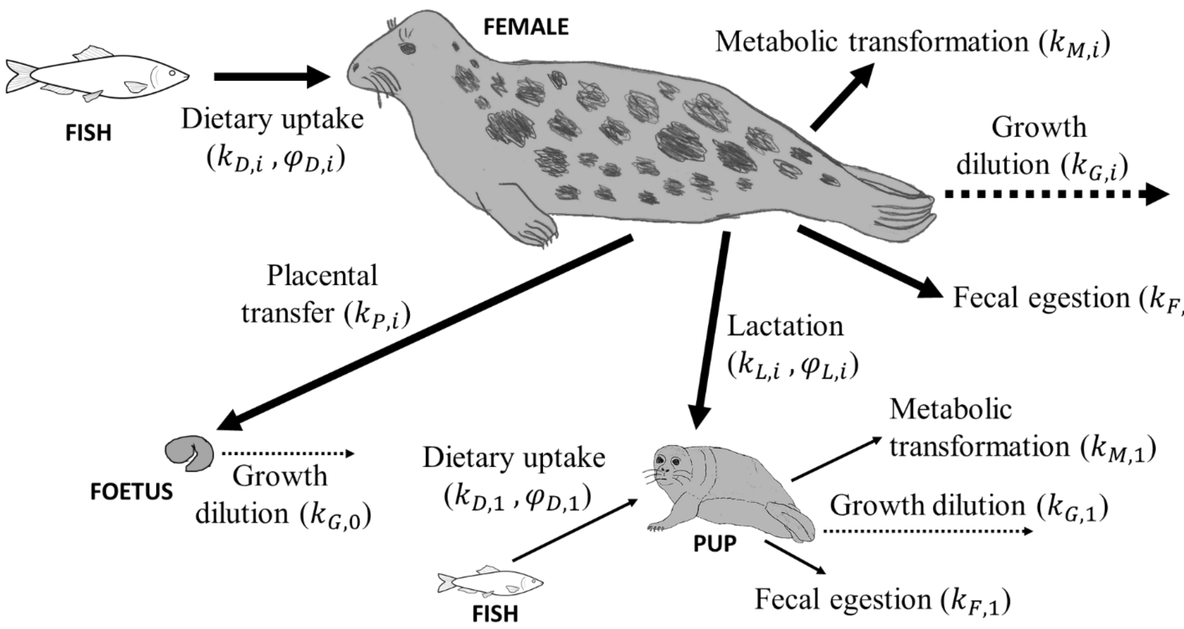

**Fig. 7** Major routes for uptake, transfer and elimination of PCB in grey seals, including associated rate constants ($k_{D,i}, k_{M,i}, k_{F,i}, k_{L,i}, k_{P,i}, k_{G,i}$) and assimilation efficiencies ($\varphi_{D,i}, \varphi_{L,i}$).

##### 3.2.1. Bioaccumulation in females

Bioaccumulation in females older than one year ($i = 2, \ldots, 46$) includes dietary uptake, elimination and vertical transfer to offspring (Fig. 7). The toxicokinetics is described by a physiologically based equation similar to the one-compartment model commonly used in DEBtox models, but extended to account also for growth dilution and vertical transfer. The rate of change of the total amount of PCB in a female is expressed as the rate by which PCB is assimilated (assumed to be proportional to the concentration in the prey) subtracted by the rate by which PCB is eliminated (assumed to be proportional to the concentration in the body):

$$\frac{\mathrm{d}}{\mathrm{d}\tau}\left(W_i c_{i,i}^j\right) = k_{D,i}^j W_i c_{D,i} - k_{E,i}^j W_i c_{i,i}^j \ , \qquad i = 2, \ldots, 46 \quad , \ j = l, d, g \ , \qquad 0 \le \tau \le \Delta\tau_j \qquad (64)$$

Here, $c_{i,i}^j(\tau)$ is the PCB concentration (mg/kg) at time $\tau$ (years) in a female of age class $i$ during time period $j$, whereas $c_{D,i}$ is the mean PCB concentration in the diet (mg/kg) and $W_i(\tau)$ is the female body weight (kg). Notice that $c_{i,i}^j$ is the total body concentration, which can be converted to blubber concentration (from an assumed relationship, see below). The dietary uptake rates (year$^{-1}$) for females (and pups) are defined as:

$$k_{D,i}^j = \begin{cases} 0 & , \quad j = l \\ \varphi_{D,i}\dot{W}_{D,i}/W_i & , \quad j = d, g \end{cases} \quad , \qquad i = 1, \ldots, 46 \qquad (65)$$

Here, $\dot{W}_{D,i}$ is the prey consumption rate (kg/year) and $\varphi_{D,i}$ is the diet assimilation efficiency; the fraction of consumed prey that is assimilated into energy, assumed to equal the fraction of consumed PCB that is kept by the seal. The assumption is motivated by studies indicating that the assimilation of hydrophobic substances is closely linked to digestion of associated food (Bodiguel et al. 2009). We assume an assimilation efficiency of 90% as described in Hickie et al. (2005) for their ringed seal bioaccumulation model. Notice that dietary uptake rates are zero when $j = l$ since neither females nor pups feed during lactation. As a simplification, it is assumed that also non-mature females ($i = 2, \dots, 5$) fast during the lactation period, though they have no pups to nurse. This simplification will only have minor effects since the lactation period is very short (18 days) compared to a whole year. According to DEB theory, the food uptake of an organism is proportional to the surface area of the structural body volume (Kooijman 2000). Assuming isometric growth and a weight proportional to the structural body volume, the prey consumption rate $\dot{W}_D$ for an individual of weight $W$ is then obtained as:

$$\dot{W}_D = \alpha_D \kappa W^{2/3} \tag{66}$$

Mature females lactate and hence they lose and regain weight during a year. Due to their rapid weight regain after lactation, they can be expected to peek their daily prey consumption during the delay period, though they have higher weight during other periods. The weight $W$ in Eq. (66) is best considered as a mean body weight for females of a certain body length (which do not reduce during fasting). As a simplification, it is here assumed that the prey consumption rate for individuals of age class $i$ can be described by Eq. (66) with $W$ as the maximum body weight of a year, i.e. the body weight at the end of the year ($\hat{\tau} = 1$). The age-specific dietary uptake rate constants for females (and pups) are approximated as:

$$k_{D,i}^{j} \approx \bar{k}_{D,i}^{j} = \begin{cases} 0 & , \quad j = l \\ \alpha_D \varphi_{D,i} [W_i(1)]^{\,2/3} / \bar{W}_i^{\,j} & , \quad j = d, g \end{cases} \quad , \quad i = 1, \dots, 46 \tag{67}$$

Here, $\bar{W}_i^{\,j}$ is the mean body weight of individuals in age class $i$ during time period $j$.

The prey consumption factor $\alpha_D$ is a constant, estimated from empirical data:

$$\alpha_D = \frac{\dot{W}_{D,max}}{W_{max}^{2/3}} \approx \frac{\bar{W}_{\max}^{\text{fish}} / (\Delta\tau_d + \Delta\tau_g)}{W_{max}^{2/3}} \tag{68}$$

The prey consumption rate $\dot{W}_{D,max}$ for the largest females (with body weight $W_{max}$) is estimated as the empirically obtained maximum yearly fish consumption of large grey seal females ($\bar{W}_{\max}^{\text{fish}}$) divided by the length of the total feeding period during a year ($\Delta\tau_d + \Delta\tau_g$). Hansson, et al. (2017), estimated the daily mean prey consumption of Baltic grey seals to 4.5-5.0 kg, corresponding to an annual per capita consumption of 1750 kg. It was here assumed that the largest females (with a weight of $W_{\max} = 160$ kg) has a yearly fish consumption of $\bar{W}_{\max}^{\text{fish}} = 1800$ kg. Notice the lactation period is excluded, whereas seals are assumed to feed throughout the whole delay period, neglecting the two weeks of fasting during molting. The female PCB elimination rates (year$^{-1}$), accounting for all elimination routes, are defined as:

$$k_{E,i}^{j} = \begin{cases} k_{L,i} & , \quad j = l \\ k_{R,i} & , \quad j = d \\ k_{R,i} + k_{P,i} & , \quad j = g \end{cases} \quad , \quad i = 2, \dots, 46 \tag{69}$$

Elimination through vertical transfer is accounted for via the lactation rate constant $k_{L,i}$ and the placental transfer rate constant $k_{P,i}$, respectively. The removal rate constant, accounting for metabolic transformation and fecal egestion, is defined as:

$$k_{R,i} = k_{M,i} + k_{F,i} \quad , \quad i = 1, \dots, 46 \tag{70}$$

Here, $k_{M,i}$ is the metabolic transformation rate constant and $k_{F,i}$ is the fecal egestion rate constant. The vertical transfer rate constants ($k_{L,i}$ and $k_{P,i}$) are considered as mean values for each period. This simplification neglects that vertical transfer rates increase with the weight of the offspring (fetus or pup), which grows and assimilates increasing amounts of nutrients, resulting in vertical transfer rates that increase with time. Fecal elimination occurs via two different routes. The first route is through the energy reserves and the corresponding elimination rate is a fraction of the uptake rate, included via the diet assimilation efficiency $\varphi_{D,i}$. The second route is elimination from the structural body volume and corresponding elimination rate is proportional to the concentration in the body, included via the fecal egestion rate constant $k_{F,i}$. The former is considered important for aquatic organisms with a major uptake of toxicants from the surrounding water (Kooijman 2000). The fecal egestion rate constant may be neglected for mammals, which do not assimilate PCB from surrounding water.

Notice that all rate constants are assumed to be age-specific, though some of them have similar or same values for all age classes. For the sake of generality, Eq. (64) accounts for all assimilation and elimination routes, including vertical transfer from female to offspring, though all these will never occur at the same time. When applying Eq. (64) to different time periods of a year, different rate constants appear according to Eqs. (67) and (69).

The toxicokinetic equation (Eq. (64)) can be rewritten as the following differential equation for the PCB concentration $c_{i,i}^{j}(\tau)$ at time $\tau$ in a female seal of age class $i$ during time period $j$:

$$\frac{\mathrm{d}c_{i,i}^{j}}{\mathrm{d}\tau} + \left(k_{E,i}^{j} + k_{G,i}(\tau)\right)c_{i,i}^{j} = \bar{k}_{D,i}^{j} c_{D,i}(\tau) \tag{71}$$

The *growth dilution rate* ($k_{G,i}$), the relative increase in body weight per unit time, was here introduced:

$$k_{G,i}(\tau) = \frac{1}{W_i}\frac{\mathrm{d}W_i}{\mathrm{d}\tau} \quad , \quad i = 1, \ldots, 46 \tag{72}$$

It is assumed that the mean PCB concentration in the prey is constant throughout each year; $c_{D,i}(\tau) = \bar{c}_{D,i}(t)$. Here, $\bar{c}_{D,i}(t)$ is the mean PCB concentration in the diet of seals in age class $i$ during year $t$. Moreover, it is assumed that the growth dilution rate is approximately constant within each period of a year. The growth dilution rate ($k_{G,i}$) for seals of age class $i$ during a time period $j$ ($\hat{\tau}_0^j \leq \hat{\tau} \leq \hat{\tau}_0^j + \Delta\tau_j$) is approximated as the mean growth dilution rate ($\bar{k}_{G,i}^{j}$) during that period:

$$k_{G,i} \approx \bar{k}_{G,i}^{j} = \frac{1}{\Delta\tau_j}\int_{\hat{\tau}_0^j}^{\hat{\tau}_0^j+\Delta\tau_j} k_{G,i}(\hat{\tau})\mathrm{d}\hat{\tau} = \frac{1}{\Delta\tau_j}\left(\ln\left[W_i\left(\hat{\tau}_0^j + \Delta\tau_j\right)\right] - \ln\left[W_i\left(\hat{\tau}_0^j\right)\right]\right) \quad , \quad \begin{cases} i = 1, \ldots, 4 \\ j = l, d, g \end{cases} \tag{73}$$

During the lactation period, body weights of pups and females are described by exponential functions (Eqs. (50) and (58)). The growth dilutions rates then simplify into the exponential growth rate constants:

$$\bar{k}_{G,1}^{l} = k_{G,1}^{l} = g_0 \quad , \quad \bar{k}_{G,\mathrm{i}}^{l} = k_{G,i}^{l} = -g_{\mathrm{L}} \qquad i = 6, \ldots, 46 \tag{74}$$

During the rest of the year, pups have constant body weight and hence no growth dilution:

$$\bar{k}_{G,1}^{j} = k_{G,1}^{j} = 0 \quad , \quad j = \mathrm{d}, \mathrm{g} \tag{75}$$

The total elimination rate for females (and pups) during a time period $j$ within a year is defined as:

$$\hat{k}_{E,i}^{j} = k_{E,i}^{j} + \bar{k}_{G,i}^{j} \quad , \quad i = 1, \ldots, 46 \quad , \quad j = l, d, g \tag{76}$$

Now, the toxicokinetic equation (Eq. (71)) can be approximated as the following first-order ordinary differential equation (ODE):

$$\frac{\mathrm{d}c_{i,i}^{j}}{\mathrm{d}\tau} + \hat{k}_{E,i}^{j} c_{i,i}^{j} = \bar{k}_{D,i}^{j}\bar{c}_{D,i} \quad , \quad i = 2, \ldots, 46 \quad , \quad j = l, d, g \tag{77}$$

The solution to Eq. (77) is:

$$c_{i,i}^{j}(\tau) = \kappa_{D,i}^{j}\bar{c}_{D,i} + C_i^j e^{-\hat{k}_{E,i}^{j}\tau} \quad , \quad \begin{cases} \kappa_{D,i}^{j} = \bar{k}_{D,i}^{j}/\hat{k}_{E,i}^{j} \\ C_i^j = c_{i,i}^{j}(0) - \kappa_{D,i}^{j}\bar{c}_{D,i} \end{cases} \tag{78}$$

Equation (78) expresses the PCB concentration in a female of age class $i$ at time $\tau$ during period $j$ within a year. The first term accounts for dietary uptake (characterized by constant $\kappa_{D,i}^{j}$) and the second term accounts for elimination (characterized by elimination rate $\hat{k}_{E,i}^{j}$). Equation (78) is used to obtain the PCB concentration in females as a result of bioaccumulation during different time periods $j$, based on a known initial PCB concentration $c_{i,i}^{j}(0)$ at the start of the period ($\tau = 0$).

The mean PCB concentration in females of age class $i+1$ at the end of the **lactation period** ($j = l,\ \tau = \Delta\tau_l$) during year $t+1$ is obtained from Eq.(78) as:

$$\bar{c}_{i+1}^{l}(t+1) = b_{i+1,i}^{l}(t+1)\bar{c}_i(t)\,, \qquad b_{i+1,i}^{l}(t) = e^{-\bar{k}_{E,i+1}^{l}(t)\Delta\tau_l} \quad , \quad i = 1,\dots,45 \tag{79}$$

Here, $\bar{c}_i(t)$ is the mean PCB concentration in females of age class $i$ at the end of year $t$. The mean elimination rate (accounting for lactational transfer and growth dilution) during the lactation period year $t$ is defined as:

$$\bar{k}_{E,i}^{l}(t) = \bar{k}_{L,i}(t) + \bar{k}_{G,i}^{l} \quad , \qquad i = 2,\dots,46 \tag{80}$$

The mean lactation rate during the lactation period year $t+1$ is calculated as:

$$\bar{k}_{L,i+1}(t+1) = 2F_i(t)k_{L,i+1} \quad , \qquad i = 1,\dots,45 \tag{81}$$

Here, $F_i(t)$ is the fertility of females in age class $i$ at the end of year $t$. Notice that the factor $2F_i(t)$ is the mean number of pups that each female (of age class $i+1$) gave birth to at the end of previous year ($t$). The factor is included to account for absence of lactation in females without pups.

The mean PCB concentration in females of age class $i+1$ at the end of the **delay period** ($j = d,\ \tau = \Delta\tau_d$) during year $t+1$ is obtained from Eq. (78) as:

$$\bar{c}_i^{d}(t+1) = b_{i,i}^{d}\bar{c}_i^{l}(t+1) + \phi_i^{d}\bar{c}_{D,i}(t+1) \quad i = 2,\dots,46 \quad , \quad \begin{cases} b_{i,i}^{d} = e^{-\hat{k}_{E,i}^{d}\Delta\tau_d} \\ \phi_i^{d} = \kappa_{D,i}^{d}\left(1 - b_{i,i}^{d}\right) \end{cases} \tag{82}$$

The mean PCB concentration in females of age class $i+1$ at the end of year $t+1$, i.e. at the end of the **gestation period** ($j = g,\ \tau = \Delta\tau_g$) year $t+1$, is obtained from Eq. (78) as:

$$\bar{c}_i(t+1) = b_{i,i}^{g}(t+1)\bar{c}_i^{d}(t+1) + \phi_i^{g}(t+1)\bar{c}_{D,i}(t+1) \quad i = 2,\dots,46 \quad , \quad \begin{cases} b_{i,i}^{g}(t) = e^{-\bar{k}_{E,i}^{g}(t)\Delta\tau_g} \\ \phi_i^{g}(t) = \bar{\kappa}_{D,i}^{g}[1 - b_{i,i}^{g}(t)] \\ \bar{\kappa}_{D,i}^{g} = \bar{k}_{D,i}^{j}/\bar{k}_{E,i}^{j} \end{cases} \tag{83}$$

The mean elimination rate (accounting for removal, placental transfer and growth dilution) during the gestation period is:

$$\bar{k}_{E,i}^{g}(t) = k_{R,i} + \bar{k}_{P,i}(t) + \bar{k}_{G,i}^{g} \quad , \qquad i = 2,\dots,46 \tag{84}$$

The mean placental transfer rate (for females of age class $i+1$) during the gestation period (year $t+1$) is:

$$\bar{k}_{P,i}(t) = 2F_i^{d}(t)k_{P,i} \quad , \qquad i = 2,\dots,46 \tag{85}$$

Here, $F_i^{d}(t)$ is the fertility for females of age class $i$ at the end of the delay period year $t$, a fertility reduced with respect to reproductive stress (obtained from the toxicodynamic model). The factor $2F_i^{d}(t)$ is an estimation of the mean number of fetuses that each female (of age class $i$) produces during year $t$. The factor is included to account for absence of placental transfer in non-pregnant females. Equations (79), (82) and (83) are used to successively calculate PCB concentrations in females at the end of the lactation period, the delay period and the gestation period for one year at a time.

##### 3.2.2. Bioaccumulation in Fetuses

Fetuses assimilate PCB only through vertical transfer via the placenta. It is assumed that all PCB that is eliminated by a mother through placental transfer is assimilated by her fetus, which has no capacity to eliminate it. The toxicokinetic equation is a mass balance where the rate of change of PCB in the fetus equals the rate at which PCB is eliminated from the mother through placental transfer. The rate of change of the total amount of PCB in a fetus with a mother of age class $i$ during the gestation period ($j = g$) is:

$$\frac{\mathrm{d}}{\mathrm{d}\tau}\left(W_0 c_{0,i}^{g}\right) = k_P W_i c_{i,i}^{g} \quad , \qquad i = 2,\dots,46\,, \qquad 0 \le \tau \le \Delta\tau_g \tag{86}$$

Here, $c_{0,i}^g$ and $c_{i,i}^g$ are PCB concentrations in fetus and mother, $W_0$ and $W_i$ are body weights of fetus and mother, whereas $k_P$ is the placental transfer rate constant. The toxicokinetic equation (Eq. (86)) can be rewritten as:

$$\frac{\mathrm{d}c_{0,i}^g}{\mathrm{d}\tau} + k_{G,0}c_{0,i}^g = k_P\omega_{0,i}c_{i,i}^g \quad , \quad i = 2, \dots, 46 \tag{87}$$

The fetus grows exponentially according to Eq. (48). Hence, the growth dilution rate $k_{G,0}$ equals the exponential growth rate:

$$k_{G,0} = \frac{1}{W_0}\frac{\mathrm{d}W_0}{\mathrm{d}\tau} = g_0 \tag{88}$$

The body weight ratio between female and fetus ($\omega_{0,i}$) is approximated as a mean value ($\bar{\omega}_{0,i}$) over the gestation period:

$$\omega_{0,i} = W_i/W_0 \approx \bar{\omega}_{0,i} = \bar{W}_i^g/\bar{W}_0 \quad , \quad i = 2, \dots, 46 \tag{89}$$

The PCB concentration in the mother ($c_{i,i}^g$) during the gestation period ($j = g$) is obtained from Eq. (78) and inserted into the toxicokinetic equation (Eq. (87)), which yields the following first order ODE:

$$\frac{\mathrm{d}c_{0,i}^g}{\mathrm{d}\tau} + k_{G,0}c_{0,i}^g = \bar{k}_{P0,i}\left(C_i^g e^{-\hat{k}_{E,i}^g\tau} + \kappa_{D,i}^g\bar{c}_{D,i}\right) \quad , \quad \bar{k}_{P0,i} = k_P\bar{\omega}_{0,i} \; , \quad i = 2, \dots, 46 \tag{90}$$

The solution to Eq. (90) is:

$$c_{0,i}^g(\tau) = \kappa_{P0,i}\bar{c}_{D,i} + \hat{\kappa}_{P0,i}C_i^g e^{-\hat{k}_{E,i}^g\tau} + C_{0,i}^g e^{-k_{G,0}\tau} \quad , \quad \begin{cases} \hat{\kappa}_{P0,i} = \bar{k}_{P0,i}/(k_{G,0} - \hat{k}_{E,i}^g) \\ \kappa_{P0,i} = \kappa_{D,i}^g\bar{k}_{P0,i}/k_{G,0} \\ C_{0,i}^g = (\kappa_{D,i}^g\,\hat{\kappa}_{P0,i} - \kappa_{P0,i})\bar{c}_D - \hat{\kappa}_{P0,i}c_{i,i}^g(0) \end{cases} \tag{91}$$

Equation (91) expresses the PCB concentration in a fetus with a mother of age class $i$ at time $\tau$ during the gestation period ($j = g$). The first term accounts for dietary uptake in the mother, transferred to the fetus. The second term accounts for reduction of transferred PCB due to elimination in the mother (characterized by elimination rate $\hat{k}_{E,i}^g$). The last term accounts for growth dilution in the fetus (characterized by growth dilution rate $k_{G,0}$). From Eq. (91), with $\tau = \Delta\tau_g$, the PCB concentration in fetuses with mothers of age class I at the end of year t+1 is obtained as:

$$\begin{gathered} \hat{c}_{0,i}(t+1) = \hat{b}_{i,i}^0\bar{c}_i^d(t+1) + \hat{\phi}_i^0\bar{c}_{D,i}(t+1) \quad , \quad i = 2, \dots, 46 \\ \hat{b}_{i,i}^0 = \hat{\kappa}_{P0,i}\left(e^{-\hat{k}_{E,i}^g\Delta\tau_g} - e^{-k_{G,0}\Delta\tau_g}\right) \quad , \quad \hat{\phi}_i^0 = \kappa_{P0,i}(1 - e^{-k_{G,0}\Delta\tau_g}) - \kappa_{D,i}^g\hat{b}_{i,i}^0 \end{gathered} \tag{92}$$

Equation (92) is used to calculate PCB concentrations in fetuses at year end, based on known PCB levels in their mothers at the end of the delay period.

##### 3.2.3. Bioaccumulation in Pups

Bioaccumulation of PCB in pups (age class $i = 1$) is described by a toxicokinetic equation, similar to Eq. (64) for females, but with no placental transfer and a positive contribution from vertical transfer through lactation. The rate of change of the total amount of PCB in a pup with a mother of age class $i$ is:

$$\begin{gathered} \frac{\mathrm{d}}{\mathrm{d}\tau}\left(W_1c_{1,i}^j\right) = k_{L1,i}^jW_ic_{i,i}^j + k_{D,1}^jW_1c_{D,1} - k_{E,1}^jW_1c_{1,i}^j \quad , \quad i = 2, \dots, 46 \, , \quad j = l, d, g \, , \\ 0 \le \tau \le \Delta\tau_j \end{gathered} \tag{93}$$

Here, $c_{1,i}^j$ and $c_{i,i}^j$ are PCB concentrations in pups and mothers, $c_{D,1}$ is the PCB concentration in the diet, whereas $W_1$ and $W_i$ are body weights of pups and mothers. The lactation rate $k_{L1,i}^j$ for pups with mothers of age class $i$, accounting for PCB uptake through breast milk during lactation, is defined as:

$$k_{L1,i}^{j} = \begin{cases} \varphi_L k_{L,i} & , \quad j = l \\ 0 & , \quad j = d, g \end{cases} \tag{94}$$

Here, $k_{L,i}$ is the lactation rate constant for females of age class $i$. The lactation assimilation efficiency $\varphi_L$ is the fraction of consumed milk that is assimilated into energy by a pup during lactation, assumed to equal the fraction of transferred PCB that is assimilated by the pup. We assumed the same assimilation efficiency as in diet (90%; Hickie et al, 2005). The dietary uptake rate for pups $k_{D,1}^{j}$ is calculated analogously to females (Eq. (65)), with prey consumption rate $\dot{W}_{D,1}$ and diet assimilation efficiency $\varphi_{D,1}$ specific to pups. The elimination rate (for pups), accounting for all elimination routes, is defined as:

$$k_{E,1}^{j} = \begin{cases} 0 & , \quad j = l \\ k_{R,1} & , \quad j = d, g \end{cases} \tag{95}$$

The removal rate constant for pups ($k_{R,1}$) is defined analogously to females (Eq. (70)), with metabolic transformation rate constant $k_{M,1}$ and fecal egestion rate constant $k_{F,1}$ specific to pups. The toxicokinetic equation (Eq. (93)) can be rewritten as the following differential equation for the PCB concentration $c_{1,i}(\tau)$ at time $\tau$ in a pup with a mother of age class $i$ during time period $j$:

$$\frac{\mathrm{d}c_{1,i}^{j}}{\mathrm{d}\tau} + \left(k_{E,1}^{j} + k_{G,1}(\tau)\right)c_{1,i}^{j} = k_{L1,i}^{j}\omega_{1,i}c_{i,i}^{j} + \bar{k}_{D,1}^{j}c_{D,1}(\tau) \quad , \quad i = 2, \dots, 46 \tag{96}$$

The body weight ratio between mother and pup is approximated as a mean value over the lactation period:

$$\omega_{1,i} = W_i/W_1 \approx \bar{\omega}_{1,i} = \bar{W}_i^{l}/\bar{W}_1^{l} \quad , \quad i = 2, \dots, 46 \tag{97}$$

Here, $\bar{W}_i^{l}$ $(i = 1, \dots, 46)$ are the mean body weights during the lactation period. The growth dilution rate $k_{G,1}$ is defined analogously to females (Eq. (72)) and is approximated as the mean growth dilution rate $\bar{k}_{G,1}^{j}$ during a time period $j$ ($\hat{\tau}_0^{j} \leq \tau \leq \hat{\tau}_0^{j} + \Delta\tau_j$) according to Eq. (73). The total elimination rate $\hat{k}_{E,1}^{j}$ for pups during time period $j$ is defined analogously to females (Eq. (76)). The mean lactation rate for a pup with a mother of age class $i$ is defined as:

$$\bar{k}_{L1,i}^{j} = k_{L1,i}^{j}\bar{\omega}_{1,i} = \begin{cases} \bar{k}_{L1,i} = \varphi_L k_{L,i}\bar{\omega}_{1,i} & , \quad j = l \\ 0 & , \quad j = d, g \end{cases} \tag{98}$$

With constant mean PCB concentration in the diet throughout each year ($c_{D,1} = \bar{c}_{D,1}$), the toxicokinetic equation (Eq. (96)) can be approximated as the following differential equation:

$$\frac{\mathrm{d}c_{1,i}^{j}}{\mathrm{d}\tau} + \hat{k}_{E,1}^{j}c_{1,i}^{j} = \bar{k}_{L1,i}^{j}c_{i,i}^{j} + \bar{k}_{D,1}^{j}\bar{c}_{D,1} \quad , \quad i = 2, \dots, 46 \quad , \quad j = l, d, g \tag{99}$$

The PCB concentration in the mother ($c_{i,i}^{j}$) during the lactation period ($j = l$) is obtained from Eq. (78) and inserted into Eq. (99), which yields the following first order ODE:

$$\frac{\mathrm{d}c_{1,i}^{j}}{\mathrm{d}\tau} + \hat{k}_{E,1}^{j}c_{1,i}^{j} = \bar{k}_{L1,i}^{j}c_{i,i}^{j}(0)e^{-\hat{k}_{E,i}^{j}\tau} + \bar{k}_{D,1}^{j}\bar{c}_{D,1} \quad , \quad i = 2, \dots, 46 \tag{100}$$

The solution to Eq. (100) for a time period $0 \leq \tau \leq \Delta\tau_j$ within a year is:

$$c_{1,i}^{j}(\tau) = \kappa_{D,1}^{j}\bar{c}_{D,1} + \hat{\kappa}_{L1,i}^{j}c_{i,i}^{j}(0)e^{-\hat{k}_{E,i}^{j}\tau} + C_{1,i}^{\mathrm{j}}e^{-\hat{k}_{E,1}^{j}\tau} \quad , \quad \begin{cases} \hat{\kappa}_{L1,i}^{j} = \bar{k}_{L1,i}^{j}/(\hat{k}_{E,1}^{j} - \hat{k}_{E,i}^{j}) \\ \kappa_{D,1}^{j} = \bar{k}_{D,1}^{j}/\hat{k}_{E,1}^{j} \\ C_{1,i}^{\mathrm{j}} = c_{1,i}(0) - \hat{\kappa}_{L1,i}^{j}c_i(0) - \kappa_{D,1}^{j}\bar{c}_{D,1} \end{cases} \tag{101}$$

Equation (101) expresses the PCB concentration in a pup with a mother of age class $i$ at time $\tau$ during time period $j$. The first term accounts for dietary uptake in the pup. The second term accounts for lactational transfer from mother to pup, reduced by elimination in the mother (characterized by elimination rate $\hat{k}_{E,i}^{j}$). The last term accounts for elimination in the pup (characterized by elimination rate $\hat{k}_{E,1}^{j}$). The initial PCB concentration in pups with mothers of age class $i+1$ (at the start of the lactation period year $t+1$) is assumed to equal the PCB concentration $\hat{c}_{0,i}(t)$ in fetuses with mothers of age class $i$ at the end of previous year $(t)$. The PCB concentration in pups with mothers of age class $i+1$ at the end of the **lactation period** $(j=l, \tau=\Delta\tau_l)$ year $t+1$ is obtained from Eq. (101) as:

$$\hat{c}_{1,\mathrm{i}+1}^{l}(t+1) = p\hat{c}_{0,i}(t) + \hat{b}_{i+1,\mathrm{i}}^{l}\bar{c}_i(t)$$
$$p = e^{-\bar{k}_{G,1}^{l}\Delta\tau_l}\ ,\ \ \hat{b}_{i+1,\mathrm{i}}^{l} = \hat{\kappa}_{L1,i+1}^{l}\left(e^{-\hat{k}_{E,i+1}^{l}\Delta\tau_l} - e^{-\bar{k}_{G,1}^{l}\Delta\tau_l}\right), \quad i = 1,\dots,45 \tag{102}$$

The PCB concentration in pups with mothers of age class $i$ at the end of the **delay period** $(j=d, \tau=\Delta\tau_d)$ year $t+1$ is obtained from Eq. (101) as:

$$\hat{c}_{1,i}^{d}(t+1) = \hat{b}^{d}\hat{c}_{1,\mathrm{i}}^{l}(t+1) + \hat{\phi}^{d}\bar{c}_{D,1}(t+1)\ ,$$
$$\hat{b}^{d} = e^{-\hat{k}_{E,1}^{d}\Delta\tau_d}\ , \qquad \hat{\phi}^{d} = \kappa_{D,1}^{d}(1-\hat{b}^{d}), \qquad i = 2,\dots,46 \tag{103}$$

The PCB concentration in pups at the end of the **gestation period** $(j=g, \tau=\Delta\tau_g)$ year $t+1$ is obtained from Eq. (101) as:

$$\hat{c}_{1,i}^{g}(t+1) = \hat{b}^{g}\hat{c}_{1,i}^{d}(t+1) + \hat{\phi}^{g}\bar{c}_{D,1}(t+1)\ ,$$
$$\hat{b}^{g} = e^{-\hat{k}_{E,1}^{g}\Delta\tau_g}\ , \qquad \hat{\phi}^{g} = \kappa_{D,1}^{g}(1-\hat{b}^{g})\ , \qquad i = 2,\dots,46 \tag{104}$$

Equations (102), (103) and (104) are used to successively calculate PCB concentrations in pups at the end of the lactation period, the delay period and the gestation period for one year at a time. Equation (104) gives the PCB concentration in pups with mothers of age class $i$ at the end of the gestation period year $t+1$.

The mean PCB concentration for all pups at the end of a year (just before deaths and births occur) is used as initial PCB level in age class 2 at the start of next year. At the end of year $t+1$, the total body weight of all female pups is:

$$W_{1,\mathrm{tot}}(t+1) = N_1(t)W_1(1) \tag{105}$$

Here, $N_1(t)$ is the number of pups born at the end of year $t$ and $W_1(1)$ is the body weight of pups at the end of a year. The total amount of PCB in all female pups (with mothers of all age classes) at the end of year $t+1$ is obtained as:

$$W_{1,\mathrm{tot}}^{\mathrm{PCB}}(t+1) = W_1(1)\sum_{i=1}^{45} P_i(t)F_i(t)N_i(t-1)\hat{c}_{1,i+1}^{g}(t+1) \tag{106}$$

Notice that $P_i(t)F_i(t)N_i(t-1)$ is the number of pups born by mothers of age class $i$ at the end of year $t$, i.e. the number of pups with mothers of age class $i+1$ during year $t+1$. Here, $P_i(t)$ and $F_i(t)$ is the survival rate and fertility of females in age class $i$ at the end of year $t$, whereas $N_i(t-1)$ is the number of females in age class $i$ at the end of year $t-1$ (after the occurrence of deaths). The mean PCB concentration for all pups is obtained as the total amount of PCB in all pups (Eq. (106)), divided by the total body weight of all pups (Eq. (105)):

$$\bar{c}_1(t+1) = \sum_{i=1}^{45} \eta_{1,i+1}(t+1)\hat{c}_{1,i+1}^{g}(t+1)\ , \quad \eta_{1,i+1}(t+1) = \begin{cases} P_i(t)F_i(t)N_i(t-1)/N_1(t) & , \quad N_1(t) \neq 0 \\ 0 & , \quad N_1(t) = 0 \end{cases} \tag{107}$$

Females that enter age class 2 are assumed to start the year with an initial PCB concentration that equals the mean PCB concentration in one year old pups at the end of previous year. Hence, Eq. (107) is used to obtain the initial PCB concentration in females of age class 2 at the start of a year.

##### 3.2.4. Prey Composition

The grey seal diet includes fish of different species. The mean PCB concentration in the diet of a seal in age class $i$ is calculated as:

$$\bar{c}_{D,i}(t) = \sum_j \phi_{ij} \bar{c}_{P,j}(t) \quad (108)$$

Here, $\bar{c}_{P,j}(t)$ is the mean PCB concentration in prey $j$ during year $t$ and the prey preference index $0 \leq \phi_{ij} \leq 1$ describes the preference for prey $j$ among seals of age class $i$ (the mean weight fraction of prey $j$ in the diet of a seal in age class $i$).

#### 3.3 Conversion of PCB concentrations

Empirical data of PCB concentrations in fish and seals and are usually presented on a lipid weight basis, whereas the TKTD population model expresses PCBs in total body concentrations. To make comparisons, it is necessary to convert total body concentrations to lipid concentrations. The body fat index for seals of age class $i$ during time period $j$ is expressed as:

$$\bar{\rho}_i^j = \bar{W}_i^{\text{lip},j} / \bar{W}_i^j \quad (109)$$

Here, $\bar{W}_i^j$ is the mean body weight (kg) of age class $i$ during time period $j$ and $\bar{W}_i^{\text{lip},j}$ is the corresponding body lipid content (kg). As a simplification it is assumed that all PCB in the body of a seal is bounded in fat tissues. Hence, the total PCB content (mg) can be expressed as:

$$\bar{W}_{\text{PCB},i}^j = \bar{c}_i^j \bar{W}_i^j = \bar{c}_i^{lip,j} \bar{W}_i^{\text{lip},j} \quad (110)$$

Here, $\bar{c}_i^j$ is the total body PCB concentration (mg/kg) and $\bar{c}_i^{lip,j}$ is the PCB lipid concentration (mg/kg). From Eqs. (109) and (110), it follows that:

$$\bar{c}_i^{lip,j} = \bar{c}_i^j / \bar{\rho}_i^j \quad (111)$$

Eq. (111) is used to convert total body PCB concentrations to PCB lipid concentrations. For grey seals, most samples are collected in the autumn, at the middle of the gestation period when seals of all age classes are quite fat. As a simplification, it is assumed that mean body fat indices during sampling are the same for all age classes and remain constant throughout the gestation period; $\bar{\rho}_i^j = \hat{\rho}$. Hence, the mean PCB lipid concentration for age class $i$ at the end of year $t$ is calculated as:

$$\bar{c}_i^{\text{lip}}(t) = \bar{c}_i(t) / \hat{\rho} \quad (112)$$

Analogously, the total body PCB concentration $\bar{c}_{P,j}(t)$ of prey $j$ during year $t$ is calculated from the PCB lipid concentration $\bar{c}_{P,j}^{\text{lip}}(t)$ as:

$$\bar{c}_{P,j}(t) = \bar{\rho}_{P,j} \bar{c}_{P,j}^{\text{lip}}(t) \quad (113)$$

Here, $\bar{\rho}_{P,j}$ is the mean body lipid index for prey $j$. It was assumed that the mean body lipid index for Baltic grey seals during the gestation period is $\hat{\rho} = 30$ % for all age classes (Iverson 2002).

### 4. Toxicodynamic Model

The toxicodynamic model describes how PCB body loads of fetuses, pups and females cause cumulation of damage, hazard and stress and how these are translated into reduced survival and fertility (Fig. 8). The toxicodynamics are based on the threshold damage model (TDM), introduced by Ashauer et al. (2007). TDM is in turn based on DEBtox theory but adds cumulative effects and capacity to recover.

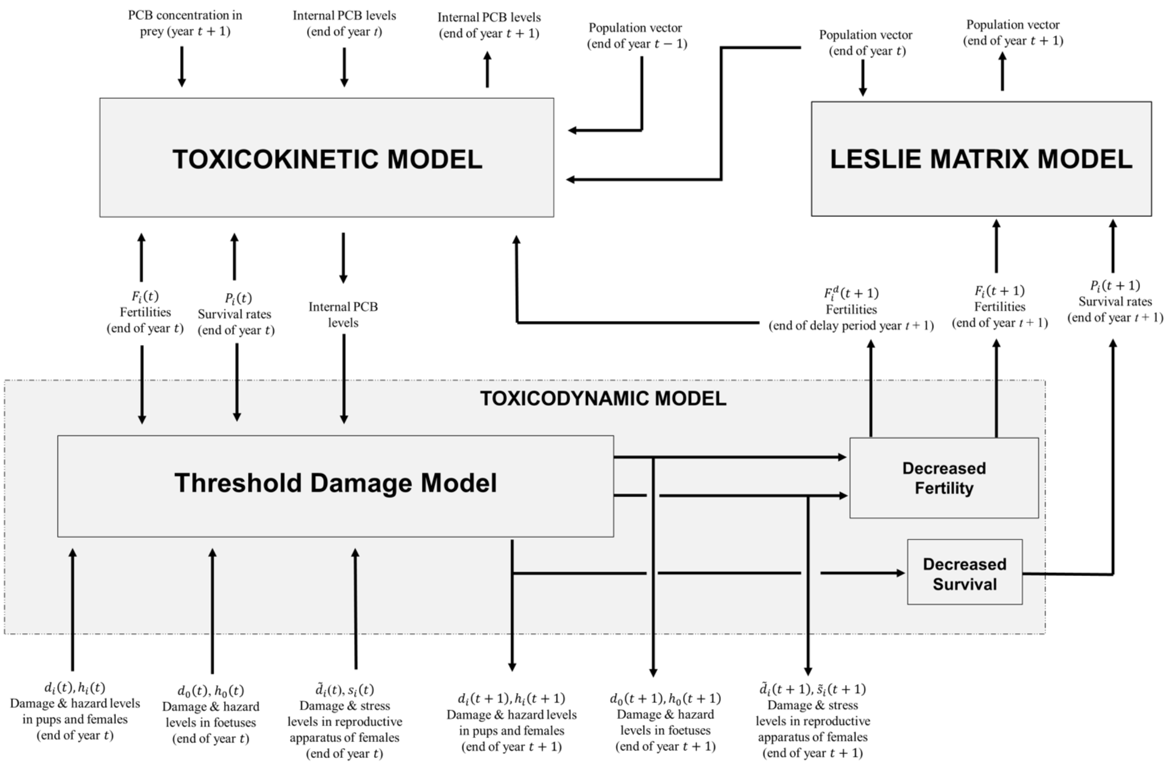

**Fig. 8** The toxicodynamic model and its connection to other sub models. Text boxes represent sub modules. Free texts are inputs and outputs, with arrows showing relations to sub modules. The flowchart illustrates computations performed for year $t + 1$, based on inputs from the previous year $t$.

DEBtox assumes several physiological modes of action (pMoA) for accumulated contaminants based on which DEB state parameters the contaminant stress is applied to, resulting in different physiological effects (e.g., growth, maturation, reproduction, etc.). Two PMoAs act directly on reproduction; *reproductions costs* (costs for *maturation* of females) and *embryonic hazard* (increased costs for *somatic maintenance* of embryos). *Reproduction costs* represent reduced efficiency in the utilization of energy for maturation of reproductive organs and offspring production, whereas *embryonic hazard* causes increased offspring mortality (an embryo dies if its somatic maintenance costs cannot be paid).

The current TD model follows the general principles of DEBtox in assuming that the toxic effects of PCBs on grey seals are considered through the lens of particular pMoAs affecting fundamental physiological processes like somatic maintenance. While our model is not based on DEB theory in the description of energetics and lifehistory traits, we interpret internal PCB concentration effects using DEBtox pMoAs using a hazard model to account for mortality of adults, pups and fetuses as well as damage to reproductive organs. In this context, we assume decreased survival can result from increased somatic maintenance costs (e.g., repair of lesions) due to cumulative damage causing hazard in pups, juveniles, and adult females. Decreased reproduction can result from increased fetal mortality (fetal hazard) and reduced offspring production (reproductive stress). Instead of adopting a costs model, where negative reproductive effects of damaged reproductive organs result from increased costs for offspring production, we adopted a hazard model and considered reduced offspring production as a result of increased costs for maintenance of the reproductive apparatus due to cumulated damage (representing reproductive organs lesions etc.). No offspring is produced if the maintenance costs of the reproductive apparatus cannot be paid. The costs model (Billoir, et al., 2007) was also explored to account for damaged reproductive organs, but found to be less consistent with empirical findings and thus not presented herein. Other indirect effects on reproduction, such as decreased feeding rate and increased growth costs are neglected. Notice that three different kinds of damage are considered: 1) damage to pups and females (causing hazard that decrease survival); 2) damage to reproductive apparatus (causing stress that reduces fertility); and 3) damage to fetuses (causing hazard that kills them and reduce the fertility of their mothers). All toxic effects and the corresponding PMoAs are summarized in Fig. 9.

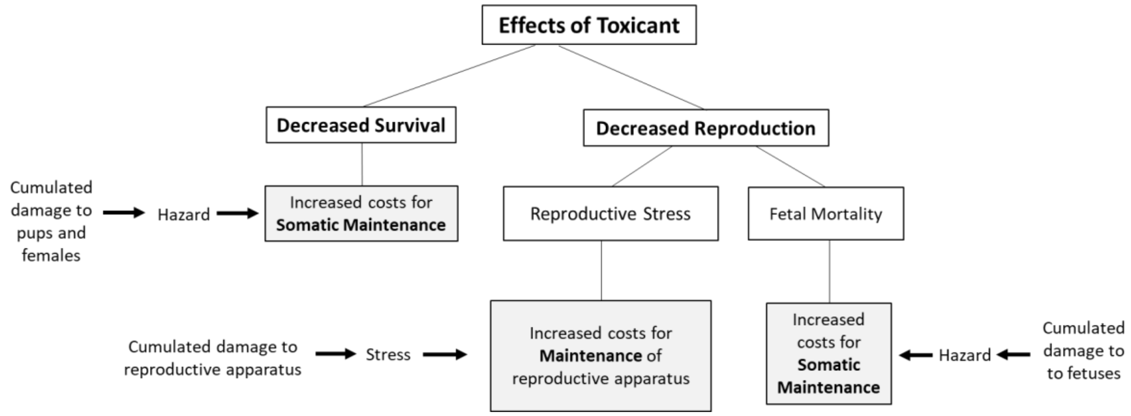

**Fig. 9** Effects of toxicant on survival and reproduction considered in the toxicodynamic model and their relation to physical processes (maintnence and maturation).

The *hazard model* (Billoir et al. 2007) accounts for *embryonic hazard* by interpreting effects on reproduction as embryo mortality during gestation. The *hazard rate* describes the instantaneous probability to die and is defined as (Jager et al. 2011):

$$\frac{\mathrm{d}h}{\mathrm{d}\tau} = -\frac{1}{P}\frac{\mathrm{d}P}{\mathrm{d}\tau} \quad (114)$$

Here, $h(\tau)$ is the hazard (the probability to die at time $\tau$) and $P(\tau)$ is probability to survive until time $\tau$. Hazard rate multiplied by a small time increment gives increase of *hazard*, the probability to die during the time interval. The survival probability (the probability of the organism to survive until time $\tau$) is obtained from Eq. (114) as:

$$P(\tau) = P^0 e^{-h(\tau)} \quad (115)$$

The *background survival* (survival probability for animals with no exposure to toxicants) is $P^0 = e^{-m_0}$, where $m_0$ is the background mortality rate (mortality rate in absence of toxicant exposure). The probability of surviving from one age class $(i - 1)$ to the next $(i)$ is given as:

$$P_i = l(i)/l(i-1) = e^{-m(i)} = P^0 e^{-h(i)} \quad (116)$$

According to the hazard model, the reduced reproduction rate at exposure time $\tau$ is:

$$R(\tau) = R^0 e^{-h(\tau)} \quad (117)$$

From the reproduction rate $R(\tau)$ at exposure time $\tau$, the number of offspring reaching age class 1 per female of age class $i$ is obtained as:

$$F_i = \int_{i-1}^{i} R(\tau)\,\mathrm{d}\tau \quad (118)$$

#### 4.1 Cumulation of Damage, Hazard and Stress

To calculate hazard and stress in the currently developed model, a TDM similar to that established by Ashauer et al. (2007) is applied. The cumulation of all three types of damage (damage to pups/females, fetuses and reproductive organs) are described by the same governing equations, which only differ in values of model parameters. According to TDM, the damage rate increases linearly with the internal PCB concentration, whereas the recovery rate is proportional the current damage:

$$\frac{\mathrm{d}d_{k,i}^{j}}{\mathrm{d}\tau} = \sigma_k c_{k,i}^{j}(\tau) - r_k d_k^{j}(\tau) \quad , \quad , \quad \begin{cases} k = 0, 1, i \quad , \quad i = 2, \dots, 46 \\ j = l, d, g \quad , \quad 0 \le \tau \le \Delta\tau_j \end{cases} \tag{119}$$

Here, $d_{k,i}^{j}(\tau)$ is the damage at time $\tau$ (during time period $j$), $c_{k,i}^{j}(\tau)$ is the internal PCB concentration (mg/kg) at time $\tau$, $\sigma_k$ is the *killing* or *stress rate constant* (kg/mg·year$^{-1}$) of age class $k$ and $r_k$ is the *recovery rate constant* (year$^{-1}$) of age class $k$. Notice that $k = 0, 1, i$ represents fetuses, pups and females, respectively, whereas $i = 2, \dots, 46$ represents different age classes of females. Damage is here a non-dimensional variable. The cumulated hazard $h(t)$ after exposure time $t$ is obtained as:

$$h(t) = [h(0) + [d(t) - d_{\mathrm{T}}]_+ - [d(0) - d_{\mathrm{T}}]_+]_+ \quad , \quad [x]_+ = \max(0, x) \tag{120}$$

The cumulated hazard at the end of the time period ($\tau = \Delta\tau_j$) is expressed according to Eq. (120) as:

$$h_{k,i}^{j}(\Delta\tau_j) = \left[h_{k,i}^{j}(0) + \left[d_{k,i}^{j}(\tau) - d_{T,k}\right]_+ - \left[d_{k,i}^{j}(0) - d_{T,k}\right]_+\right]_+ \quad , \quad \begin{cases} k = 0, 1, i \\ i = 2, \dots, 46 \\ j = l, d, g \end{cases} \tag{121}$$

Here, $h_{k,i}^{j}$ is the hazard at time $\tau$ (during time period $j$) and $d_{T,k}$ is the *damage threshold level* for age class $k$. Equations (119) and (121) are expressed in notations used for describing damage that causes decreased survival of females, pups and fetuses, but the same equations are applied to describe all types of damage, only differing in values of model parameters ($\sigma_k$, $r_k$ and $d_{T,k}$). In DEB theory, embryos and juveniles invest no energy in reproduction. In line with this, fetuses and pups are assumed to be unaffected by stress to the reproductive apparatus ($\tilde{\sigma}_0 = \tilde{\sigma}_1 = 0$), which yields $\tilde{d}_{0,i}^{g} = \tilde{d}_{1,i}^{j} = 0$ and $s_{0,i}^{g} = s_{1,i}^{gj} = 0$. Damage to reproductive organs starts to cumulate when seals enter age class 2. The assumption is probably realistic since PCBs are hormone-like substances that can be expected to mainly affect adults (with higher oestrogen levels than pups). All terms including damage or stress to fetuses or pups may be directly cancelled, when applying equations to find reproductive damage and stress.

For a general formulation of the toxicodynamic model, it is useful to summarize equations (78), (91) and (101) as one single equation, expressing the PCB concentrations in fetuses, pups or females at time $\tau$ during any time period:

$$c_{k,i}^{j}(\tau) = \hat{C}_{k,i}^{j} e^{-\hat{k}_{E,k}^{j}\tau} + \widehat{K}_{k,i}^{j} e^{-\hat{k}_{E,i}^{j}\tau} + \overline{K}_{k,i}^{j} \quad , \quad \begin{cases} k = 0, 1, i \quad , \quad i = 2, \dots, 46 \\ j = l, d, g \quad , \quad 0 \le \tau \le \Delta\tau_j \end{cases} \tag{122}$$

The following definitions were introduced:

$$\hat{C}_{k,i}^{j} = \begin{cases} (\kappa_{D,i}^{g}\, \hat{\kappa}_{P0,i} - \kappa_{P0,i})\bar{c}_{D,i} - \hat{\kappa}_{P0,i} c_{i,i}^{g}(0) & , \ k = 0 \\ c_{1,i}(0) - \hat{\kappa}_{L1,i}^{j} c_i(0) - \kappa_{D,1}^{j} \bar{c}_{D,1} & , \ k = 1 \\ c_{i,i}^{j}(0) - \kappa_{D,i}^{j} \bar{c}_{D,i} & , \ k = i \end{cases} \quad , \quad \hat{k}_{E,0}^{g} = \bar{k}_{G,0}$$

$$\widehat{K}_{k,i}^{j} = \begin{cases} \hat{\kappa}_{P0,i}\left(c_{i,i}^{g}(0) - \kappa_{D,i}^{g} \bar{c}_{D,i}\right) & , \ k = 0 \\ \hat{\kappa}_{L1,i}^{j} c_{i,i}^{j}(0) & , \ k = 1 \\ 0 & , \ k = i \end{cases} \quad , \quad \overline{K}_{k,i}^{j} = \begin{cases} \kappa_{P0,i} \bar{c}_{D,i} & , \ k = 0 \\ \kappa_{D,1}^{j} \bar{c}_{D,1} & , \ k = 1 \\ \kappa_{D,i}^{j} \bar{c}_{D,i} & , \ k = i \end{cases} \tag{123}$$

Insertion of Eq. (122) into the toxicodynamic equation for damage rate (Eq. (119)) yields the following first order ODE:

$$\frac{\mathrm{d}d_{k,i}^{j}}{\mathrm{d}\tau}+r_k d_k^j(\tau)=\sigma_k\left(\hat{C}_{k,i}^{j}e^{-\hat{k}_{E,k}^{j}\tau}+\hat{K}_{k,i}^{j}e^{-\hat{k}_{E,i}^{j}\tau}+\bar{K}_{k,i}^{j}\right) \tag{124}$$

The solution to Eq. (124) is:

$$\begin{aligned}&d_{k,i}^{j}(\tau)=\left(d_{k,i}^{j}(0)-D_{k,i}^{j}\right)e^{-r_k\tau}+E_{k,i}^{j}e^{-\hat{k}_{E,k}^{j}\tau}+F_{k,i}^{j}e^{-\hat{k}_{E,i}^{j}\tau}+B_{k,i}^{j}\\&k=0,1,i\,,\qquad i=2,\dots,46\,,\qquad j=l,d,g\,,\qquad 0\le\tau\le\Delta\tau_j\end{aligned} \tag{125}$$

Here, the following definitions were introduced:

$$B_{k,i}^{j}=\frac{\sigma_k}{r_k}\bar{K}_{k,i}^{j}\quad,\quad E_{k,i}^{j}=\frac{\sigma_k}{r_k-\hat{k}_{E,k}^{j}}\hat{C}_{k,i}^{j}\quad,\quad F_{k,i}^{j}=\frac{\sigma_k}{r_k-\hat{k}_{E,i}^{j}}\hat{K}_{k,i}^{j}\quad,\quad D_{k,i}^{j}=B_{k,i}^{j}+E_{k,i}^{j}+F_{k,i}^{j} \tag{126}$$

Equation (125) can be used to calculate the cumulated damage $d_{k,i}^{j}(\tau)$ in fetuses ($k=0$), pups ($k=1$) or females ($k=i$) at a time $\tau$ during a time period $j$ within a year when the initial damage $d_{k,i}^{j}(0)$ at the start of the period is known.

For **fetuses** ($k=0, j=g$), the initial damage and hazard are zero ($d_{0,i}^{g}(0)=0$ and $h_{0,i}^{g}(0)=0$). The damage and hazard of fetuses at the end of the gestation period ($\tau=\Delta\tau_g$) are then obtained from Eqs. (125) and (121) as:

$$d_{0,i}^{g}(\Delta\tau_g)=\Gamma_{i,i}^{0}c_{i,i}^{g}(0)+\gamma_i^0\bar{c}_{D,i}\,,\qquad i=2,\dots,46 \tag{127}$$

$$h_{0,i}^{g}(\Delta\tau_g)=\left[d_{0,i}^{g}(\Delta\tau_g)-d_{T,0}\right]_+\,,\qquad i=2,\dots,46 \tag{128}$$

The damage and hazard in fetuses (with mothers of age class $i$) at the end of year $t+1$ are obtained from Eqs. (127) and (128) as:

$$\left.\begin{aligned}\hat{d}_{0,i}(t+1)&=\Gamma_{i,i}^{0}\hat{c}_i^d(t+1)+\gamma_i^0\bar{c}_{D,i}(t+1)\\\hat{h}_{0,i}(t+1)&=\left[\hat{d}_{0,i}(t+1)-d_{T,0}\right]_+\end{aligned}\right\}\quad i=2,\dots,46 \tag{129}$$

For **pups** ($k=1$), the damage and hazard at the end of period $j$ ($\tau=\Delta\tau_j$) are obtained from Eqs. (125) and (121) as:

$$d_{1,i}^{j}(\Delta\tau_j)=Y_1^j d_{1,i}^{j}(0)+\Lambda^j c_{1,i}^{j}(0)+\hat{\Gamma}_i^j c_{i,i}^{j}(0)+\gamma_1^j\bar{c}_{D,1}\qquad(i=2,\dots,46\,,\ j=l,d,g) \tag{130}$$

$$h_{1,i}^{j}(\Delta\tau_j)=\left[h_{1,i}^{j}(0)+\left[d_{1,i}^{j}(\Delta\tau_j)-d_{T,1}\right]_+-\left[d_{1,i}^{j}(0)-d_{T,1}\right]_+\right]_+\qquad(i=2,\dots,46\,,\ j=l,d,g) \tag{131}$$

Notice that application of Eqs. (130) and (131) requires knowledge of initial PCB, damage and hazard level in pups ($c_{1,i}^{j}(0)$, $d_{1,i}^{j}(0)$, $h_{1,i}^{j}(0)$) at the start of the period. The damage and hazard in pups at the end of the three periods ($j=l,d,g$) year $t+1$ are obtained from Eqs. (130) and (131) as:

$$\begin{aligned}&\left.\begin{aligned}\hat{d}_{1,i+1}^{l}(t+1)&=Y_1^l\hat{d}_{0,i}(t)+\Lambda^l\hat{c}_{0,i}(t)+\hat{\Gamma}_{i+1,i}^{l}\bar{c}_i(t)\\\hat{h}_{1,i+1}^{l}(t+1)&=\left[\hat{h}_{0,i}(t)+\left[\hat{d}_{1,i+1}^{l}(t+1)-d_{T,1}\right]_+-\left[\hat{d}_{0,i}(t)-d_{T,1}\right]_+\right]_+\end{aligned}\right\}\quad i=1,\dots,45\\&\left.\begin{aligned}\hat{d}_{1,i}^{d}(t+1)&=Y_1^d\hat{d}_{1,i}^{l}(t+1)+\Lambda^d\hat{c}_{1,i}^{l}(t+1)+\gamma_1^d\bar{c}_{D,1}(t+1)\\\hat{h}_{1,i}^{d}(t+1)&=\left[\hat{h}_{1,i}^{l}(t+1)+\left[\hat{d}_{1,i}^{d}(t+1)-d_{T,1}\right]_+-\left[\hat{d}_{1,i}^{l}(t+1)-d_{T,1}\right]_+\right]_+\\\hat{d}_{1,i}^{g}(t+1)&=Y_1^g\hat{d}_{1,i}^{d}(t+1)+\Lambda^g\hat{c}_{1,i}^{d}(t+1)+\gamma_1^g\bar{c}_{D,1}(t+1)\\\hat{h}_{1,i}^{g}(t+1)&=\left[\hat{h}_{1,i}^{d}(t+1)+\left[\hat{d}_{1,i}^{g}(t+1)-d_{T,1}\right]_+-\left[\hat{d}_{1,i}^{d}(t+1)-d_{T,1}\right]_+\right]_+\end{aligned}\right\}\quad i=2,\dots,46\end{aligned} \tag{132}$$

Notice that the damage and hazard level in new-born pups with mothers of age class $i+1$ equal the levels in fetuses with mothers of age class $i$ at the end of previous year.

For **females** ($k=i$), the damage and hazard at the end of period $j$ ($\tau = \Delta\tau_j$) are obtained from Eqs. (125) and (121) as:

$$d_{i,i}^{j}(\Delta\tau_j) = Y_i^j d_{i,i}^j(0) + \Gamma_i^j c_{i,i}^j(0) + \gamma_i^j \bar{c}_{D,i} \qquad (i = 2, \dots, 46\,, \qquad j = l, d, g) \tag{133}$$

$$h_{i,i}^{j}(\Delta\tau_j) = \left[h_{i,i}^j(0) + \left[d_{i,i}^j(\Delta\tau_j) - d_{T,i}\right]_+ - \left[d_{i,i}^j(0) - d_{T,i}\right]_+\right]_+ \qquad (i = 2, \dots, 46\,, \qquad j = l, d, g) \tag{134}$$

Notice that application of Eqs. (133) and (134) requires knowledge of initial PCB concentration ($c_{i,i}^j(0)$), damage ($d_{i,i}^j(0)$) and hazard ($h_{i,i}^j(0)$) at the start of the period. The damage and hazard in females at the end of the three periods ($j = l, d, g$) year $t+1$ are obtained from Eqs. (133) and (134) as:

$$\left.\begin{aligned} \bar{d}_{i+1}^{l}(t+1) &= Y_{i+1,i}^{l}\bar{d}_i(t) + \Gamma_{i,i+1}^{l}(t+1)\bar{c}_i(t) \\ \bar{h}_{i+1}^{l}(t+1) &= \left[\bar{h}_i(t) + \left[\bar{d}_i^l(t+1) - d_{T,i}\right]_+ - \left[\bar{d}_i(t) - d_{T,i}\right]_+\right]_+ \end{aligned}\right\} \quad i = 1, \dots, 45$$

$$\left.\begin{aligned} \bar{d}_i^d(t+1) &= Y_{i,i}^d \bar{d}_i^l(t+1) + \Gamma_{i,i}^d \bar{c}_i^l(t+1) + \gamma_i^d \bar{c}_{D,i}(t+1) \\ \bar{h}_i^d(t+1) &= \left[\bar{h}_i^l(t+1) + \left[\bar{d}_i^d(t+1) - d_{T,i}\right]_+ - \left[\bar{d}_i^l(t+1) - d_{T,i}\right]_+\right]_+ \\ \bar{d}_i(t+1) &= Y_{i,i}^g \bar{d}_i^d(t+1) + \Gamma_{i,i}^g(t+1)\bar{c}_i^d(t+1) + \gamma_i^g(t+1)\bar{c}_{D,i}(t+1) \\ \bar{h}_i(t+1) &= \left[\bar{h}_i^d(t+1) + \left[\bar{d}_i(t+1) - d_{T,i}\right]_+ - \left[\bar{d}_i^d(t+1) - d_{T,i}\right]_+\right]_+ \end{aligned}\right\} i = 2, \dots, 46 \tag{135}$$

$$\bar{d}_1(t+1) = \sum_{i=1}^{45} \eta_{1,i+1}(t+1)\hat{d}_{1,i+1}^{g}(t+1) \quad , \qquad \bar{h}_1(t+1) = \sum_{i=1}^{45} \eta_{1,i+1}(t+1)\hat{h}_{1,i+1}^{g}(t+1)$$

Notice that the damage and hazard in females of age class $i = 1$ at the end of the year are calculated as the mean damage and hazard in pups at the end of the year, similar to how concentrations are treated in Eq. (107). These calculations are required to obtain initial damage and hazard levels for age class 2 at the start of the next year.

#### 4.2 Fertility and Survival Rates

Fertilities are affected by PCB exposure in two different ways; reduction due to reproductive stress and reduction due to fetal hazard. It is here assumed that the state of females at the end of the delay period (just before gestation starts) determines fertility reduction due to reproductive stress and that the state of fetuses at the end of the gestation period determines fertility reduction due to fetal hazard.

The hazard model, accounting for decreased pregnancy rates due to reproductive organ lesions, yields the fertility of females in age class $i$ at the end of the delay period year $t+1$ as:

$$F_i^d(t+1) = F_i^0 e^{-\hat{s}_i^d(t+1)}\,, \qquad i = 2, \dots, 46 \tag{136}$$

The fertility of females in age class $i$ at the end of year $t+1$ is obtained by multiplying the fertility at the end of the delay period with a reduction factor accounting for fetal hazard:

$$F_i(t+1) = F_i^d(t+1)e^{-\hat{h}_{0,i}(t+1)}\ , \qquad i = 2, \dots, 46 \tag{137}$$

Survival rates of pups and females are decreased by cumulated hazards $\hat{h}_i$. The survival rate for seals of age class $i$ at the end of year $t+1$ is obtained as:

$$P_i(t+1) = P_i^0 e^{-\hat{h}_i(t+1)}\ , \qquad i = 1, \dots, 46 \tag{138}$$

The ideal survival rates $P_i^0$ are survival rates for a population with maximal possible reproduction.

### 5. Leslie Matrix Model

Like Harding, et al. (2007), a full age-structured Leslie matrix model for Baltic grey seals is adopted, using annual time steps and 46 age classes. Since an insufficient number of males is unlikely to restrict growth of a seal population (Harding et al. 2007), only the size of the female population is considered. Individuals of age class $i = 1$ (with an age of 0-1 year) are here referred to as *pups*, whereas individuals of age class $i = 2, \ldots, 46$ are referred to as *females*. Unlike Harding, et al. (2007), the Leslie matrix elements are not constants. Fertilities ($F_i$) and survival rates ($P_i$) are linked to damage levels that have cumulated as a result of the history of internal PCB concentrations, which in turn depend on the history of PCB concentrations in the prey. Hence, the elements of the Leslie matrix are changing over time. It is assumed that all deaths occur at the end of a year and immediately after, all pups are born. Since only females that survive to the end of a year give birth to new pups, the first row of the Leslie matrix contains products of survival rates and fertilities, also a difference from Harding, et al. (2007). The Leslie matrix model is formulated as:

$$\mathbf{n}(t+1) = \mathbf{A}(t+1)\mathbf{n}(t) \tag{139}$$

$$\mathbf{A}(t) = \begin{bmatrix} P_1(t)F_1(t) & P_2(t)F_2(t) & \cdots & P_{45}(t)F_{45}(t) & 0 \\ P_1(t) & 0 & \cdots & 0 & 0 \\ 0 & P_2(t) & \cdots & 0 & 0 \\ \vdots & \vdots & \ddots & \vdots & \vdots \\ 0 & 0 & \cdots & P_{45}(t) & 0 \end{bmatrix} \tag{140}$$

The population vector $\mathbf{n}(t)$ includes the number of individuals in different age classes at the end of year $t$. The Leslie matrix $\mathbf{A}(t)$ includes fertilities and survival rates at the end of year $t$. Equation (139) is used to calculate the population vector at the end of a year, based on the population vector at the end of previous year. Since the population size is accounted for just after breeding, the Leslie matrix model may be characterized as a *post-breeding model*. Ideal fertilities ($F_i^0$) and survival rates ($P_i^0$), corresponding to a grey seal population with maximal possible reproduction (in absence of toxicant exposure), were obtained from Harding, et al. (2007).

PCBs concentrations, damage levels and hazard/stress levels in fetuses, pups and females at the end of the different time periods within a year are successively calculated. Fertilities and survival rates are updated based on these computations and a new population vector is calculated, including the number of individuals in different age classes at the end of a year. Outputs from calculations for one year are used as inputs to calculations for the next year. The procedure is repeated for one year at a time and may be continued for as many years as desired.

#### 5.1 Density dependence

Reduced fertilities due to density dependence are calculated in accordance with Caswell (2001) as:

$$F_i^{\text{red}}(t+1) = \frac{F_i(t+1)}{1 + \mathbf{f}_i \bullet \mathbf{n}(t)} \tag{141}$$

Here, $\mathbf{f}_i$ is the *i*th row of $\mathbf{f}$, where $f_{ij}$ is the effect of an individual in age class *j* on the fertility of an individual in age class *i*.

Reduced survival rates due to density dependence are similarly calculated as:

$$P_i^{\text{red}}(t+1) = \frac{P_i(t+1)}{1 + \mathbf{p}_i \bullet \mathbf{n}(t)} \tag{142}$$

Here, $\mathbf{p}_i$ is the *i*th row of $\mathbf{p}$, where $p_{ij}$ is the effect of an individual in age class *j* on the survival of an individual in age class *i*. To include density dependence in the TKTD population model, fertilities and survival rates in the Leslie matrix (Eq. (140)) are substituted by reduced values according to Eqs. (141) and (142). Unless otherwise stated, all performed analyses included density dependence.

#### 5.2 Stochasticity

To incorporate stochastic processes that affect populations we use normally distributed synchronized fluctuations, generated by numerical software, to fertilities and survival rates.

### 6. Model Simulations

#### 6.1 Steady state

If prey PCB concentrations are constant over time, a stable state will finally be reached where PCB, damage, hazard and stress levels are stationary. If density dependence is neglected, also fertilities and survival rates stabilize at constant values and the population grows (or declines) at constant rate. Under these conditions, biomagnification factors and stable population growth rate can be defined. The mean biomagnification factor for all age classes (based on lipid concentrations) is:

$$\overline{\mathrm{BMF}} = \lim_{t\to\infty} \frac{\bar{c}^{\mathrm{lip}}(t)}{\bar{c}_D^{\mathrm{lip}}} \qquad (143)$$

Here $\bar{c}_i^{\mathrm{lip}}(t)$ is the mean PCB lipid concentration in seals of all age classes and $\bar{c}_D^{\mathrm{lip}}$ is the mean PCB lipid concentration in the diet of all seals. The stable population growth rate $\lambda$ is calculated as the dominant eigenvalue of the Leslie matrix.

$$\lambda = \lim_{t\to\infty}[\max(|\lambda_i(t)|)] \qquad (144)$$

Here, $\lambda_i(t)$ are the eigenvalues of the Leslie matrix. Different levels of constant PCB lipid concentration in prey (0-10 mg/kg) were used as inputs to the TKTD population model and 100 years were simulated to investigate temporal changes in mean PCB lipid concentrations and population size for the Baltic grey seals.

#### 6.2 Sensitivity analysis

A *sensitivity analysis* investigates how uncertainty in input parameters causes uncertainty in population growth and can be used to identify parameters that are critical for population viability (Lacy et al. 2018). Assuming steady state conditions without density dependence (when growth rate is constant and nonzero), sensitivity analyses were performed for three toxicokinetic parameters (lactation rate constants $k_{L,i}$, placental transfer rate constants $k_{P,i}$ and metabolic transformation rate constants $k_{M,i}$) as well as for all toxicodynamic parameters (killing/stress rate constants, damage threshold levels and recovery rate constants).

A constant PCB lipid concentration in prey, generating positive stable growth rate over time, was used as input and the investigated parameters were varied one at time, whereas other parameters were held constant. The mean prey lipid concentration was set to 2 mg/kg, corresponding to Baltic levels of the early nineties. The relative value of a parameter ($p_{rel}$) is defined as the perturbed value ($p_{pert}$) divided by the value adopted in the model ($p_{mod}$):

$$p_{rel} = p_{pert}/p_{mod} \qquad (145)$$

Stable population growth rate was plotted versus perturbed relative value of investigated model parameter.

### References

Addison, R. F. B., P. F. 1977. Organochlorine residues in maternal blubber, milk, and pup blubber from grey seals (Halichoerus grypus) from Sable Island, Nova Scotia. J. Fish Res. Board Can. **34**:937–941.

Ashauer, R., A. B. Boxall, and C. D. Brown. 2007. New Ecotoxicological Model To Simulate Survival of Aquatic Invertebrates after Exposure to Fluctuating and Sequential Pulses of Pesticides. Environmental Science & Technology **41**:1480-1486. doi:10.1021/es061727b

Baas, J., and S. A. L. M. Kooijman. 2015. Sensitivity of animals to chemical compounds links to metabolic rate. Ecotoxicology **24**:657–663. doi:10.1007/s10646-014-1413-5

Berghe, M. V., L. Weijs, S. Habran, K. Das, C. Bugli, and J.-F. Rees. 2012. Selective transfer of persistent organic pollutants and their metabolites in grey seals during lactation. Environment International **46**:6–15. doi:10.1016/j.envint.2012.04.011

Bergman, A. 1999. Health condition of the Baltic grey seal (Halichoerus grypus) during two decades - Gynaecological health improvement but increased prevalence of colonic ulcers. APMIS **107**:270-282. doi:10.1111/j.1699-0463.1999.tb01554.x

Bergman, A. 2007. Pathological Changes in Seals in Swedish Waters: The Relation to Environmental Pollution - Tendencies during a 25-year Period. Swedish University of Agricultural Sciences, Uppsala, Sweden, PHD thesis.

Bignert, A., M. Olsson, W. Persson, S. Jensen, S. Zakrisson, K. Litzfn, U. Eriksson, L. Häggberg, and T. Alsberg. 1998. Temporal trends of organochlorines in Northern Europe, 1967-1995. Relation to global fractionation, leakage from sediments and international mesures. Environmental Pollution **99**:177-198. doi:10.1016/S0269-7491(97)00191-7

Billoir, E., A. R. R. Péry, and S. Charlesa. 2007. Integrating the lethal and sublethal effects of toxic compounds into the population dynamics of Daphnia magna: A combination of the DEBtox and matrix population models. Ecological Modelling **203**:204-214. doi:10.1016/j.ecolmodel.2006.11.021

Bodiguel, X., O. Maury, C. Mellon-Duval, and F. Roupsard. 2009. A dynamic and mechanistic model of PCB bioaccumulation in the European hake (Merluccius merluccius). Journal of Sea Research **62**:124–134. doi:10.1016/j.seares.2009.02.006

Boyd, I. L. 1984. The relationship between body condition and the timing of implantation in pregnant Grey seals (Halichoerus grypus). J. Zool. **203**:113–123.

Bredhult, C., B.-M. Bäcklin, A. Bignert, and M. Olovsson. 2008. Study of the relation between the incidence of uterine leiomyomas and the concentrations of PCB and DDT in Baltic gray seals. Reproductive Toxicology **25**:247–255. doi:10.1016/j.reprotox.2007.11.008

Bäcklin, B.-M., L. Eriksson, and M. Olovsson. 2003. Histology of Uterine leiomyoma and occurrence in relation to reproductive activity in the Baltic gray seal (Halichoerus grypus). Veterinary Pathology **40**:175-180. doi:10.1354/vp.40-2-175

Bäcklin, B.-M., and C. K. Moraeus, K. Isomursu, M. 2013. Pregnancy rates of the marine mammals - Particular emphasis on Baltic grey and ringed seals. HELCOM.

Castellini, M. A., and J.-A. Mellish. 2016. Marine Mammal Physiology: Requisites for Ocean Living. CRC Press, Boca Raton, US.

Caswell, H. 2001. Matrix population models: Construction, analysis, and interpretation. Sinauer Associates, Inc. Publishers, Sunderland, Massachusetts, USA.

Desforges, J. P., C. Sonne, and R. Dietz. 2017. Using energy budgets to combine ecology and toxicology in a mammalian sentinel species. Scientific Reports **7:46267**. doi:10.1038/srep46267

Fiskbasen. 1996. Näringsvärden i fisk. Fiskbasen.

Hansson, S., U. Bergström, E. Bonsdorff, T. Härkönen, N. Jepsen, L. Kautsky, K. Lundström, S.-G. Lunneryd, M. Ovegård, J. Salmi, D. Sendek, and M. Vetemaa. 2017. Competition for the fish – fish extraction from the Baltic Sea by humans, aquatic mammals, and birds. ICES Journal of Marine Science. doi:10.1093/icesjms/fsx207

Harding, K., and T. Härkönen. 1999. Development in the Baltic grey seal (Halichoerus grypus) and ringed seal (Phoca hispida) populations during the 20th century. Ambio **28**:619-627.

Harding, K., T. Härkönen, B. Helander, and O. Karlsson. 2007. Status of Baltic grey seals: Population assessment and extinction risk. NAMMCO Scientific Publications **6**:33-56. doi:10.7557/3.2720

HELCOM. 2018. Nutritional status of seals. HELCOM core indicator report. HELCOM.

Hickie, B. E., D. C. G. Muir, R. F. Addison, and P. F. Hoekstra. 2005. Development and application of bioaccumulation models to assess persistent organic pollutant temporal trends in arctic ringed seal (Phoca hispida) populations. Science of the Total Environment **351–352**:413–426. doi:10.1016/j.scitotenv.2004.12.085

Härkönen, T. 2016. Halichoerus grypus (Baltic Sea subpopulation). The IUCN Red List of Threatened Species 2016 (e.T74491261A74491289).

Iverson, S. 2002. Blubber. Pages 107–112 *in* B. G. W. W. F. Perrin, J. G. M. Thewissen, editor. Encyclopedia of Marine Mammals. Academic Press, San Diego, California, USA.

Iverson, S. J. 1993. Milk secretion in marine mammals in relation to foraging: can milk fatty acids predict diet? Symp Zool Soc Lond **66**:263-291.

Jager, T., C. Albert, T. G. Preuss, and R. Ashauer. 2011. General Unified Threshold Model of Survival - a Toxicokinetic-Toxicodynamic Framework for Ecotoxicology. Environmental Science & Technology **45**:2529–2540.

Jefferson, T. A., M. A. Webber, and R. L. Pitman. 2008. Marine mammals of the world: A comprehensive guide to their identification. Academic Press, London.

Jüssi, M., T. Härkönen, I. Jüssi, and E. Helle. 2008. Decreasing ice coverage will reduce the breeding success of Baltic grey seal (Halichoerus grypus) females. Ambio **37**:80–85. doi:10.1579/0044-7447

Kauhala, K., M. P. Ahola, and M. Kunnasranta. 2014. Decline in the pregnancy rate of Baltic grey seal females during the 2000s. Ann. Zool. Fennici **51**:313–324. doi:10.5735/086.051.0303

Klanjscek, T., R. M. Nisbet, H. Caswell, and M. G. Neubert. 2007. A model for energetics and bioaccumulation in marine mammals with applications to the right whale. Ecological Applications **17**:2233-2250. doi:10.1890/06-0426.1

Kleiber, M. 1962. The Fire of Life. John Wiley & Sons, New York.

Kooijman, B., D. Lika, and S. Augustine. 2019. Add-my-Pet.

Kooijman, S. A. L. M. 2000. Dynamic Energy and Mass Budgets in Biological Systems. Cambridge University Press, Cambridge, UK.

Lacy, R. C., P. S. Miller, and K. Traylor-Holzer. 2018. Vortex 10 - A Stochastic Simulation of the Extinction Process - User's Manual. IUCN SSC Conservation Breeding Group & Chicago Zoological Society, Chicago.

Lang, S. L. C., S. J. Iverson, and W. D. Bowen. 2011. The Influence of Reproductive Experience on Milk Energy Output and Lactation Performance in the Grey Seal (Halichoerus grypus). PLoS ONE **6**:e19487. doi:10.1371/journal.pone.0019487

Lundström, K., O. Hjerne, S.-G. Lunneryd, and O. Karlsson. 2010. Understanding the diet composition of marine mammals: grey seals (Halichoerus grypus) in the Baltic Sea. ICES Journal of Marine Science **67**:1230–1239. doi:10.1093/icesjms/fsq022

Peters, R. H. 1983. The ecological implications of body size. Cambridge University Press, Cambridge.

Roos, A. M., B.-M. V. M. Bäcklin, B. O. Helander, F. F. Rigét, and U. C. Eriksson. 2012. Improved reproductive success in otters (Lutra lutra), grey seals (Halichoerus grypus) and sea eagles (Haliaeetus albicilla) from Sweden in relation to concentrations of organochlorine contaminants. Environmental Pollution **170**:268-275. doi:10.1016/j.envpol.2012.07.017

Segerstedt, R. 2019. PCB's Bioaccumulation Levels in the Food Chain of Marine Top Predators in the Baltic Sea. University of Gothenburg, Gothenburg, Sweden, Degree project for Master of Science in Conservation Biology.

Subramanian, A. N., S. Tanabe, and R. Tatsukawa. 1988. Use of organochlorines as chemical tracers in determining some reproductive parameters in Dalli-type Dall's porpoise Phocoenoides dalli. Marine Environmental Research **25**:61–74.

Svensson, C. J., A. Eriksson, T. Harkonen, and K. C. Harding. 2010. Detecting Density Dependence in Recovering Seal Populations. Ambio.

Tverin, M., R. Esparza-Salas, A. Stromberg, P. Tang, I. Kokkonen, A. Herrero, K. Kauhala, O. Karlsson, R. Tiilikainen, T. Sinisalo, R. Kakela, and K. Lundstrom. 2019. Complementary methods assessing short and long-term prey of a marine top predator – Application to the grey seal-fishery conflict in the Baltic Sea. PLoS ONE **14**:1-26. doi:10.1371/journal.pone.0208694

von Bertalanffy, L. 1938. A quantitative theory of organic growth (inquiries on growth laws II). Human Biology **10**:181-213.
